# Premature Aging and Reduced Cancer Incidence Associated with Near-Complete Body-Wide *Myc* Inactivation

**DOI:** 10.1101/2023.01.30.526354

**Authors:** Huabo Wang, Jie Lu, Taylor Stevens, Alexander Roberts, Jordan Mandel, Raghunandan Avula, Bingwei Ma, Yijen Wu, Jinglin Wang, Clinton Van’t Land, Toren Finkel, Jerry E. Vockley, Merlin Airik, Rannar Airik, Radhika Muzumdar, Zhenwei Gong, Michel S. Torbenson, Edward V. Prochownik

**Affiliations:** Division of Hematology/Oncology, UPMC Children’s Hospital of Pittsburgh, Pittsburgh, PA, 15224, USA; The University of Pittsburgh School of Medicine, Pittsburgh, PA,15224, USA; Tongji University School of Medicine, Shanghai, China; Department of Developmental Biology, The University of Pittsburgh; Central South University, Xiangya School of Medicine, Changsha, Hunan, 410013, P.R.China; Division of Medical Genetics, UPMC Children’s Hospital of Pittsburgh, Pittsburgh, PA, 15224, USA; Division of Cardiology, The Department of Internal Medicine and the UPMC Aging Institute, Pittsburgh, Pittsburgh, PA, 15224, USA; Division of Nephrology, Children’s Hospital of Pittsburgh, Pittsburgh, PA, 15224, USA; Division of Endocrinology, UPMC Children’s Hospital of Pittsburgh, Pittsburgh, PA, 15224, USA; Division of Laboratory Medicine and Pathology, The Mayo Clinic, Rochester, MN, 55905, USA; The Department of Microbiology and Molecular Genetics, UPMC, Pittsburgh, PA, 15261, USA; The Hillman Cancer Center of UPMC, Pittsburgh, PA, 15232, USA; The Pittsburgh Liver Research Center, UPMC, Pittsburgh, PA, 15261, USA

**Keywords:** Cancer metabolism, DNA damage, mitochondria, reactive oxygen species, Mlx, telomeres, Warburg effect, TCA cycle

## Abstract

*MYC* proto-oncogene dysregulation alters metabolism, translation and other functions in ways that support tumor induction and maintenance. Although *Myc+/-* mice are healthier and longer-lived than control mice, the long-term ramifications of more complete *Myc* loss remain unknown. We now describe the chronic consequences of body-wide *Myc* inactivation initiated postnatally. “*Myc*KO” mice acquire numerous features of premature aging including altered body composition and habitus, metabolic dysfunction, hepatic steatosis and the dysregulation of numerous gene sets involved in functions that normally deteriorate with aging. Yet, *Myc*KO mice have extended life spans that correlate with a 3-4-fold lower lifetime cancer incidence. Aging tissues from normal mice and humans also down-regulate Myc and gradually deregulate many of the same Myc target gene sets that are dysregulated in *Myc*KO mice. Normal aging and its associated cancer predisposition are thus highly linked via Myc and its target genes and can be genetically separated.

## Introduction

The means by which the c-Myc oncoprotein (hereafter Myc) contributes to the pathogenesis of cancer have been amply chronicled over nearly 4 decades. ^1, 2^ Tumors often deregulate the *MYC* gene as a consequence of aberrant growth factor signaling or amplification, translocation or mutation of the gene that stabilize its encoded mRNA or protein. ^1–3^ *MYC* encodes a bHLH-ZIP transcription factor that, upon dimerizing with its bHLH-ZIP partner protein Max, binds to canonical “E box” elements in the proximal promoters of its numerous target genes and increases transcriptional initiation and read-through. ^4–7^ Negative regulation by Myc-Max heterodimers is indirect and mediated by their interaction with and suppression of positively-acting transcription factors such as Miz1 and Sp1. ^6, 8, 9^ Overall, much, if not the entirety, of the cellular transcriptome is subject to some form of Myc-mediated control and involves pathways that oversee cell cycle progression, metabolism, ribosomal biogenesis and translation, with the relative importance of these being tissue and context-dependent. ^4, 6, 10–14^ The magnitude of target gene regulation is further reliant on the amount of Myc protein, its accessibility to and affinity for various E boxes and the extent to which these sites are occupied by competing factors. ^6, 11^ Both primary and secondary roles for Myc in initiating and/or supporting transformation *in vivo* have been demonstrated in numerous tissues with the latter including the promotion of tumor angiogenesis and immune system evasion. ^15–18^ With some exceptions, continuous Myc expression is needed to maintain high rates of proliferation and/or the viability of normal and transformed cells both *in vitro* and *in vivo.* ^6, 13, 14, 19–23^

Despite our broad understanding of Myc’s participation in oncogenesis, considerably less is known about its role in normal development and tissue homeostasis. This is largely a consequence of germ-line *Myc* gene inactivation in mice being embryonic lethal at ∼e10.5 due to the aberrant development of placental, hematopoietic and vascular compartments. ^24–27^ In adult mice, body-wide Myc inhibition for up to 2 months using a dominant interfering bHLH-ZIP domain of Myc known as Omomyc causes transient and mild aplastic anemia and colonic epithelial hypoplasia while also allowing for the regression of pre-existing mutant K-Ras-driven lung tumors. ^21^ However, neither the degree to which Myc was incapacitated in different tissues nor any long-term consequences were described. Inactivating *Myc* in individual organs or specific cellular compartments within them has clearly demonstrated differential tissue-, age- and tumor-specific dependencies as well as Myc-dosage effects although these too have generally not been observed for extended periods of time. ^12, 27–30^ Additionally, because the above studies were performed in adult mice, the consequences of Myc inactivation initiated early in life on more general aspects of long-term growth and development and tissue and organ integrity remain unknown.

In contrast to the embryonic lethality of *Myc-/-* mice, *Myc+/-* mice are not only viable but age more slowly, have longer lifespans and are smaller than their *Myc+/+* counterparts. ^27, 31^ They are also healthier and more active throughout life and develop fewer age-related pathologies. ^31^ Their longer survival has been partially attributed to a lower incidence of cancer, which is the most common associated finding at the time of demise in inbred normal rodents. ^32, 33^ However, the degree to which *Myc+/-* mouse survival was influenced by slower aging rather than altered Myc levels is unclear given that aging is the strongest independent predictor of cancer incidence in both mice and humans. ^31, 32, 34, 35^ Collectively, these findings raise important questions concerning Myc’s role in maintaining normal body composition and tissue homeostasis and, together with previous studies, imply that the consequences of Myc loss are incremental in nature. ^27, 31^ The relatively small number of individual gene expression differences between *Myc+/-* and *Myc+/+* mouse tissues also suggested that residual Myc levels in the former animals are sufficient to exert normal or near-normal control over its numerous target genes, particularly those with the highest-affinity Myc binding sites and greatest physiologic relevance. ^11, 31^ Partial Myc expression might therefore forestall the emergence of more deleterious phenotypes that would only be revealed by a more thorough suppression. ^27^ Such a scenario would not be inconsistent with the general concept of “heterozygous advantage” whereby mutation or loss of a single allele confers a survival advantage whereas mutational homozygosity is disadvantageous or lethal. ^36^

We now describe the long-term consequences in mice of body-wide and near-complete elimination of *Myc* initiated at the time of weaning. This delay avoids the inevitable mortality associated with silencing *Myc* during embryogenesis. ^25, 27^ Unlike their *Myc+/-* counterparts, *Myc-/-* mice (hereafter “*Myc*KO” mice) display numerous features of premature aging yet live significantly longer than wild-type (WT) mice. This is at least partly attributable to a 4-5-fold lower life-long incidence of cancer. Transcriptional profiling in 3 tissues known to be particularly impacted by *Myc* loss and/or aging proviides evidence in the *Myc*KO cohort for widespread and early-onset dysregulation of genes with roles in mitochondrial and ribosomal structure and function, oxidative stress, aging, senescence, DNA damage recognition and repair and mRNA splicing. ^12–14, 23, 31, 37–39^ These findings are consistent with many of the properties described in livers and embryonic fibroblasts (MEFs) generated from these same animals. ^12–14, 23^ The transcriptomic changes also often closely resemble and precede by several months those that arise during normal aging. These include declines in both Myc itself and its numerous target genes that are recapitulated in aged human tissues. The long-known relationship between aging and cancer can therefore be genetically dissociated by inactivating *Myc*.

## Results

### Efficient generation of near total-body *Myc*KO mice

B6.129S6-*Myc^tm2Fwa^*/Mmjax mice, with LoxP sites flanking the second and third exons of the *Myc* locus ^13, 29^ (Figure S1A & B), were crossed with B6.129-*Gt(ROSA)26Sortm1(cre/ERT2)Tyj*/J mice, which express a Cre recombinase-estrogen receptor (CreER) fusion transgene driven by the ubiquitously-expressed ROSA26 promoter. ^40^ Progeny strains, containing either one or 2 copies of the CreER transgene, were examined to determine the extent to which CreER copy number influenced the efficiency of *Myc* locus excision initiated at the time of weaning (ca. 4 wks). A Taqman-based assay (Figure S1A-D) performed on multiple tissues from randomly selected mice 2 wks after finishing a 5 day course of daily tamoxifen administration showed *Myc* locus excision efficiency to be dependent on CreER transgene copy number in about half the tissues (Figure S1E). All subsequent studies were therefore performed with the strain carrying 2 copies of CreER that allowed for ∼75% - >95% excision in nearly all tissues (Figure S1E and F and Table S1). The offspring of matings between B6.129S6-*Myc^tm2Fwa^*/Mmjax and C57BL/6 mice were used as a source of mice with non-excised *Myc* alleles and are referred to hereafter as wild-type (WT) mice. They were similarly treated with tamoxifen and were used for all subsequent studies as age-matched control animals. Myc transcript levels tended to correlate well with the degree of *Myc* deletion (Figure S1E and Table S1). Follow-up qPCR and qRT-PCR studies indicated that *Myc* locus loss persisted beyond 30 months although in most tissues it was incomplete and in some cases became less prominent with aging (Figure S1F and Table S1). Although western blotting to detect Myc protein can be challenging due to its low-level expression in quiescent tissues, we were able to show a loss of expression in certain tissues with proliferative epithelial components such as lung, stomach and small and large intestine (Fig. S1G). Immunohistochemical staining confirmed the loss of localized Myc expression in the intestinal crypts of the latter two tissues as well (Fig. S1G).

### *Myc*KO mice prematurely acquire numerous age-related pathologies but survive longer than WT mice and have a significantly lower cancer incidence

Growth rates and body masses of WT and *Myc*KO cohorts were initially indistinguishable and remained so until ∼10 months of age when they began to diverge in both sexes. They then converged at 18-20 months when body masses peaked and began their age-related declines (Figure 1A). *Myc*KO mice showed earlier decreases in lean mass, increases in adiposity and increases in fat:lean mass ratios that explained the otherwise identical weights of younger mice. Thus, the overall body habitus of younger *Myc*KO mice tended to prematurely assume that of older WT mice. ^41^ In females, the differences became less pronounced as WT mice eventually acquired the same overall body composition at *Myc*KO mice. Although male *Myc*KO mice showed the same tendencies, the differences from WT mice persisted throughout life.

**Figure 1.**
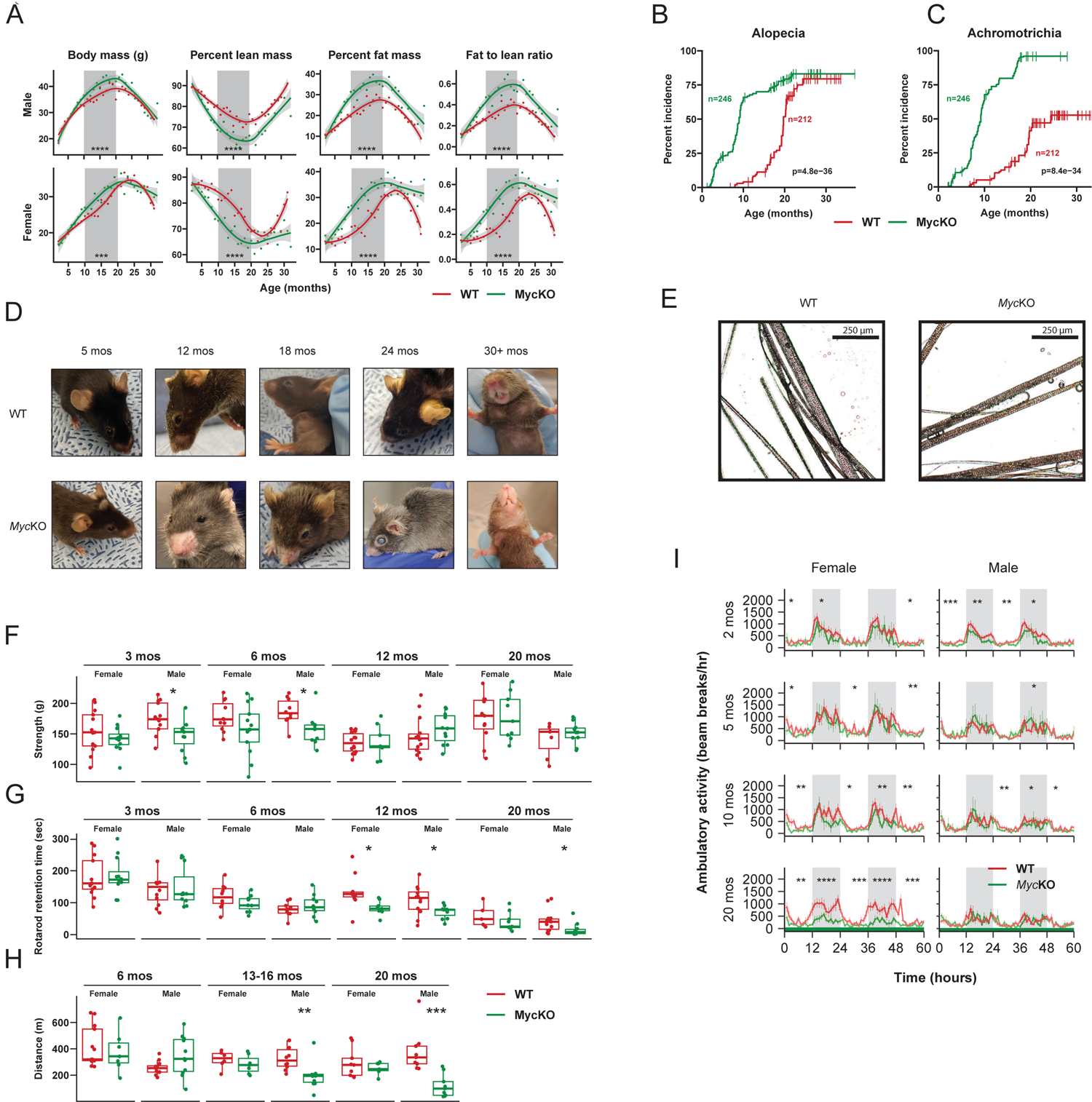
Young *Myc*KO mice display numerous aging-related phenotypes. (A). Weights and body composition of male and female WT and *Myc*KO mice. Beginning at weaning (∼4 wks of age) and continuing throughout life, WT and *Myc*KO mice were weighed weekly for the first 4-5 months and then every 2-3 wks thereafter. Results for males and females are shown separately. Percent body fat and lean mass were determined by EchoMRI at the time of each weighing. Each point represents the mean of measurements performed on 10-20 animals performed over a 2-3 day period. Times during which significant differences existed between the 2 groups are indicated by gray shading. Unpaired t test, ***=P<0.001; ****=P<0.0001. (B). Appearance of alopecia in *Myc*KO mice as a function of age. Log-rank test. (C). Appearance of achromotrichia in *Myc*KO mice as a function of age. Log-rank test. (D). Appearance of representative WT and *Myc*KO mice at the indicated ages. Note more extensive alopecia and achromotrichia in *Myc*KO mice. See Video S1 for additional examples. (E). Close up images of fur from 20 month-old WT and *Myc*KO mice showing the interspersion of dark and gray strands in the former cohort versus the greater uniformity of gray color among individual strands in the latter. (F). Four limb GripMeter testing performed at the indicated ages on male and female animals. Each mouse was tested 3 times over the course of 2-3 days with the mean result being indicated by a point. Unpaired t test, *=P<0.05, N = 9-13 for each group. (G). Rotarod testing of WT and *Myc*KO mice at the indicated ages. Each mouse was tested 3 times over the course of 2-3 days and the means are shown by a point as described in G. Unpaired t test, *=P<0.05. N = 5-14 for each group. (H). Treadmill running. Cohorts of WT and *Myc*KO mice of the indicated ages were allowed to maintain a continuous pace on an automated treadmill until becoming exhausted. Each mouse was tested 3 times over the course of 2-3 days with the mean result being indicated by a point as described in G and H. Unpaired t test, *=P<0.05. N = 6-13 for each group. (I). Diurnal activity of WT and *Myc*KO mice of the indicated ages as measured in metabolic cages over 60 hrs. N=5-10 males and 5-10 females at each age. Alternating white and gray regions of the plots denote day and night, respectively. ANOVA^147^, *=p < 0.05, **=p < 0.01, ***=p < 0.001, ****=p < 0.0001, error bars denote standard error of the mean (SEM).

*Myc*KO mice of both sexes developed alopecia and graying of their coats (achromotrichia) as early as 3-4 months of age (Figure 1B and C and Video S1). Both features tended to first appear peri-orbitally and/or peri-nasally and to then progress to the legs, neck and other areas (Figure 1D and Video S1). Achromotrichia was distinctly different between the 2 groups. In WT mice, it involved areas that were comprised of alternating dark and light gray hairs whereas in *Myc*KO mice of all ages, all hairs were uniformly light gray, occurred in patches and resembled those previously described in mice with melanocyte-specific *Myc* excisional inactivation generated during embryogenesis (Figure 1E). ^42, 43^ Some hair shafts were also comprised of alternating light-dark segments. Histologic examination of skin from areas associated with alopecia in *Myc*KO mice and identical regions from WT mice without alopecia showed the former to be associated with epidermal thickening and hyperkeratinization, loss of surface invaginations and reduced numbers of hair follicles and sebaceous glands (Figure S2A). Focal regions of peri-follicular cells that stained for senescence-associated β-galactosidase were also noted in these samples (Figure S2B).

*Myc*KO mice, particularly younger males, were generally weaker, less coordinated and less active than age-matched WT mice (Figure 1F-H). However, the magnitude of these differences, when they were first detected and their duration were influenced by age, sex and the type of test. For example, reduced grip strength was first noted in male *Myc*KO mice at 3 months of age but did not persist beyond about 10 months (Figure 1F). This occurred in parallel with the premature loss of muscle mass and its eventual equalization as WT mice aged (Figure 1A). Similarly, lessened ability to maintain balance on a Rotarod apparatus was noted in *Myc*KO mice of both sexes by 11 months and persisted in males (Figure 1G). Beginning at 13-16 months, male *Myc*KO mice also showed less endurance when subjected to a continuous running task on a treadmill (Figure 1H). ^44^ Finally, while the diurnal ambulatory activity of *Myc*KO mice was modestly reduced in younger animals, it decreased even more significantly by 20 mos. of age in *Myc*KO females (Figure 1I). Collectively, these findings indicate that *Myc*KO mice tend to acquire numerous age-related features and behaviors earlier than WT mice although at different rates. ^45–48^ In some cases, the differences persisted whereas in others they eventually converged as WT mice aged and their various deficits came to match those of their *Myc*KO counterparts.

Profound bone marrow failure underlies much of the embryonal lethality of *Myc-/-* mice. ^25, 27^ However, its severity diminishes with age such that Myc inhibition in the adult is associated with only mild and transitory marrow hypoplasia and peripheral pancytopenia ^21^. *Myc*KO mice also showed hemoglobin and leukocyte levels as low as ∼4.8-5.0 g/dl and ∼1000/μl, respectively within 10-15 days of initiating tamoxifen treatment (Figure S3A&B). The most severely affected individuals also demonstrated mild-moderate bone marrow hypoplasia that reflected the degree of peripheral cytopenias and was well-tolerated (Figure S3C). The peripheral hematologic findings resolved within several weeks and did not recur despite the long-term persistence of *Myc* loss (Figure S3A&B). ^21^ However, relative to age-matched WT mice, the bone marrow of some *Myc*KO mice tended to remain mildly hypoplastic so as to resemble that of middle-aged normal animals (Figure 3C). Inactivating *Myc* prior to weaning and/or attaining a weight of 15-16 g was associated with a near 90% incidence of severe and usually fatal pancytopenia. These findings indicated that the fetal bone marrow’s most significant period of Myc-dependency extends into the immediate post-natal period and persists for about the first month of life. ^27^

Loss of Myc in both the small and large intestine has previously been shown to be associated with a flattening of the epithelium and a loss of normal crypts that, when examined longitudinally, are of a transient nature. ^21, 28, 49^ We observed similar changes in the colonic epithelium in young *Myc*KO mice (ca. 2.5 months), that normalized by the age of 5-6 months despite the persistence of *Myc* gene loss (Figure S3D).

Nonalcoholic fatty liver disease (NAFLD) is a prominent feature of aging, particularly in the context of other age-related metabolic disorders such as dyslipidemia, obesity and insulin resistance. ^37, 50^ Neutral lipid accumulation also follows Myc loss/inhibition but is neither confined to the liver nor strictly limited to the inhibition of Myc as it has been well-documented in *Myc*KO MEFs, in cancer cell lines treated with Myc inhibitors *in vitro*, in similarly treated mice with *Mycn*-driven neuroblastomas and in the livers of mice lacking other members of the “Extended Myc Network” such as ChREBP and/or Mlx. ^14, 51–54^ Yet, the contribution of aging to Myc-dependent NAFLD development has not been thoroughly explored. ^13, 14, 29^ Although a short-term study previously showed that young WT and *Myc*KO mice accumulate increasingly more hepatic neutral lipid with age, it was particularly striking in the latter group. ^29^ Indeed, the neutral lipid and triglyceride content of 5 month-old *Myc*KO mouse livers was significantly higher than that of age-matched WT control tissue and easily rivaled that of even the oldest WT mice (Figure 2). The differences between WT and *Myc*KO livers became less pronounced with age as the excess hepatic lipid associated with latter mice was eventually matched by WT mice. The findings in *Myc*KO livers thus mimic an otherwise normal age-related process.

**Figure 2.**
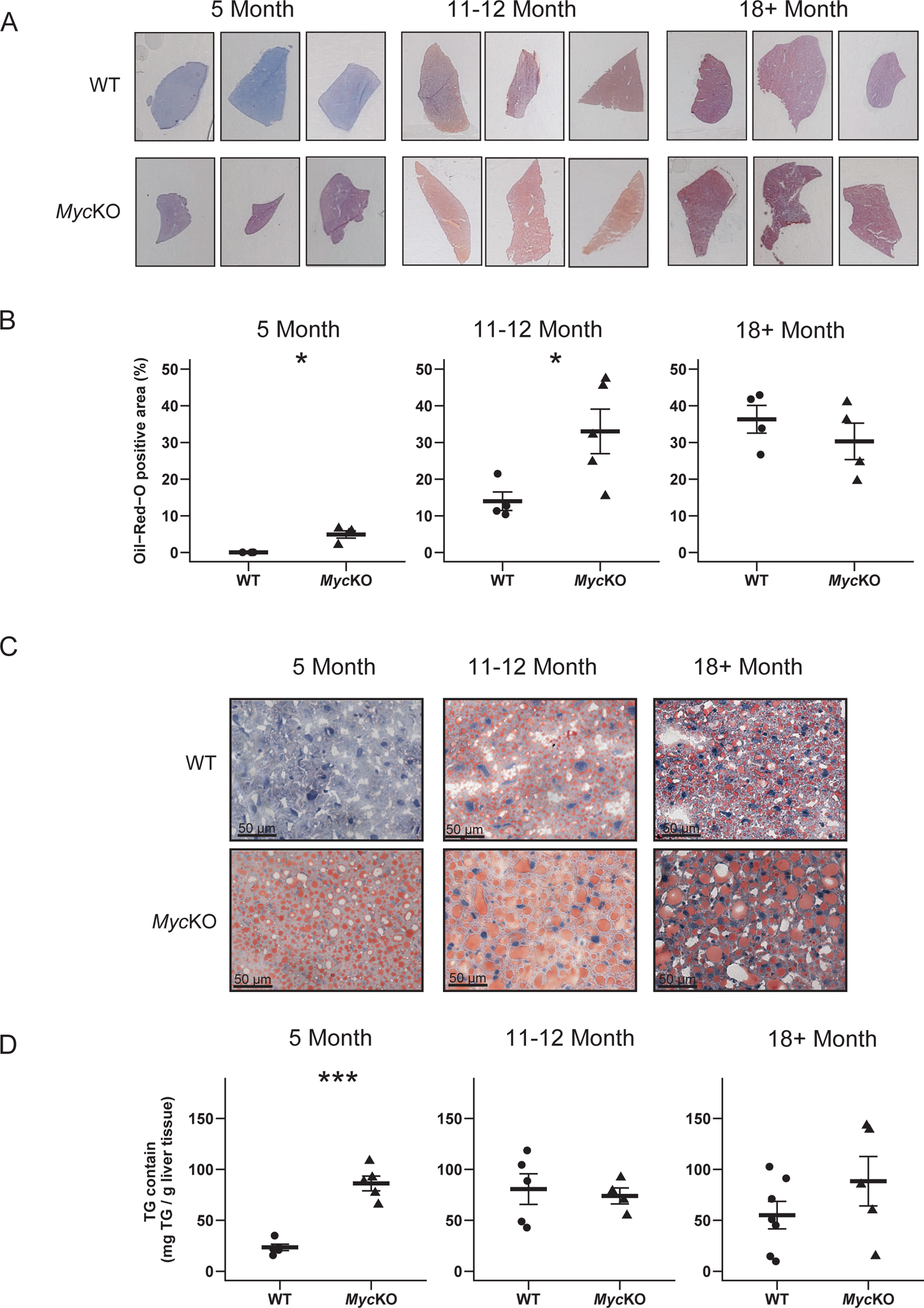
*Myc*KO prematurely develop NAFLD. (A). Low power images of representative ORO-stained liver sections of WT and *Myc*KO mice of the indicated ages. (B). Quantification of ORO-stained sections using Image J. At least 3 liver sections from each of 4-5 mice were scanned, quantified and combined. Unpaired t test, *=P<0.05, error bars denote standard deviation (SD). (C). Higher power magnification of the representative ORO-stained sections from A showing a greater prominence of large lipid droplets in *Myc*KO livers at all ages. (D). Triglyceride content of WT and *Myc*KO livers at the indicated ages. Unpaired t test, ***=p < 0.001, error bars denote standard deviation (SD).

Despite their many features of premature aging, *Myc*KO mice collectively lived significantly longer than WT mice (median survival 32.7 months vs. 28.3 months (P=1.2×10^-7^) (Figure 3A). For males the difference was 32.3 months vs. 30.2 months (p=0.037) whereas for females the difference was 33.5 months vs. 27.3 months (p=1.5 x 10^-8^). To explore the basis of this unanticipated longevity, we compiled a list of associated gross post-mortem pathologic findings (Figure 3B). Among the evaluable mice, 58.1% of WT animals had tumors, with 64.3% of these resembling B cell lymphomas, the most common cancer of in-bred mouse strains (not shown). ^33, 34^ These were frequently of high-grade appearance, were associated with massive hepatosplenomegaly and often displayed leukemic involvement due to their advanced stage. In contrast, only 17.3% of *Myc*KO mice had obvious tumors at the time of death (P<0.0001). The tumor spectrum was similar to that of WT mice and in the few cases where mice had 2 or more tumors, they were all histologically indistinguishable lymphomas (Figure 3C-H). The 3.4-fold lower cancer incidence observed in *Myc*KO mice is consistent with previous reports that many tumors rely on Myc for their initiation and maintenance. ^12–14, 19, 21, 51, 55^ Thus, the most striking aspect of the low tumor incidence of *Myc*KO mice ^31^ was that it was dissociated from a marked premature aging phenotype. It therefore seems quite likely that the prolonged survival of *Myc*KO mice can be attributed to their lower cancer incidence.

Tumor samples from 2 different organs from each of 3 *Myc*KO mice with disseminated lymphomas, likely of monoclonal origin, were examined for Myc protein expression. Control tissues included a WT normal adult liver that expressed low-undetectable levels of endogenous Myc, and a murine hepatoblastoma (HB) that expressed high levels. ^14, 56^ Myc expression in tumors from *Myc*KO mice ranged from undetectable-low to levels that were several fold higher than those in HB (Figure 3I). qPCR performed as described in Figure S1 showed that intact *Myc* gene loci were detected in each of the lymphoma samples (Figure 3J). In two cases (mouse #1 and #2), this occurred even though little to no Myc expression could be detected in the tumor. These results indicate that at least some tumors arising in *Myc*KO mice originate in rare subpopulations of cells with intact *Myc* alleles that retain the ability to be amplified in neoplastic tissues.

**Figure 3.**
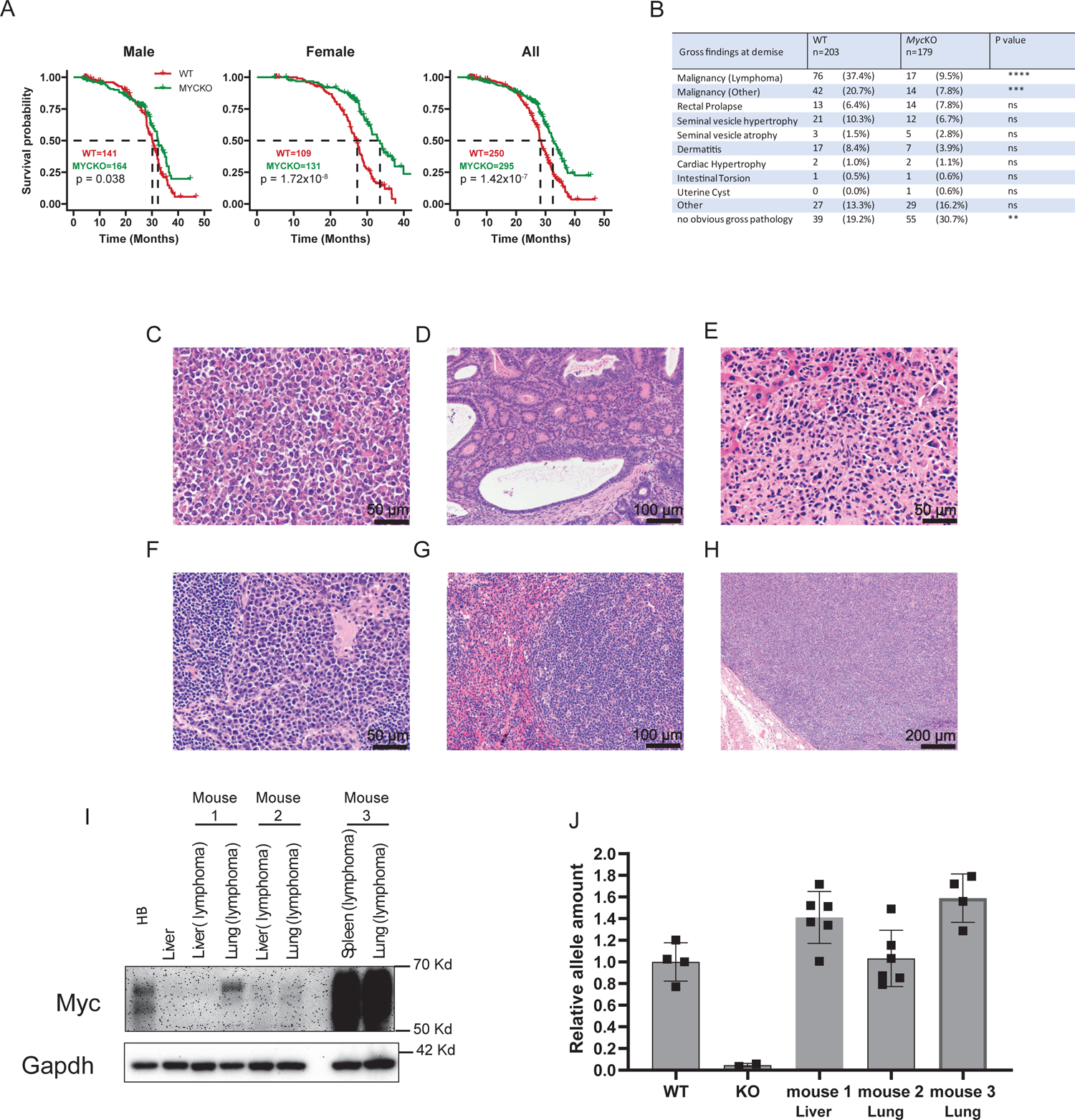
*Myc*KO mice have normal or extended lifespans and a lower incidence of tumors. (A). Natural lifespans of WT and *Myc*KO male, female and all mice depicted as Kaplan-Meier plots. Log-rank test. (B). Incidence of associated gross pathologies in WT and *Myc*KO mice at the time of demise. Specific tumor types were subsequently confirmed by histopathologic examination. Unpaired t test, **=p < 0.01, ***=p < 0.001, ****=p < 0.0001, ns: not significant. (C). A high-grade lymphoma from a *Myc*KO mouse forming a nodular mass adjacent or connected to a loop of bowel. (D). Large intestine with well-differentiated adenocarcinoma from a *Myc*KO mouse. (E). High-grade lymphoma largely replacing the normal liver parenchyma in a *Myc*KO mouse. (F). Possible plasmacytoma arising from a superficial tumor of the head and neck in a *Myc*KO mouse. (G). Lymphoma involving the spleen of a *Myc*KO mouse. (H). Lymphoma from the mouse in G effacing a lymph node adjacent to the pleural surface. (I). Immuno-blots for Myc expression. Control tissues included normal liver and a hepatoblastoma (HB) generated by mutant forms of β-catenin and yes-associated protein (YAP^S^^127^^A^) ^56^. Lymphomas from 3 *Myc*KO mice (#1-#3) were each sampled from the indicated two sites. (J). Quantification of intact *Myc* alleles in lymphomas from the 3 *Myc*KO mice depicted in panel I. DNAs were purified from several sections of each tumor and *Myc* allele quantification was performed as described for Figure S1. DNAs from WT and *Myc*KO primary MEFs (N=4 each) served as controls for 2 copies or 0 copies, respectively, of an intact *Myc* allele ^23^. Error bars denote standard deviation (SD).

### *Myc*KO mice display metabolic and mitochondrial dysfunction

A progressive deterioration of mitochondrial structure and function accompanies normal aging and senescence and notably affects organelle size, electron transport chain (ETC) activity, fatty acid β-oxidation (FAO) and redox balance. ^38, 57–62^ These defects can be independently compounded by aging-related co-morbidities such as obesity, NAFLD and insulin resistance. ^63, 64^ Conversely, mitochondrial dysfunction alone and its accompanying elevated production of reactive oxygen species (ROS) can accelerate aging and establish a positive feedback loop. ^62, 64, 65^ Reductions in mtDNA content have been noted in individuals with type II diabetes and metabolic syndrome while other studies have emphasized the coordinate down-regulation of mitochondrial proteins and abnormal ETC function in the context of NAFLD and diabetes. ^63, 66^ These findings emphasize the close co-dependencies that exist between mitochondrial dysfunction and aging. Myc’s roles in the above processes include its maintenance of mitochondrial structure and function and the mitochondrial-based oxidation of glucose, glutamine and fatty acids. ^67–73^ Linked to this is the ROS over-production that occurs in response to the ETC dysfunction that accompanies both the over- and under-expression of Myc. ^70, 72–75^

To determine how widespread *Myc* loss impacts the integrated function of some of the above pathways, we conducted longitudinal metabolic cage studies during the lifetimes of WT and *Myc*KO mice maintained on standard *ad lib* diets. The high nocturnal respiratory exchange ratios (RERs) of the youngest WT mice indicated a near complete reliance on glucose as the primary energy source (Figure 4A). RERs >1 are normally seen in juvenile mice and following post-starvation re-feeding in older mice where they signify both high levels of *de novo* fatty acid synthesis (FAS) and glucose utilization. ^76, 77^ The lower, more adult-like nocturnal RERs of *Myc*KO mice during this time of otherwise ample dietary glucose availability indicated their disproportionate reliance on fatty acid oxidation (FAO) as an energy source and/or a reduced efficiency of FAS. A switch from glycolysis to FAO following the hepatocyte-specific loss of Myc has been proposed to increase fatty acid uptake in excess of that needed for energy-generation, with the difference being stored as neutral lipid that contributes to the premature development of NAFLD (Figure 2). ^13, 14, 23, 29^ The greater reliance on FAO by *Myc*KO mice might reflect the loss of Myc’s support of glycolysis, the supply and/or transport of pyruvate for mitochondrial ATP production or the diversion of pyruvate into other pathways. ^68, 72, 78–80^ Regardless of the cause(s), the RERs of these younger *Myc*KO more closely resembled those of older animals. The differences between WT and *Myc*KO RERs persisted as mice aged but became more erratic, with the lower RERs of the latter tending to be observed during the day as well.

**Figure 4.**
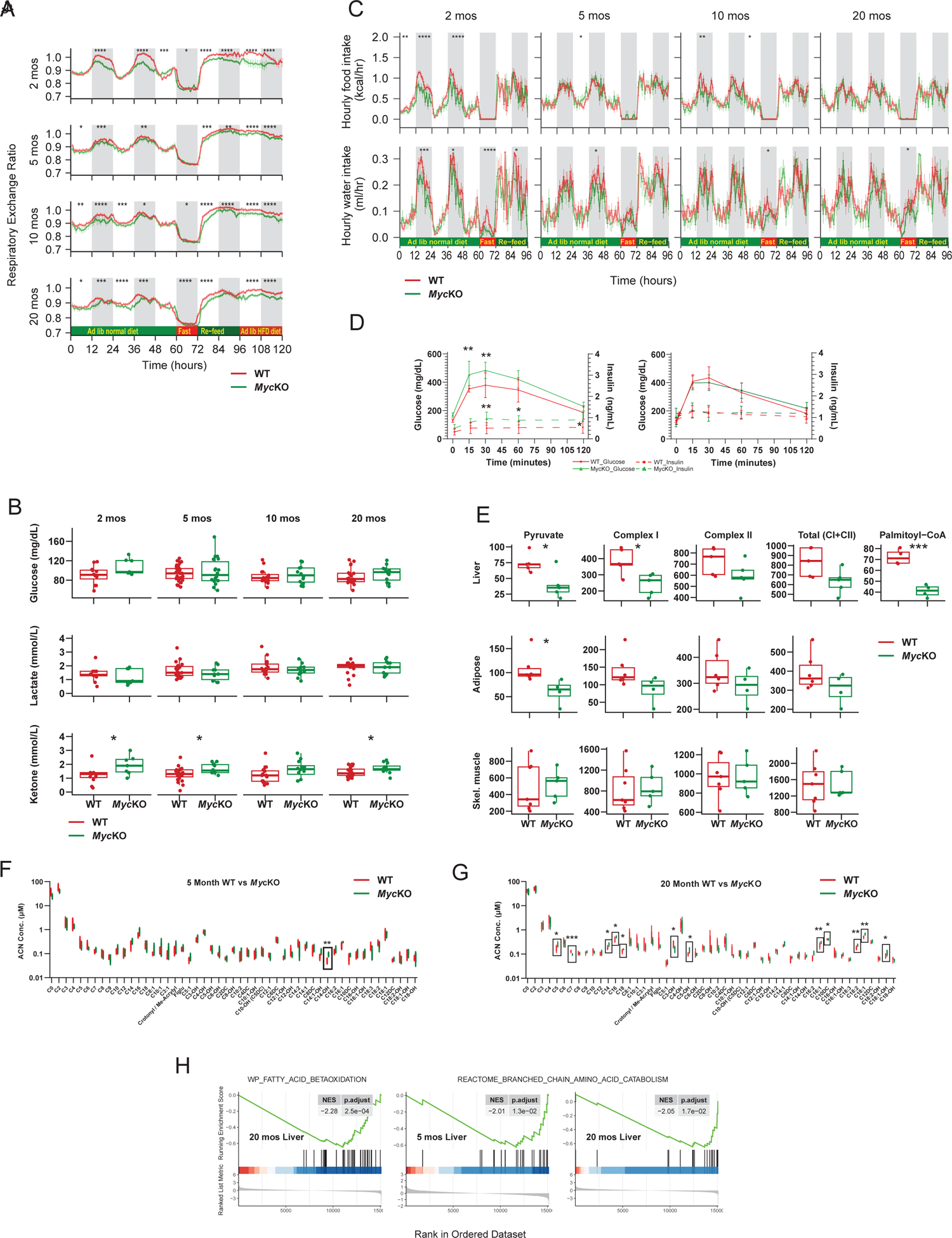
*Myc*KO mice show energy metabolism defects that are consistent with premature aging. (A). Diurnal respiratory exchange ratios (RERs). All mice were maintained in metabolic cages on *ad lib* standard diets. After 2-3 days of acclimation, RERs were calculated from the formula: RER=VCO_2_/VO_2_ ^29^. At 60 hr, mice were fasted for 12 hr and then provided with *ad* lib standard (Re-feed) or high-fat diets (HFDs) for consecutive 24 hr periods each as indicated. Each point represents the mean values obtained from N=11-13 mice/group +/- 1 S.E. ANOVA^147^, *=p < 0.05, **=p < 0.01, ***=p < 0.001, ****=p < 0.0001 (B). Fasting glucose, lactate and ketone levels on cohorts of WT and *Myc*KO mice of the indicated ages. Upon concluding the studies depicted in A, mice were starved overnight prior to obtaining these values. Unpaired t test, *=p < 0.05. (C). Summaries of hourly food and water intake measured during the course of metabolic cage studies (panel A). ANOVA^147^, *=p < 0.05, **=p < 0.01, ***=p < 0.001, ****=p < 0.0001. For actual results see Data S1. (D). Glucose tolerance tests and serum insulin levels. Mice of the indicated ages were fasted for 5 hr. and then administered a single i.p. bolus of glucose. Serum glucose and insulin levels were measured at the indicated times. N=5 mice/group. Unpaired t test, **=p < 0.01, error bars denote standard deviation (SD). (E). Oroboros respirometry results performed on partially purified mitochondria from the indicated WT and *Myc*KO tissues. Pyruvate responses were determined following the addition of malate, and ADP whereas total Complex I activity was determined following the subsequent addition to glutamate. ^12, 13, 23^ Unpaired t test, *=p < 0.05, ***=p < 0.001 (F). Top: Profiles of 51 serum acyl carnitines in 5 month-old WT and KO mice obtained after overnight fasting. Note that C14-OH (box) was the only acyl carnitine for which significant differences were observed between the 2 cohorts. N=5 mice/group, Also see Figures S4 and S5. Unpaired t test, **=p < 0.01 (G). The same 51 serum acyl carnitines were assessed in the sera of ∼20 month-old WT and KO mice as described in (F). N=5 mice/group. Boxes indicate the significant inter-group differences. Also see Figures S4 and S5. Unpaired t test, *=p < 0.05, **=p < 0.01, ***=p < 0.001 (H). GSEA for transcripts specifically involved in FAO in the livers of 20 month-old *Myc*KO mice as well as additional negative enrichment in both 5 month old and 20 month old *Myc*KO mice for genes comprising the BCAA catabolic pathway. Results were generated from RNAseq data obtained from liver, adipose and skeletal muscle of each of the indicated cohorts but were significant only in the liver as shown.

During 12 hr overnight fasts, the RERs of WT and *Myc*KO mice converged, indicating that the 2 groups could respond similarly when demands for FAO were particularly acute. However, the resumption of *ad lib* feeding with either standard or high-fat diets, was again associated with lower RERs in *Myc*KO mice, emphasizing their continued over-reliance on FAO and/or an inability to fully utilize glucose. This blunted response persisted throughout life. In keeping with their greater reliance upon FAO, *Myc*KO mice also generally displayed higher levels of fasting serum ketones in the face of normal serum glucose and lactate levels (Figure 4B). Finally, younger *Myc*KO mice showed episodic reduced levels of water and food intake (Figure 4C and Data S1). While the latter might have forced a somewhat greater reliance on FAO as an energy source in the youngest of these mice (Figure 4A) it seems unlikely that it fully explains the lower RERs, given that they were not associated with peripheral hypoglycemia (Figure 4B) and persisted with aging and post-starvation feeding when food intake equalized. Collectively, these findings, like previous ones, show that many cell and tissue types in which Myc or N-Myc are compromised have dysfunctional glycolysis and oxidative phosphorylation (Oxphos) and a disproportionate reliance on FAO. ^6, 52–54, 70, 72, 73^

Less efficient FAO commonly accompanies aging and correlates not only with increased adiposity and NAFLD but also with ketosis, hyperglycemia and insulin resistance. ^61, 81^ In contrast, the reduced glycolysis and Oxphos seen in response to Myc inhibition is accompanied by a switch to and greater reliance upon FAO that attempts to maintain a positive ATP balance. ^12, 13, 23, 52, 53, 72, 73, 82^ Thus, the lower RERs of young *Myc*KO mice might reflect an integration of these opposing factors that favors FAO. The trend for the RERs of aging WT and *Myc*KO mice to converge (Figure 4A) might also indicate a more rapid age-related decline in FAO in the latter group balanced by the persistence of the high FAO needed to maintain dysfunctional mitochondria. To examine this, we performed glucose tolerance tests (GTTs) and, in parallel, measured insulin levels in juvenile (2 mos. of age) and young adult (5 mos. of age) WT and *Myc*KO mice. Baseline fasting glucose levels were similar in the two groups (Figure 4B and D). However, in response to a glucose challenge, younger *Myc*KO mice displayed the exaggerated hyperglycemia and hyperinsulinemia that characterizes Type 2 diabetes (Figure 4D). Taken together, these findings indicated that *Myc*KO mice demonstrate defects in glucose metabolism that can explain their greater reliance on FAO. However, as these mice age and undergo metabolic compensation with regard to glucose tolerance, their continued over-reliance on FAO appears to be the result of other defects.

Partially purified mitochondria from liver, white adipose tissue and skeletal muscle of 5 month-old mice were assessed for their oxygen consumption rates (OCRs) in response to pyruvate and succinate, which serve as measures of Complex I and Complex II function, respectively. ^12–14^ Even when pyruvate availability was non-rate-limiting, the Complex I responses from *Myc*KO liver and adipose tissue mitochondria were lower than those from WT mitochondria (Figure 4E). No differences were observed in Complex II activity. In livers, where the response was sufficiently strong and reproducible, we also measured OCRs in response to palmitoyl CoA, the β-oxidation of which also donates electrons to Complex I. ^83^ OCRs from *Myc*KO mice were markedly lower, which together with the defect in pyruvate oxidation, suggested a generalized dysfunction of Complex I.

The conjugation of carnitine to long-chain fatty acids and the ensuing bi-directional active transport of these acyl carnitines across the inner and outer mitochondrial membranes are key initial steps in the FAO pathway. ^84^ Both inherited and acquired Complex I disorders are associated with elevated serum levels of 3-hydroxy-C14-carnitine (C14-OH) which originates from the inefficient oxidation of long chain carbon fatty acids. ^84^ Indeed, a mass spectrometry-based evaluation of 51 serum acylcarnitines in 5 month-old mice confirmed that, on average, C14-OH levels were 1.5-fold higher in the *Myc*KO group (Figure 4F and Figure S4A). Although this difference disappeared by 20 months of age, it was replaced by 12 new changes, most notably involving the accumulation of even longer chain (C16 and C18) serum acylcarnitines (Figure 4G and Figure S4B). This finding is consistent with a progressive deterioration of the FAO pathway in aging *Myc*KO mice and closely resembled defects that have been described in aging humans with Type 2 diabetes. ^85^ The apparent normalization of C14-OH in this older cohort likely reflected a decline in the C14 pool resulting from the accumulation of the longer chain precursors and their defective oxidation to shorter chain acylcarnitines. As mice aged, each cohort continued to accumulate distinct but overlapping aberrant serum acylcarnitine profiles (Figure S5). The above-noted indication of inefficiencies in FAO by *Myc*KO mice are consistent with the observed NAFLD, low RERs and relative insulin resistance of the youngest members of this group (Figures 2 and 4A&D). Also noted in the 20 month-old *Myc*KO cohort was an accumulation of C5-carnitine (Figure S4B). Clinically, this finding is diagnostic of errors in mitochondrial branched chain amino acid (BCAA) catabolism and implied that the energy-generating defects of the aging *Myc*KO cohort had broadened to include the catabolism of valine, leucine and isoleucine. ^86^ Supporting this idea as well as the above-described defects in FAO was an examination of RNAseq data that revealed significant negative enrichment for gene sets specifically involved in FAO in the livers of 20 month-old *Myc*KO mice as well as additional negative enrichment in both 5 and 20 month-old *Myc*KO mice for gene sets comprising the BCAA catabolic pathway (Figure 4H).

Blue native gel electrophoresis (BNGE) of ETC complexes and measurements of their *in situ* enzymatic activities showed no significant cohort- or age-related differences in their structure or function (Figure 5A&B). ^56, 73, 87^ However, tissue- and age-dependent differences between WT and *Myc*KO mice were found upon examination of a subset of important, and in some cases, rate-limiting Myc-regulated glucose transporters and enzymes that function in glycolysis and link it to the TCA cycle (Figure 5C). ^14, 72, 79, 88^ For example, the glucose transporters Glut1, which is encoded by a direct positive Myc target gene ^79^, and Glut4 ^88^ were regulated in opposite directions in 5 month-old *Myc*KO livers whereas Glut2 levels did not change. Glut4 is normally expressed at low-undetectable levels in the liver thus suggesting that, in this context, it might be a negative Myc target. ^88^ This appeared to be true in skeletal muscle of *Myc*KO mice as well where Glut4 is the major glucose transporter. ^88^ Also demonstrating differences in expression between WT and *Myc*KO 5 month-old mice was the skeletal muscle-specific isoform of phosphofructokinase (PFK-M) that contrasted with no change in the liver-specific PFK-L isoform. Finally, pyruvate dehydrogenase (PDH) activity appeared to be increased in 5 month-old *Myc*KO livers by virtue of the enzyme’s reduced inhibitory phosphorylation (pPDH) whereas in skeletal muscle, this activity was decreased. ^80^ These latter findings suggested that distinct tissue-specific increases in PDH activity might represent a response to the impaired hepatic function of mitochondrial Complex I as a way of increasing the supply of acetyl-coenzyme A and the activity of the ETC (Figure 4E-H).

**Figure 5.**
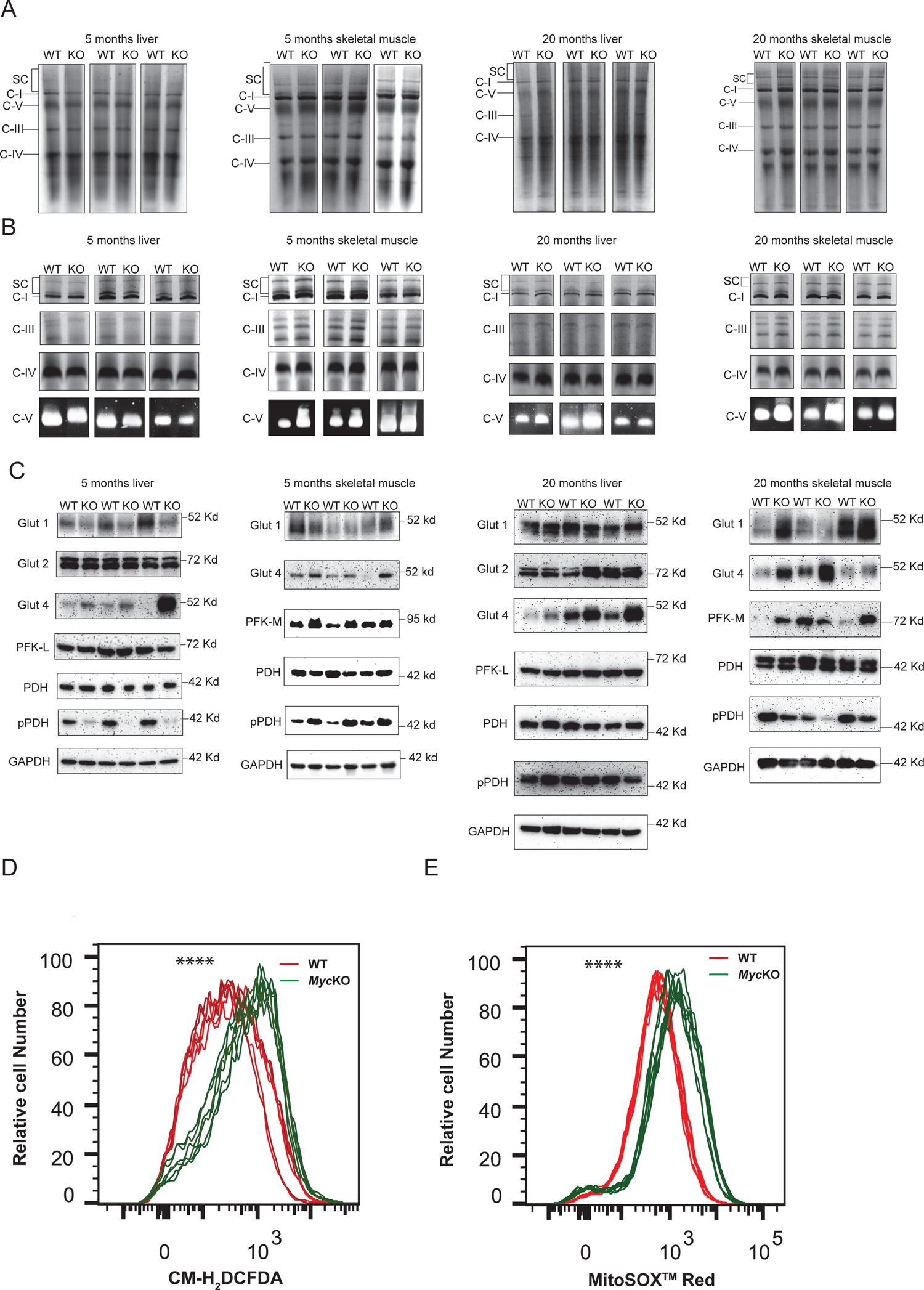
ETC structure and function and glucose handling differ between WT and *Myc*KO livers and skeletal muscle. (A). BNGE profiles of liver and skeletal muscle ETC Complexes I-IV, Complex V and supercomplexes (SCs) purified from mitochondria of 5 month old and 20 month old mice ^73^. SCs are comprised of higher order assemblies of Complexes I, III and IV. ^73^ (B). *In situ* enzymatic activity for Complexes I, III, IV and V from A. ^73^ (C). Immunoblot analyses of select key factors involved in glucose and pyruvate transport and metabolism from the tissues shown in A and B. (D). Total ROS production by MEFs isolated from WT and *Myc*KO mice as measured by the uptake and oxidation of CM-H_2_DCFDA. ^23^ N=6 biological replicas per group, all of which are depicted. (E). Mitochondrial-specific ROS production as measured by the uptake and superoxide-mediated oxidation of MitoSox^TM^-Red. N=6 biological replicas per group, all of which are depicted. Unpaired t test, ****=p < 0.0001

Similar immunoblotting performed on 20 month-old mice also revealed tissue- and cohort-specific differences while providing evidence for age-dependent changes (Figure 5C). For example, the lower levels of Glut1 originally observed in *Myc*KO livers and skeletal muscle persisted only in the latter tissue of the older mice, whereas the initially observed higher levels of Glut4 expression in *Myc*KO mice remained elevated in both sets of older tissues. Elevated skeletal muscle PFK levels also persisted. In contrast, neither PDH nor pPDH levels were significantly altered in livers whereas pPDH levels in *Myc*KO skeletal muscle were reduced as originally observed in 5 month-old livers.

ROS over-production occurs in response to both the over- and under-expression of Myc. ^23, 73, 75^ In the former case, this is presumably due to hyperactive but otherwise normal mitochondria that retain high ETC efficiency but simply generate more ROS, whereas in the latter case, it is due to ETC dysfunction and increased fractional electron “leakage” across the mitochondrial membrane. ^70, 71, 89^ To investigate this, we employed early passage WT and *Myc*KO primary MEFs from the above mice. The latter were derived by exposing WT MEFs for 10 days to 4-hydroxytamoxifen to achieve >95% excision of *Myc*, complete loss of Myc protein expression and proliferative arrest. ^23^ Despite their growth-arrested state, sub-confluent cultures of these *Myc*KO cells generated more total ROS based on staining with CM-H_2_DCFDA (Figure 5D). Parallel staining with MitoSox^TM^-Red, which detects superoxide, indicated that much if not all of the aberrantly produced ROS were of mitochondrial origin (Figure 5E). Collectively, these findings, and those depicted in Figure 4A, E, F and G and Figures S4 and S5, argue that younger *Myc*KO mice acquire some of the same mitochondrial and ETC functional defects normally observed in aged WT mice.

#### Tissue-specific gene expression differences between WT and *Myc*KO tissues correlate with observable age-related phenotypes

Comparative whole transcriptome profiling was performed on liver, mesenteric white adipose tissue and hind limb skeletal muscle from 5 month-old mice because of their known reliance on Myc for maintaining tissue homeostasis and/or because they undergo age-related phenotypic and gene expression changes (Figure 1A&B). ^31, 32, 37–39, 90^ We first verified the expected dysregulation of multiple Myc target genes in these *Myc*KO tissues using gene set enrichment analysis (GSEA) from available transcript collections in the Molecular Signatures Database (MSigDB), v2022.1. The gene expression changes were consistent with the previously documented inactivation of *Myc* (Figure S1E and Table S1), although the precise numbers and identities of the affected transcript sets and their relevant enrichment were distinct for each tissue (Figure S6A-F). ^14^

The paucity of individual gene expression differences between WT and *Myc+/-* tissues likely reflects the low basal Myc levels in WT tissues and/or the relatively modest effect of Myc haplo-insufficiency on genes with high-affinity Myc binding sites. ^11, 19, 31^ Thus, to capture the greatest possible transcriptional variation among WT and *Myc*KO tissues, to avoid computational bias and to categorize transcripts into relevant functional categories, we performed gene set enrichment analysis (GSEA) using the MSigDB and Enrichr data bases, which contain a comprehensive collection of gene sets from numerous sources. The largest, most significant and representative of these were selected and arranged using Ridgeline plots. 7 particularly noteworthy categories were immediately apparent due to the robustness and degree of dysregulation of their collective gene sets. All had been previously identified in association with aging, senescence or the conditional inactivation of Myc (Figure 6A, Figure S7 and Data S1). ^23, 57–59, 91–98^

**Figure 6.**
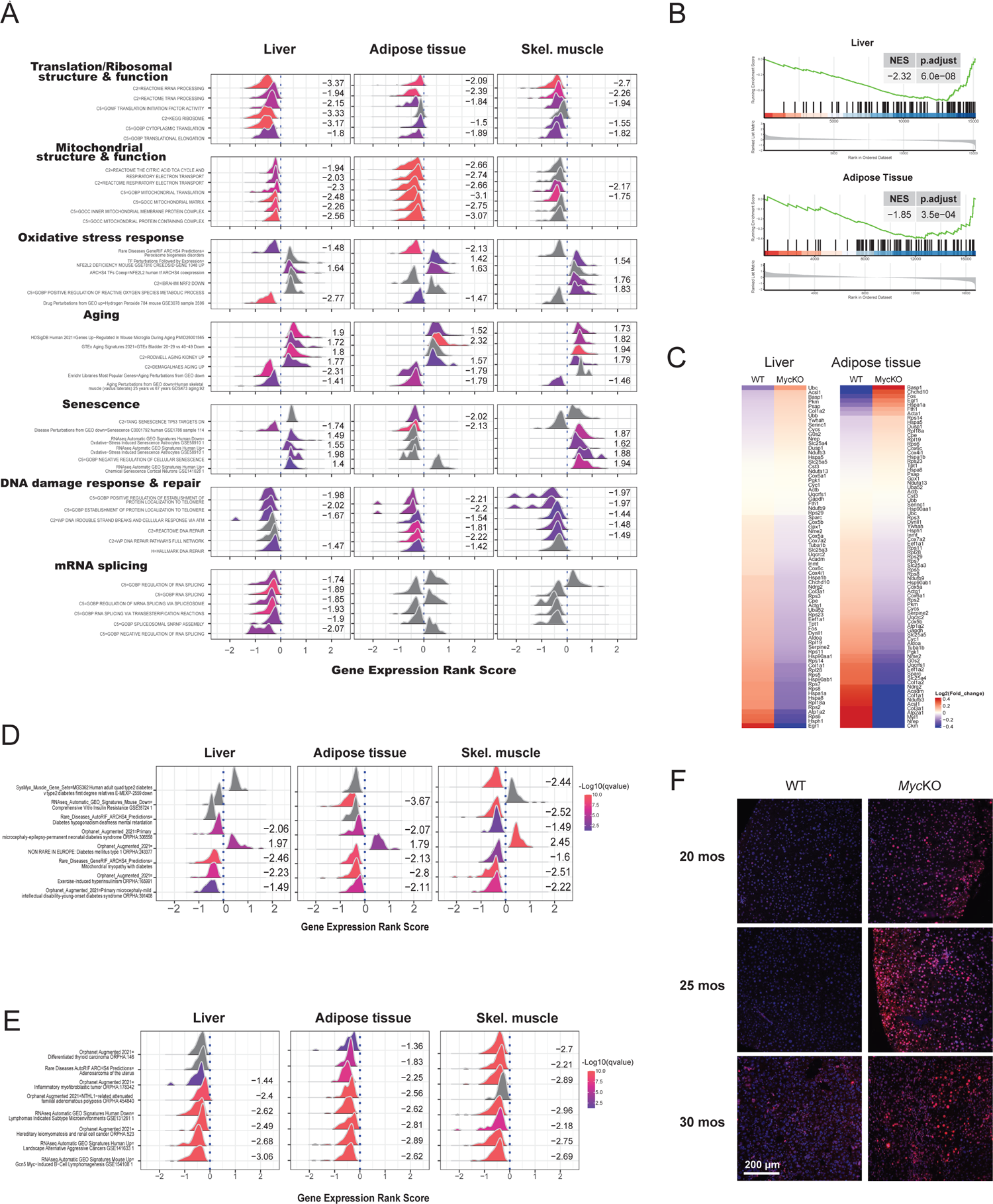
Tissues from 5 month-old *Myc*KO mice are enriched for transcripts associated with aging and senescence. (A). GSEA was performed with gene sets from the Enrichr and MsigDB data base and compared between the indicated tissues from 5 month-old WT and *Myc*KO mice. ^148–150^ Clusterprofiler was used to display representative examples of the most recurrent and prominent of the gene sets within each category using the Ridgeline plot application tool. ^151, 152^ Numbers to the right of each profile indicate its normalized enrichment score. Curves shown in gray and without any adjacent enrichment scores indicate gene sets that were not significantly enriched in that tissue but were enriched in at least one other. Values >0 along the abscissas indicate gene sets that were up-regulated in *Myc*KO tissues whereas values <0 indicate gene sets that were down-regulated. The top 7 most recurrent and highly enriched functional categories are shown. See Figure S7 for standard GSEA plots of these and other significant gene sets from each of these categories. Data S1 – Source data contains the full names of all relevant gene sets and relevant information to allow further evaluation of individual gene sets within each category. (B,C) GSEA and heat map for transcripts from the Enrichr database that correlate with aging in most tissues and across species. Results are from the livers and adipose tissues of 5 month old WT and *Myc*KO mice. See Data S1 for the complete list of these genes and their levels of expression. (D). Gene sets associated with Types 1 and 2 diabetes selectively enriched in the indicated tissues of 5 mo *Myc*KO mice. Gene sets are from the Enrichr and MSigDB data bases. See Data S1 for a complete list of these genes and their relative levels of expression. (E). Gene sets associated with cancer selectively enriched in the indicated tissues of 5 month old *Myc*KO mice. Gene sets are from the Enrichr and MSigDB data bases. See Data S1 for a complete list of these genes and their relative levels of expression. (F). Double-stranded DNA breaks in livers of MycKO mice. Typical examples of immuno-staining for γ-H2AX in the indicated mice are depicted. Shown are merged micrographs obtained from separate images for γ-H2AX immunostaining (red) and DAPI staining (blue).

The first of the above categories was designated “Translation/Ribosomal Structure & Function” given that it contained gene sets encoding ribosomal subunits, factors involved in the synthesis and processing of rRNAs and tRNAs and the regulation of translation (Figure 6A and Figure S7A). A second broad category, “Mitochondrial Structure & Function”, contained multiple gene sets encoding components of the inner and outer mitochondrial membranes, the matrix, the ETC and mitochondrial ribosomes (Figure 6A and Figure S7B). We previously identified both of these gene categories in MEFs and hepatocytes lacking *Myc* and/or members of the Extended Myc Network, namely ChREBP and/or Mlx. ^6, 12–14, 23^ The current work extends these earlier findings to livers with *Myc* depletion involving other cellular components and to 2 other *Myc*KO tissues. In these latter cases, the precise gene set enrichment profiles were again highly tissue-specific.

In keeping with the dysregulation of gene sets within the “Mitochondrial Structure and Function” category, as well as with our above-cited evidence for ETC Complex I dysfunction and aberrant ROS production (Figures 4E-G and 5D and E) was a third gene set category that we dubbed “Oxidative Stress Response” (Figure 6A and Figure S7C). Among its notable component gene sets were those associated with the potent redox-responsive transcription factor NFE2L2/NRF2, the generation of superoxide and the response to hydrogen peroxide ^87^. These findings were consistent with previously noted aberrant ROS production in other *Myc*KO cells and tissues (Figure 5D&E).^23, 29, 71^

Also, strongly dysregulated in 5 month-old *Myc*KO tissues were 2 large categories of transcripts that are normally associated with aging and senescence (Figure 6A and Figure S7D&E). A 79-member transcript subset from the Enrichr age-related gene collection and specifically selected for its near-universal association with aging in both mice and humans was also dysregulated in *Myc*KO liver and adipose tissue in ways that again marked them as possessing an “older” transcriptional profile (Figure 6B&C and Data S1). Further related to these were several gene sets previously identified as being enriched in tissues from individuals with types I and II diabetes and cancer (Figure 6D&E and Data S1). In *Myc*KO tissues, the former group of gene sets was generally dysregulated in the directions seen in diabetic tissues whereas in the latter, the directions of enrichment were opposite those seen in cancers and thus consistent with the low lifetime cancer risk of *Myc*KO mice (Figure 3B).

“DNA Damage Recognition and Repair” comprised the sixth functional category of enriched gene sets in *Myc*KO tissues (Figure 6A and Figure S7F). These genes encode multiple proteins that recognize, respond to or repair radiation-induced lesions, single- and double-stranded DNA breaks and other genotoxic damage and maintain functional telomere/shelterin complexes. Many of these gene sets are also highly dysregulated in *Myc*KO MEFs, which demonstrate abnormal responses to a broad and mechanistically diverse array of DNA damaging agents as well as high levels of double-stranded DNA breaks. ^23^ Monogenic disorders involving individual members of these gene sets include Werner syndrome, Nijmegen break syndrome, ataxia telangiectasia, Faconi’s anemia, xeroderma pigmentosum and “telomeropathies” such as dyskeratosis congenita, idiopathic pulmonary fibrosis and aplastic anemia. Most of these disorders are associated with premature aging and senescence and an exceedingly high early cancer incidence. ^96, 99–101^ In most cases, gene sets in *Myc*KO tissues were enriched in the same direction as occurs in the human condition and in *Myc*KO MEFs. ^23^ *Myc*KO livers also showed significantly higher levels of double-stranded DNA breaks as determined by immuno-staining for γ-H2AX (Fig. 6F). These findings indicate that Myc oversees an assortment of inter-connected pathways that normally participate in the tissue-specific recognition and response to DNA damage and are dysregulated in syndromes associated with premature aging and high cancer susceptibility.

Comprising the final category, and enriched only in *Myc*KO livers, were gene sets encoding components of the spliceosome, which orchestrates intron-exon junction recognition, lariat formation/removal and exon-exon ligation (Figure 6A and Figure S7G). ^102^ Despite the significant dysregulation of this category, we found no evidence for a higher incidence of the frame shifts or indels that can accumulate during aging and senescence as a result of novel splicing events.^91, 92, 103^

### Myc target gene dysregulation in *Myc*KO mice mimics that of normal aging in WT mice and humans

Approximately 10% of the transcript differences that distinguish young and old WT and *Myc+/-* mice have been attributed to direct Myc targets. ^31^ To determine whether this remained true in tissues with more complete Myc loss, we compared the tissue RNAseq profiles of both young (5 month) and old (∼20 month) WT and *Myc*KO mice. Focusing on the previously enriched gene sets (Figure 6 and Figure S7) allowed for two major observations. First, in liver and adipose tissue, more gene set differences existed between WT and KO tissues at 5 months than at ∼20 months (Figure 7A, columns 1 vs. 2 and 5 vs. 6 and Data S1). This indicated that much of the transcriptional dysregulation of young *Myc*KO tissues eventually occurred in the tissues of older WT mice, thus tending to equalize the earlier gene expression differences. Stated another way, the extensive gene set enrichment seen in young *Myc*KO mice reflects the fact that more of these sets were prematurely dysregulated so as to resemble the profiles of older animals. Indeed, many of the gene sets that distinguished the livers and adipose tissue of younger from older WT mice, were identical to those that were dysregulated following Myc loss. The premature appearance of this Myc-dependent transcript fingerprint in young mice was consistent with their numerous age-related features (Figures 1A, 2 and 4). ^32, 37, 77, 95^

**Figure 7.**
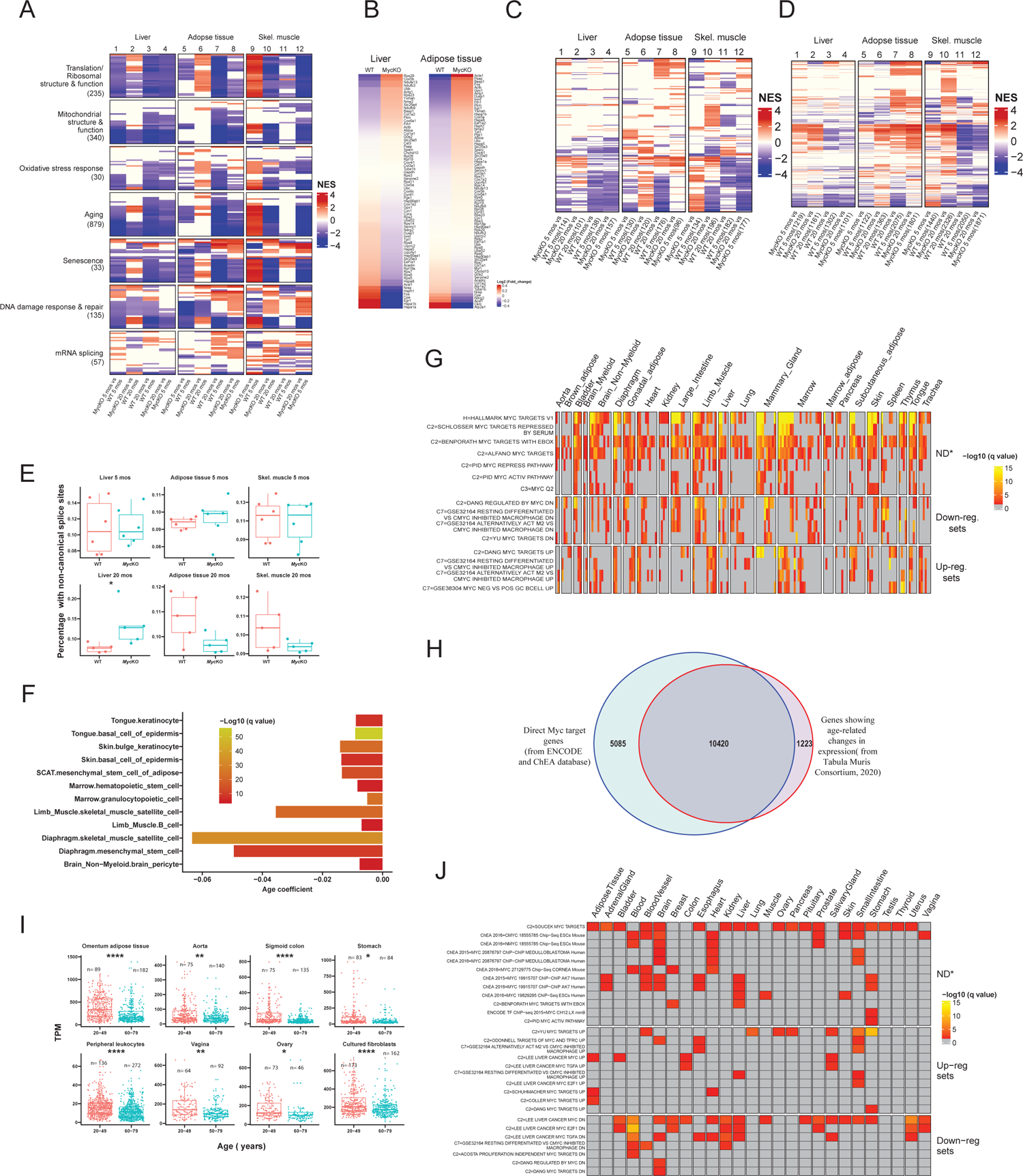
Gene expression differences in young and old mouse tissues reflect declines in Myc and Myc target genes. (A). Age- and Myc-dependent gene set enrichment differences among young (5 month) and old (20 month) WT and *Myc*KO tissues. 5 replicas of the indicated tissues were used for RNAseq analysis from each cohort of mice. GSEA again identified the 7 major categories of gene sets shown in Figure 6A. The total number of gene sets for which significant enrichment was observed is indicated beneath each category. Individual colored lines within each category represent a single gene set, the top 30 of which are shown for each category. Data S1 lists all relevant gene sets and others depicted in this panel. (B). Heat map for the 79 transcripts shown in Figure 6 B&C that correlate with aging in most tissues examined and across species. Data S1 lists all relevant gene sets depicted in this panel. (C). Heat map for the expression of individual gene sets related to Type I and Type 2 diabetes, including those depicted in Figure 6D in the indicated tissues from 5 month-old and 20 month-old WT and *Myc*KO mice. Data S1 lists all relevant gene sets depicted in this panel. (D). Heat map for the expression of individual gene sets related to cancer, including those depicted in Figure 6E in the indicated tissues from 5 month-old and 20 month-old WT and *Myc*KO mice. Data S1 lists all relevant gene sets depicted in this panel. (E). Transcriptome-wide quantification of non-canonically-spliced transcripts in the indicated tissues. ^151–154^ Unpaired t test, *=p < 0.05. (F). Significant declines in Myc transcript levels in 12 of 90 single cell populations derived from 23 individual young (1-3 months) and old (18-30 months) mouse tissues. ^104^ The results are expressed as q values based upon correlation coefficients that compared transcript levels across aging populations. (G). Over-representation analysis of Myc target gene sets analyzed using the above-cited single cell RNAseq data. ^104^ 58 Myc target gene sets from the MSigDB data base were screened to determine if their component transcripts were over- or under-represented in young versus old mice. Gene sets for which significant dysregulation was observed in at least 40 of the 90 single cell populations are shown, although 74 of the cell populations (82.2%) showed enriched representation of at least one gene set (Data S1). “Down-regulated sets”: those that are canonically down-regulated in response to Myc over-expression, “Up-regulated sets”: those that are up-regulated in response to Myc over-expression, “ND”: sets comprised of both positive and negative targets whose overall direction of response could not be determined. (H). Overlap between direct Myc target genes and those which undergo significant age-related changes in expression (q<0.05). Direct Myc targets ^105, 106^ were obtained from the Enrichr data base. Gene expression differences were compiled by comparing transcript levels of 76 single cell populations derived from 23 tissues from 1-3 month-old and 18-30 month-old mice. ^104^ (I). Differences in Myc transcript levels in tissues obtained from young and old human tissues. All results were obtained with RNA samples isolated directly from the indicated tissues except in the case of primary fibroblasts, which were transiently expanded *in vitro*. Results are from the Broad Institute’s GTEx data base. Unpaired t test, *=p < 0.05, **=p < 0.01, ***=p < 0.001, ****=p < 0.0001 (J). Enrichment of Myc target gene sets (see Figure S6A-C) in aging and senescent human tissues and cell lines.

The second observation was made in skeletal muscle where young WT and *Myc*KO mice again showed dysregulation of many of the same Myc-dependent gene sets previously documented in livers and adipose tissue (Figure 7A, column 9). However, a number of these, notably in the “Aging” category, were enriched in directions opposite of those seen in livers and adipose tissue. Also seen was a more pronounced persistence of gene set enrichment in WT and *Myc*KO skeletal muscle at 20 months of age than had been seen in liver and adipose tissue (column 10 versus columns 2 and 6). This was particularly so for gene sets contained within with “Translation /Ribosomal structure & function” and “Mitochondrial structure & function” categories. This suggested that gene set enrichment differences in skeletal muscle were less well equalized in response to aging and that many of the Myc-dependent gene sets with these categories continued to show additional dysregulation beyond what could be attributed to aging alone. Indeed, the overall marked directional change of the enrichment of some gene sets in skeletal muscle from down in young *Myc*KO mice to up old mice, which was seen to a much lesser degree in both liver and adipose tissues, appeared to be due to an overall aging-related decline in WT mice (Figure 7A, column 11).

The previously mentioned 79-member transcript set that is highly associated with aging in mice and humans (Figure 6B&C and Data S1) was re-examined in older livers and adipose tissues where the differences between WT and *Myc*KO was again noted, despite the differences being fewer and less pronounced than they had been in 5 month-old tissues (Figure 7B and Data S1). This again suggested that the aging-related gene signatures associated with younger *Myc*KO tissues were eventually partially reproduced by 20 months of age.

A more comprehensive assessment of Types 1 and 2 diabetes-associated gene sets than used previously (Figure 6D) was performed on young and old tissues from the WT and *Myc*KO cohorts. Extensive tissue-specific dysregulation of these sets was again observed when tissues from young mice were compared (Figure 7C columns 1, 5 and 9 and Data S1). This dysregulation persisted in older mice although the numbers and identities of the gene sets and the degree to which they were enriched changed in tissue-specific ways (Figure 7C columns 2,6 and 10). These findings indicated that the gene set dysregulation associated with insulin deficiency or resistance was already quite extensive in the tissues of *Myc*KO mice and remained so throughout life. Similar analyses performed with a larger number of gene sets associated with and/or deregulated in cancer (Figure 6E) also showed enrichment in *Myc*KO tissues regardless of age (Figure 7D and Data S1).

As noted above (Figure 6A and Figure S7G), an “RNA splicing” category of gene sets was particularly enriched in the livers of young *Myc*KO mice but was not associated with any increases in non-canonical mRNA splicing. Only 8 such gene sets remained enriched in the livers of 20 month-old *Myc*KO mice, again indicating that the dysregulation in younger mice represented an acceleration of the changes that eventually occurred in WT mice (Table S2). However, in these older *Myc*KO livers, a significant increase of non-canonically spliced transcripts was now observed (Figure 7E). Thus, while significant dysregulation of genes encoding components of the splicing apparatus occurred in some young *Myc*KO tissues, the functional consequences of this, as manifested by increases in aberrantly spliced transcripts, occurred only in older livers and thus likely relied upon additional age-dependent and Myc-independent functions as well as tissue context.

RNAseq results from a previous comparative study of young (1-3 months) and old (18-30 months) mice ^104^ showed highly significant age-related declines in Myc transcript levels in 12 of 90 (13.3%) single cell populations isolated from 23 tissues (Figure 7F and Data S1). 35 of 58 Myc target gene sets (60.3%) from the MSigDB data base were also dysregulated in one or more single cell populations of most of these 23 tissues (Figure 7G and Data S1). In the vast majority of cases for which the directionality of dysregulation could be determined, it correlated with the age-related declines in Myc levels in that positive Myc target gene sets were down-regulated and negative Myc target gene sets were up-regulated. These results thus documented more extensive age-related alterations of direct Myc target gene transcript collections than would have been anticipated based solely on changes in Myc expression alone.

The afore-mentioned single cell RNAseq data were next combined and used to search the ENCODE and ChEA data bases to determine how many of the proximal promoters of the genes encoding these transcripts are directly bound by Myc. ^105, 106^ 89.5% of genes whose expression changed significantly during normal aging were found to be direct Myc targets and 67.2% of ChIP-seq-confirmed direct Myc target genes from ENCODE and ChEA significantly altered their expression during aging (Figure 7H).

Myc expression also declines during the prolonged *in vitro* propagation of primary human fibroblasts in which the accompanying senescence can be either delayed/eliminated or hastened in response to the respective enforced over-expression or inhibition of Myc. ^23, 107^ In light of these findings, and those above indicating that Myc expression in normal mouse tissues diminishes with age and correlates with the expression of numerous Myc target genes, we queried the Broad Institute’s GTEx data base, which contains RNAseq results from a large number of young and aged normal human tissues (ca. 20-40 and 60-80 years of age respectively) where we found declines in *Myc* transcript levels to be common, albeit variable (Figure 7I). The most significant of these occurred in adipose tissue, sigmoid colon and peripheral leukocytes with less dramatic but still significant changes in aorta, stomach, vagina and ovary. *Myc* transcript levels were also lower in *in vitro-*propagated fibroblasts from older individuals and reflected the above-noted findings in murine fibroblasts. ^107, 108^ Consistent with Myc expression being needed to maintain the replication of normal primary MEFs and most other cell types, a previous study of >650 primary human fibroblast lines has shown that those derived from older individuals undergo replicative senescence sooner than those derived from younger individuals. ^22, 23, 27, 52, 109^

The coordinated loss of Myc expression in aging and senescent human cells and tissues documented above implied that Myc target genes should respond accordingly. Interrogating the above samples with the previously mentioned collection of direct Myc target gene sets from the MSigDB data base (Figure S6A-C) indeed confirmed that positively-regulated Myc target gene sets tended to be negatively enriched in older tissues and negatively-regulated gene sets tended to be positively enriched (Figure 7J). Collectively, these findings indicated that, in both mice and humans, normal aging and senescence is often accompanied by the down-regulation of Myc family members in parallel with the expected expression changes in Myc target gene expression. The deliberate inactivation of Myc and the ensuing dysregulation of Myc-dependent genes in young *Myc*KO mice thus mimics what otherwise occurs with normal aging and hastens the onset of age-related phenotypes and their associated biochemical and molecular defects.

## Discussion

Previous attempts to understand Myc’s role in normal development have been hampered by the embryonic lethality associated with the gene’s homozygous inactivation. ^25, 27^ By postponing Myc inactivation until the time of weaning, we avoided the factors that contribute most strongly to prenatal demise such as placental insufficiency, vascular agenesis and hematopoietic catastrophe. ^24, 26, 27^ The compatibility of this delay with survival allowed us to assess the consequences of *Myc* loss in the context of complex whole body and inter-dependent phenotypes such as body weight and composition, strength, metabolism, longevity and cancer incidence. While differing from the method previously used to generate *Myc+/-* mice ^31^, our approach nonetheless allowed for comparisons of the 2 mouse strains over their lifetimes, which has not been previously possible.

*Myc*KO mice presented 2 disadvantages. First, they did not permit an evaluation of *Myc*’s roles in the substantial growth and development that occur immediately after birth since its inactivation prior to 4 wks of age or before attaining a weight of 15-16 g was associated with >80% mortality and severe bone marrow failure (Figure S3A-C) (H. Wang, not shown). ^21^ Second, the relative prominence of the observed *Myc*KO phenotypes may be skewed so as to favor tissues with the highest and most persistent levels of *Myc* loss and/or where its influence extends into adulthood. *Myc* gene deletion in excess of 95% was routinely achieved in liver, adipose tissue, skeletal muscle and stomach, particularly in younger mice, whereas less efficient excision occurred in the brain and heart (Figure S1E&F and Table S1). The eventual partial reappearance of intact *Myc* alleles in some tissues of some mice suggested that stem cell populations with incomplete *Myc* excision and a proliferative advantage contributed to this recovery (Figure S1F and Table S1). Similarly, the few tumors that arose in aging *Myc*KO mice appeared to originate from a minority population of cells that were susceptible to transformation due to *Myc* retention (Figure 3I and J). Non-mutually exclusive reasons for *Myc*’s excisional variability include the degree to which tamoxifen penetrates different tissues and enters cells, its rate and efficiency of conversion to its active form (4-hydroxytamoxifen) by cytochrome P450 enzymes and differential accessibility of the *Myc* locus to CreER. ^110^ Despite these limitations, post-natal *Myc* inactivation provided a precisely timed and integrated assessment of the life-long consequences of its global loss on health and fitness. Our studies with *Myc*KO mice also demonstrate that some of the pathologies and phenotypes observed in models that previously relied on tissue-specific and/or pre-natal *Myc* inactivation can be largely replicated when the inactivation is delayed and includes additional cellular compartments of those tissues. ^14, 27, 29, 43^ Certain previously described phenotypes of *Myc+/-* mice which we did not observe, such as an overall reduced body size, likely have their origins much earlier in development. ^27, 31^

As mentioned above, *Myc* inactivation in adult mice and the juveniles described here was associated with generally mild bone marrow aplasia, peripheral cytopenias, colonic epithelial flattening and villous atrophy (Figure S3). ^21, 24–28, 111^ These features tended to resolve although the bone marrow of some mice remained hypoplastic and otherwise resembled that of much older normal mice (Fig. S3C). The normal weights of young *Myc*KO mice and the absence of steatorrhea provided additional direct and physiologically-based evidence that any malabsorption attributable to intestinal, hepatic or pancreatic insufficiencies is not consequential enough to impair growth (Figure 1A and not shown). These observations imply that Myc performs functions of variable importance at different stages of development and that the most deleterious consequences of embryonic *Myc* loss are mitigated when its inactivation is delayed. ^6^ It is also possible that the incomplete *Myc* excision seen in some tissues contributes to mitigating what would otherwise be more severe phenotypes (Figure S1E and Table S1).

In marked contrast to *Myc+/-* mice, even relatively young *Myc*KO mice displayed numerous and often progressive phenotypes that are normally associated with aging. Appearing at different times and manifesting different severities and natural histories, these were in some cases also influenced by gender. In addition to the bone marrow hypoplasia, they include increased fat:lean mass ratios; alopecia and achromotrichia and reduced strength, endurance and balance (Figure 1). Notable histologic and biochemical findings include NAFLD, the preferential use of fatty acids as an energy source, mitochondrial dysfunction and glucose intolerance (Figures 1, 2 and 5D). ^47, 57, 62, 63, 65^ We have previously described similar mitochondrial abnormalities and advanced steatosis in association with hepatocyte-specific *Myc* loss of relatively short duration. ^13, 14, 29^ The current findings confirm and extend these earlier ones by showing that the maximal lipid content associated with the oldest WT mice is achieved significantly earlier in *Myc*KO mice (Figure 2). Similarly, age-related changes in skin (Figure S2) recapitulate in less pronounced ways some of the previously reported consequences of embryonic melanocyte-specific *Myc* inactivation. ^43^ Many, if not all, of the abnormal phenotypes of young *Myc*KO mice are thus explainable by a combination of factors that include both the direct consequence of Myc inactivation and the secondary ones associated with normal aging (Figure 8).

**Figure 8.**
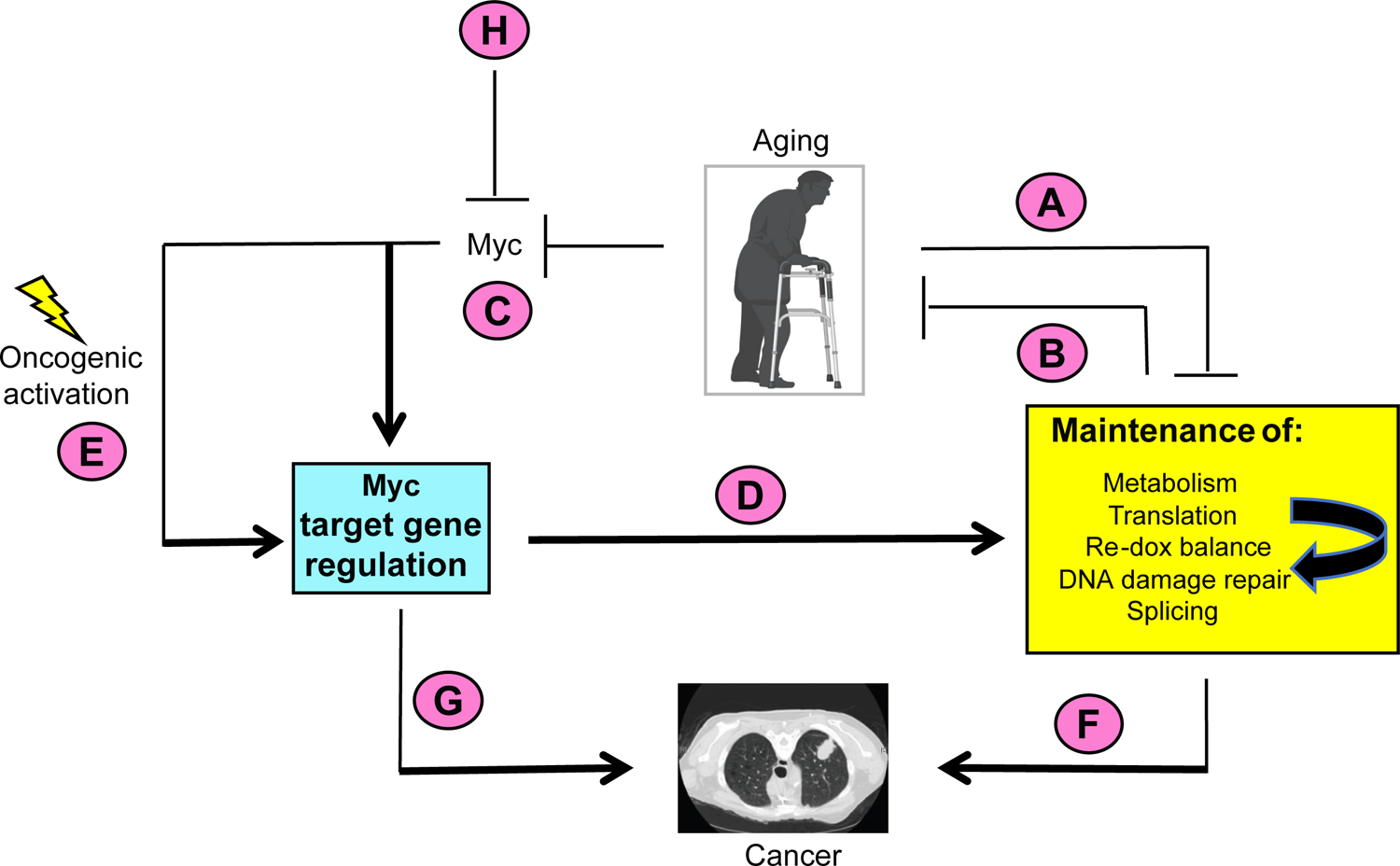
Model depicting the cooperation between normal aging and Myc regulation. (A). Normal aging and senescence are associated with the accumulation of Myc-independent defects involving the maintenance of mitochondrial and ribosomal structure and function, redox balance, DNA damage recognition and repair and splicing (yellow box) (Figure 7A). ^38, 57, 58, 62, 91– 93, 97, 98, 100, 103, 124, 128, 129^ Individual functional defects therein can negatively impact other functions (yellow box: curved arrow). For example, defective mitochondrial function can generate ROS, which in turn can cause oxidative DNA damage and impair translation. ^64, 75, 89, 126, 133^ (B). Defects in the above functions and pathways conversely accelerate aging. ^58–60, 65, 100, 127–129^ (C). Normal aging is also associated with declines in Myc levels, causing dysregulation of Myc target gene expression (blue box) (Figure 7F-I). (D). Normal levels of Myc and its target genes are needed to maintain the baseline cellular functions depicted in the yellow box. ^4, 19, 23^ These cooperate with the age-dependent and Myc-independent pathways shown in A. (E). Oncogenic activation of Myc can dysregulate Myc target genes, thus altering the functions depicted in the yellow box in ways that sustain maximal rates of tumor growth. ^4, 6, 19, 23, 55^ (F). Dysregulation of the functions described for E and D support the initiation, maintenance and/or evolution of cancer. ^4, 6, 19, 55^ (G). Other Myc target genes not shown in the yellow box, such as those pertaining to cell cycle and survival, can independently contribute to the development of cancer when they are dysregulated as a result of Myc over-expression. ^6, 15, 68, 69^ Some of these genes may be so-called “pathological targets” that contain low-affinity Myc binding sites and are only activated in response to the excessive levels of Myc associated with certain tumors. ^6, 11^ (H). *Myc*KO mice no longer regulate their target genes. As a result, they lose the ability to maintain the functions depicted in the yellow box, thereby hastening the onset of aging via the Myc-dependent pathway that support these functions (B, C and D). Reductions in Myc also eliminate a major age-dependent oncogenic pathway (E), thus leading to a reduced lifetime incidence of cancer (F and G) that contributes to the increased longevity of these mice.

While the extended lifespans of *Myc+/-* mice were originally attributed to a lower cancer incidence, it was also possible that they were due to their relative youthfulness. ^31^ Aging and cancer therefore remained temporally linked. Neoplasms are among the most frequent findings in normal mice at the time of death and age is the strongest independent predictor of cancer development in both mice and humans. ^32, 34, 35^ This closely maintained association is exaggerated in human disorders of premature aging and the animal models that mimic them despite the chronological youthfulness of the affected individual. ^94, 97, 100, 112, 113^ Underscoring this relationship is the critical contribution of Myc to the pathogenesis or maintenance of most cancers. ^2, 6, 13–15^ The finding that *Myc*KO mice have a 3.4-fold lower lifetime incidence of cancer than WT mice (Figure 3B) indicates that the usual strict association between this disease and aging can be genetically separated and is maintained by a single gene, namely *Myc*. Indeed, its elimination reduced the lifetime cancer incidence and extended lifespans by up to ∼20% (Fig. 3A). The reduced cancer incidence and increased longevity of *Myc*KO mice are even more remarkable given that they manifest other comorbidities that normally present independent risk factors for cancer development and shortened lifespans including increased adiposity, glucose intolerance, insulin resistance and multiple DNA repair defects. ^94, 100, 114–117^ Future studies may reveal factors other than a reduced cancer incidence that contribute to the extended lifespans of *Myc*KO mice.

Given Myc’s well-known contributions to tumor growth, the genetic dissociation of aging and cancer in *Myc*KO mice raised the question of how the few evaluable tumors that did arise (Figure 3B) were initiated and/or maintained. ^11, 13, 14, 55^ The expression of variable levels of Myc protein by these tumors suggested that at least some of them originated from a minority population of cells with retained *Myc* alleles (Figure 3I&J). Whether tumorigenesis in *Myc*KO mice is reduced due to a lower initiation rate or a slower growth rate (Figure 3B) will require further investigation since a role for Myc in both steps has been demonstrated, even within the same tissue. ^13, 14, 19, 23, 55^

Aside from Myc’s direct roles in tumorigenesis ^8, 12–14, 22, 55, 67, 118^ other less direct mechanisms may contribute to the low cancer incidence in *Myc*KO mice. These include increased competition for Myc target gene binding sites by transcriptionally suppressive Max-Mxd family heterodimers or by the Mlx Network, which co-regulates many Myc target genes. ^6, 119^ Mlx itself, which is the Max-like equivalent of this Network, is a recently described tumor suppressor. ^14^ Another way by which tumor growth in *Myc*KO mice might be suppressed involves higher baseline rates of neo-antigen generation resulting from DNA damage response/repair and mRNA splicing defects that could improve immune surveillance (Figures 6A, 7A&E and Figure S7F&G). ^23, 120–122^ This might be further enhanced in tumors whose growth rates were slowed by Myc loss simply by providing longer periods for neoantigens to be generated and for immune responses to mature. ^12–14^ On the other hand, the degree to which such anti-tumor immune responses are enhanced might be limited given that T cell proliferation and expansion are highly Myc-dependent. ^123^

The relationship between *Myc*, aging and cancer-related demise likely cannot be explained by any single mechanism since many of the multiple gene sets under Myc’s control functionally converge upon the so-called “Hallmarks” of both aging and cancer (Figures 6A, 7 & 8). ^38, 62, 93, 95, 98, 124–127^ Both normal aging and Myc loss are associated with excessive ROS production due to progressive ETC dysfunction and/or increased reliance on FAO (Figures 4A and 8). ^64, 89, 128^ Excessive ROS and impaired ribosomal biogenesis and translation can themselves accelerate aging under some circumstances ^98^. Both nuclear and mitochondrial DNA damage, aberrant mRNA splicing and senescence are also increased to variable degrees in the face of both aging and Myc loss. ^91, 92, 103, 129–132^ In addition to their genotoxicity, ROS also tend to inhibit translation, thus underscoring how the loss of integrity of individual Myc- and/or age-linked functions can affect one another (Figure 8). ^133^

Given the compelling premature aging phenotypes and gene expression profiles of *Myc*KO mice (Figures 6 and 7, Figure S7 and Data S1), it is reasonable to ask whether they more faithfully mimic normal aging than do other models, which are largely based upon rare monogenic disorders of DNA damage recognition or repair. ^134, 135^ While these replicate many aspects of normal aging, their molecular underpinnings do not, thus explaining their phenotypic differences, even within the same tissues. In contrast, the numerous gene set category differences between *Myc*KO and WT tissues were in many cases identified as being Myc-specific subsets of a larger group that distinguishes young and old WT mice (Figures 6 and 7). Additionally, the enrichment of these subsets correlated with normal age-associated reductions in Myc transcripts and the altered expression of Myc target genes in both mice and humans (Figure 7F-J). Together, these findings indicate that *Myc* inactivation in juvenile mice rapidly recreates its normal age-related declines, the deregulation of its downstream target genes and the deterioration of their collective functions with similar molecular and phenotypic consequences (Figures 7F-J and 8).

In our aforementioned analyses of normal murine and human tissues and single cell populations, the aging-related enrichment of Myc target gene sets tended to involve more tissues than did the decline in Myc levels. There are at least 3 non-mutually exclusive explanations for this finding. First, in some tissues, *Myc* paralogs might play a larger role in regulating these gene sets than Myc itself. For example, although none of the above cells or tissues expressed significant levels of *Mycn,* some expressed *Mycl* at levels equal to or exceeding that of *Myc*; in some cases these levels declined during aging when *Myc* itself did not (not shown). Myc target gene sets may therefore be preferentially responsive to *Mycl* in certain tissues. Second, the regulation of Myc target gene sets may be selectively sensitive to one or more Mxd proteins. ^6, 11^ Finally, competing members of the Mlx Network might displace Myc-Max complexes in some tissues, thus perhaps providing less transcriptional impact than does Myc. ^6, 11^

The term “heterozygous advantage” has been applied in reference to genes such as those encoding the α and β globins and the cystic fibrosis transmembrane conductance regulato.r ^36^ In these examples, possession of a single mutant or null allele is advantageous in that it confers relative resistance to malaria and diarrheal diseases, respectively, whereas mutational homozygosity is associated with potentially lethal disease states, i.e. thalassemia and sickle cell anemia in the former case and cystic fibrosis in the latter. ^136–138^ Similarly, hemizygosity of the triose phosphate isomerase gene confers lifelong resistance to oxidative stress whereas the null state is lethal. ^139^ *Myc* hemizygosity’s association with a spectrum of health benefits compared to the age-related pathologies of *Myc*KO mice described here are consistent with *Myc* being a somewhat different example of heterozygous advantage. ^31^ Given that *Myc* is not normally associated with germ-line mutations or copy number variations, however, this occurs in its purest form only in the context of the experimental conditions described in the current work. ^11, 25, 27, 31^ Nonetheless, single nucleotide polymorphisms within a region far upstream of the *Myc* coding region can exert significant influence over its expression and correlate with various cancer susceptibilities. ^140, 141^ Normal age-related declines in *Myc* and the ensuing target gene dysregulation might thus have very different long-term consequences depending upon genetically pre-determined levels of Myc expression. The heterozygous advantage of Myc might thus relate more to the genetic constraints upon its normal expression rather than mutation and become increasingly consequential as its levels decline with age and reach pathologic thresholds at different times.

The pharmacologic inhibition of Myc, with the intent of providing a broadly applicable cancer therapeutic, has proved to be elusive for both theoretical and practical reasons. ^142^ While attaining this goal has at times appeared to progress asymptotically, recent advances have provided cause for optimism. ^143–145^ The unexpected finding that *Myc+/-* mice displayed increased longevity, a lower overall cancer incidence and additional health dividends provided even greater reason to pursue this objective. ^31^ However, the current work indicates that caution may be warranted in pursuing the use of Myc inhibitors, particularly if the primary intention is to increase lifespan in otherwise healthy individuals. ^142^ Within these contexts our work raises a number questions that will need to be addressed before such inhibitors can be employed clinically, particularly in children with cancer where even short-term treatment with traditional chemotherapeutic agents can accelerate some features of aging. ^146^ Chief among these questions is the degree to which Myc inhibition will unintentionally accelerate aging, whether certain aging-associated phenotypes will be differentially manifested and whether some phenotypes are subject to “rejuvenation” when Myc expression is restored. A related question is the extent to which such phenotypes will still appear when Myc inactivation is delayed until later in life. Finally, might young age represent a contraindication when cancer therapy demands that Myc inhibition be both complete and/or prolonged? More refined evaluation in appropriate animal models and clinical settings will likely be necessary before answers to such questions are forthcoming.

### Limitations of the study

While the loss of Myc ultimately underlies all the phenotypes described in this report, there remain several unanswered questions. For example, to what extent do the different pathways that are impacted by *Myc*’s loss cooperate to promote premature aging and to what degree are they tissue-dependent (Fig. 8)? The alterations in mitochondrial and ribosomal structure and function, energy metabolism and DNA damage recognition/repair that distinguish *Myc*KO and WT mice have all been described previously and assigned varying degrees of importance concerning their roles in both normal and premature aging. The approach we utilized to inactivate *Myc*, based on the total body, post-natal activation of CreER is also not 100% efficient (Figure S1E and F and Table S1). Thus, it is not clear whether a more complete and widespread deletion of *Myc* remains compatible with the extended longevity we have observed and whether it would allow for a full cataloging of all potential phenotypes. Even low-levels of residual Myc expression in certain tissues may be sufficient to slow or prevent the emergence of additional features in much the same way as happens in *Myc+/-* mice. Thus, the phenotypes described here may be incomplete and/or milder than those that are potentially achievable. Contributing to this is the fact that some tissues that were not carefully examined in this study may nonetheless possess significant structural and/or functional abnormalities. Finally, it remains to be seen how delaying *Myc* inactivation until later in life will impact the age-related findings we have reported.

## Acknowledgements

This work was supported by NIH grant RO1 CA174713, a Hyundai Hope on Wheels Scholar grant, a Rally Foundation Independent Investigator Grant #22IN42 and by The UPMC Children’s Hospital of Pittsburgh Foundation (all to E.V.P). GEV was supported by NIH grant DK RO1 109907. MT was supported by NIH grant P50 CA 210964. Analysis of RNAseq data was partly supported by The University of Pittsburgh Center for Research Computing. Acylcarnitine profiling was performed in collaboration with the Rangos Metabolic Core LC/MS/MS services at the Department of Pediatrics, University of Pittsburgh Medical Center.

## Author contributions

E.V.P. conceived the study. H.W., Y.W., T.F., J.E.V., R. Ai, R.M. and E.V.P. designed experiments. H.W., J.L., T.S., A.R., J.M., B.M., R.Av, Y.W., J.W., C.V.L., M.A., R.Ai., E.V.P and Z.G. performed experiments and/or analyzed or interpreted data. H.W, J.M. and R.A. analyzed transcriptomic data. H.W. and E.V.P. interpreted transcriptomic analyses. M.S.T. performed histo-pathologic analyses; H.W., J.L. and T.S. generated figures; E.V.P. and H.W. wrote the paper. All authors read and approved the final version of the manuscript prior to submission for publication.

## Inclusion and Diversity

We support inclusive, diverse, and equitable conduct of research.

## STAR★Methods

### Key resources table

| REAGENT or RESOURCE | SOURCE | IDENTIFIER |
| --- | --- | --- |
| Antibodies |  |  |
| Glut 1 | Abcam | Cat#Ab115730 |
| Glut 2 | Proteintech | Cat#20436-1-AP |
| Glut 4 | Cell signaling | Cat#2213 |
| GAPDH | Sigma | Cat#G8795 |
| γH2A.X | Abcam | Cat#ab81299 |
| PDH | CST | Cat#3205 |
| c-Myc | Cell signaling | Cat#13987 |
| c-Myc | SantaCruz | Cat#N262 |
| p-PDH | Cal biochem | Cat#AP1062 |
| PFK-L | Avivva sys bio | Cat#855570 |
| PFK-M | R+D systems | Cat#MAB7687 |
| Rabbit IgG | Cell signaling | Cat#7074 |
| Mouse IgG | Cell signaling | Cat#7076 |
| Chemicals, peptides, and recombinant proteins |  |  |
| ABC-HRP Kit | Vector Laboratories | Cat#PK-6101 |
| 0.25% trypsin-EDTA | Corning | Cat#25-053-CI |
| DMEM | Cytiva | Cat#SH30022.01 |
| Fetal Bovine Serum (FBS) | Biowest | Cat# S1620 |
| 4-hydroxytamoxifen (4-OHT) | Sigma-Aldrich | Cat#H7904 |
| Tamoxifen | Sigma-Aldrich | Cat#T10540-29-1 |
| D-(+)-Glucose solution | Sigma-Aldrich | Cat#G8769 |
| MitoSOX™ Red | Thermo Fisher Scientific | Cat#M36008 |
| CM-H<sub>2</sub>DCFDA | Thermo Fisher Scientific | Cat#D399 |
| Critical commercial assays |  |  |
| Ultra Sensitive Mouse Insulin ELISA Kit | Crystal Chem | Cat#90080 |
| Senescence Detection Kit | Abcam | Cat#ab65351 |
| Deposited data |  |  |
| Raw data of RNAseq | This paper | GEO: GSE223676 |
| Tabula Muris Consortium mouse single cell RNAseq data | Tabula Muris Consortium | https://figshare.com/ndownloader/files/27856758 |
| RNAseq gene TPMs from young and old human tissues (GTEx Analysis V8 release: RNAseq gene TPMs by tissue) | GTEx | https://gtexportal.org/home/datasets |
| Experimental models: Cell lines |  |  |
| MEFs | In this paper | Wang et al <sup>23</sup> |
| Experimental models: Organisms/strains |  |  |
| Mouse: B6.129S6-Myctm2Fwa/Mmjax | JACKSON LABORATORY | Strain # 032046-JAX |
| Mouse: B6.129-Gt(ROSA)26Sortm1(cre/ERT2)Tyj/J | JACKSON LABORATORY | Strain # 008463 |
| Mouse: C57BL/6 | JACKSON LABORATORY | Strain # 000664 |
| Oligonucleotides |  |  |
| Primers and probes for qRT-PCR | In this paper | Figure S1D |
| **Software and algorithms** |  |  |
| ImageJ- Fiji | NIH | https://fiji.sc/# |
| FlowJo v10 | Becton-Dickinson Biosciences | https://www.flowjo.com/solutions/flowjo/ |
| R studio | Posit | https://posit.co/download/rstudio-desktop/ |
| GraphPad Prism v9.0 | GraphPad Software Inc. | https://www.graphpad.com |
| Clusterprofiler | Wu et al <sup>151</sup> | https://bioconductor.org/packages/release/bioc/html/clusterProfiler.html |
| ComplexHeatmap | Gu et al <sup>155</sup> | https://bioconductor.org/packages/release/bioc/html/ComplexHeatmap.html |
| ggplot2 | CRAN | https://ggplot2.tidyverse.org/index.html |
| survminer | CRAN | https://cran.r-project.org/web/packages/survminer/index.html |
| CLC Genomic Workbench version 21 | QIAGEN | https://digitalinsights.qiagen.com/ |
| nf-core/rnaseq-3.4 | Ewels<sup>154</sup> | https://github.com/nf-core/rnaseq |
| Other |  |  |
| DNeasy Blood & Tissue Kit | QIAGEN | Cat#69504 |
| RNeasy kit | QIAGEN | Cat#74004 |
| QIAzol Lysis Reagent | QIAGEN | Cat# 79306 |
| RNeasy Lipid Tissue Mini Kit | QIAGEN | Cat#74804 |
| SuperScript™ IV First-Strand Synthesis System | Thermo Fisher Scientific | Cat#18091050 |
| 3-12% Native PAGE Novex Bis-Tris gel | Thermo Fisher Scientific | Cat#BN1003BOX |

| REAGENT or RESOURCE | SOURCE | IDENTIFIER |
| --- | --- | --- |
| **Antibodies** |  |  |
| Glut 1 | Abcam | Cat#Ab115730 |
| Glut 2 | Proteintech | Cat#20436-1-AP |
| Glut 4 | Cell signaling | Cat#2213 |
| GAPDH | Sigma | Cat#G8795 |
| γH2A.X | Abcam | Cat#ab81299 |
| PDH | CST | Cat#3205 |
| c-Myc | Cell signaling | Cat#13987 |
| c-Myc | SantaCruz | Cat#N262 |
| p-PDH | Cal biochem | Cat#AP1062 |
| PFK-L | Avivva sys bio | Cat#855570 |
| PFK-M | R+D systems | Cat#MAB7687 |
| Rabbit IgG | Cell signaling | Cat#7074 |
| Mouse IgG | Cell signaling | Cat#7076 |
| **Chemicals, peptides, and recombinant proteins** |  |  |
| ABC-HRP Kit | Vector Laboratories | Cat#PK-6101 |
| 0.25% trypsin-EDTA | Corning | Cat#25-053-CI |
| DMEM | Cytiva | Cat#SH30022.01 |
| Fetal Bovine Serum (FBS) | Biowest | Cat# S1620 |
| 4-hydroxytamoxifen (4-OHT) | Sigma-Aldrich | Cat#H7904 |
| Tamoxifen | Sigma-Aldrich | Cat#T10540-29-1 |
| D-(+)-Glucose solution | Sigma-Aldrich | Cat#G8769 |
| MitoSOX™ Red | Thermo Fisher Scientific | Cat#M36008 |
| CM-H<sub>2</sub>DCFDA | Thermo Fisher Scientific | Cat#D399 |
| **Critical commercial assays** |  |  |
| Ultra Sensitive Mouse Insulin ELISA Kit | Crystal Chem | Cat#90080 |
| Senescence Detection Kit | Abcam | Cat#ab65351 |
| **Deposited data** |  |  |
| Raw data of RNAseq | This paper | GEO: GSE223676 |
| Tabula Muris Consortium mouse single cell RNAseq data | Tabula Muris Consortium | https://figshare.com/ndownloader/files/27856758 |
| RNAseq gene TPMs from young and old human tissues (GTEx Analysis V8 release: RNAseq gene TPMs by tissue) | GTEx | https://gtexportal.org/home/datasets |
| **Experimental models: Cell lines** |  |  |
| MEFs | In this paper | Wang et al <sup>23</sup> |
| **Experimental models: Organisms/strains** |  |  |
| Mouse: B6.129S6-Myctm2Fwa/Mmjax | JACKSON LABORATORY | Strain # 032046-JAX |
| Mouse: B6.129-Gt(ROSA)26Sortm1(cre/ERT2)Tyj/J | JACKSON LABORATORY | Strain # 008463 |
| Mouse: C57BL/6 | JACKSON LABORATORY | Strain # 000664 |
| **Oligonucleotides** |  |  |
| Primers and probes for qRT-PCR | In this paper | Figure S1D |

| Software and algorithms |  |  |
| --- | --- | --- |
| ImageJ- Fiji | NIH | https://fiji.sc/# |
| FlowJo v10 | Becton-Dickinson Biosciences | https://www.flowjo.com/solutions/flowjo/ |
| R studio | Posit | https://posit.co/download/rstudio-desktop/ |
| GraphPad Prism v9.0 | GraphPad Software Inc. | https://www.graphpad.com |
| Clusterprofiler | Wu et al <sup>151</sup> | https://bioconductor.org/packages/release/bioc/html/clusterProfiler.html |
| ComplexHeatmap | Gu et al <sup>155</sup> | https://bioconductor.org/packages/release/bioc/html/ComplexHeatmap.html |
| ggplot2 | CRAN | https://ggplot2.tidyverse.org/index.html |
| survminer | CRAN | https://cran.r-project.org/web/packages/survminer/index.html |
| CLC Genomic Workbench version 21 | QIAGEN | https://digitalinsights.qiagen.com/ |
| nf-core/rnaseq-3.4 | Ewels<sup>154</sup> | https://github.com/nf-core/rnaseq |
| Other |  |  |
| DNeasy Blood & Tissue Kit | QIAGEN | Cat#69504 |
| RNeasy kit | QIAGEN | Cat#74004 |
| QIAzol Lysis Reagent | QIAGEN | Cat# 79306 |
| RNeasy Lipid Tissue Mini Kit | QIAGEN | Cat#74804 |
| SuperScript™ IV First-Strand Synthesis System | Thermo Fisher Scientific | Cat#18091050 |
| 3-12% Native PAGE Novex Bis-Tris gel | Thermo Fisher Scientific | Cat#BN1003BOX |

### Resource availability

#### Lead contact

All additional information and requests for resources, reagents and methods should be directed to the lead contact, Edward V. Prochownik

#### Materials availability

All unique reagents generated in this study will be made available from the Lead Contact (E.V.P.) and may require a completed Materials Transfer Agreement.

#### Data and code availability

All raw RNAseq files have been deposited in the NCBI Gene Expression Omnibus ^156^ and are accessible through GEO Series accession number GSE223676 database (https://www.ncbi.nlm.nih.gov/geo/query/acc.cgi?acc=GSE223676).

This paper does not report original code. Data underlying the display items in the manuscript, related to Figure 1, Figure 2, Figure 3, Figure 4, Figure 5, Figure 6, Figure 7, and S1–S7 are available as Data S1 – Source data. The original full length western blots for Fig. 3I, Fig. 5C, and Fig. S1G are available as Data S2.

Any additional information required to reanalyze the data reported in this paper is available from the lead contact upon request.

### Experimental model and study participant details

#### Animal models

Animal work was conducted in compliance with the Public Health Service Policy on Humane Care and Use of Laboratory Animal Research (ILAR) Guide for Care and Use of Laboratory Animals. All experimental procedures, diets and tests were approved by the Institutional Animal Care and Use Committee (IACUC) at the University of Pittsburgh. All mice were housed in a specific pathogen-free facility, maintained under standard conditions at UPMC Children’s Hospital of Pittsburgh. The B6.129S6-*Myc^tm2Fwa^*/Mmjax mouse strain, in which the second and third exons of the *Myc* gene are flanked by loxP sites, was originally obtained as a gift from I. Moreno de Alboran. ^13, 14, 29^ These were crossed with the B6.129-*Gt(ROSA)26Sortm1(cre/ERT2)Tyj*/J strain, which expresses a Cre recombinase-estrogen receptor (*CreER*) fusion transgene under the control of the ROSA26 promoter. ^40^. 2 *Myc^LoxP/LoxP^* progeny strains were derived, containing one or 2 *CreER* transgene copies, which allowed for a determination of the efficiency of *Myc* excision in response to CreER dose. *CreER* activation and Myc excision were initiated at the time of weaning in mice that had attained a weight of 15 g or greater. Each mouse received 5 daily i.p. injections of freshly-prepared tamoxifen (75 mg/Kg) in corn oil. To ensure the complete metabolism and excretion of tamoxifen and to avoid any of its non-specific side effects, we allowed at least 8 wks before initiating any testing other than that specifically designed to confirm the extent of *Myc* exon 2 excision and full-length *Myc* transcript expression (Figure S1). As a further control for any long-term effects of tamoxifen treatment, control (WT) mice for all studies consisted of the offspring of matings between B6.129S6-*Myc^tm2Fwa^*/Mmjax and wild-type C57BL/6 mice treated with tamoxifen in the manner described above.

*Myc* excisional efficiency was determined using a quantitative TaqMan-based qPCR assay that compared the exon 2: exon 1 ratio using tissues from the above mice and standard curves generated with known ratios of WT and *Myc*KO DNAs as described previously (Figure S1A-E). ^13, 14, 29^ Cre-ER transgene copy number was determined by a separate TaqMan-based assay using the primers listed in Figure S1D. 10 ng of total DNA was used in each TaqMan assay. 3 primer sets were designed to amplify regions to identify specifically unfloxed, floxed (WT) and *Myc*KO alleles. All primers and probes (Figure S1D) were synthesized by IDT, Inc. (Coralville, IA). PCR reactions were performed on CFX96 Touch^TM^ Real-Time PCR Detection System (Bio-Rad, Inc.) using the following conditions: 95°C for 5 min; 10 cycles at 95 °C for 20 s, and 65 °C∼60 °C (decreasing by 0.5 °C per cycle) for 15 s, and 68°C for 10 s; 40 cycles at 95 °C for 15 s, and 60 °C for 1 min.

## Method details

### Derivation and propagation of primary murine embryo fibroblasts (MEFs)

Briefly, 10-12 e14 embryos from pregnant WT mothers were decapitated, eviscerated, rinsed in PBS, placed into sterile 0.25% trypsin-EDTA and incubated 1 hr at 37C as described previously. ^23, 157^ They were then finely minced and digested for an additional 1-2 h at 37C before transferring to fresh Dulbecco’s modified minimum essential medium (DMEM) containing 10% FBS, 100 mM glutamine and penicillin/streptomycin as previously described.^70^ After expanding for 3-4 days, these early passage cells were trypsinized and frozen at −80C to serve as subsequent stocks. These primary MEFs were designated as passage 1. To excise the floxed *Myc* alleles from the above cells, *in vitro* culturing was continued in fresh medium containing 500 nM 4-hydroxytamoxifen (4-OHT) (Sigma-Aldrich, St. Louis, MO), which was changed daily. On day 8 an aliquot of cells was harvested, DNA was isolated as described below and the ratio of WT and *Myc*KO Myc alleles was calculated using the same approach as described above for individual mouse tissues. Under these conditions, *Myc* allele excision routinely exceeded 95%. ^23^

### Strength and endurance testing

Strength testing was performed using a Grip Strength Meter (Harvard Apparatus, Holliston, MA) according to the direction of the supplier. Rotarod testing (SPW Industrial, Laguna Hills, CA) was based on a modification of the standard operating procedure from Jackson Laboratories: https://www.jax.org/-/media/jaxweb/files/research-and-faculty/tools-and-resources/peripheral-neuropathy-resource/rotarod.pdf?la=en&hash=78228ECB294E38BC773843500CDE2E8C99A96316.

Briefly, animals were initially placed on the slowly rotating rod (5 rpm) and maintained at this speed for 20 sec. The speed was then increased by 5 rpm increments each lasting 20 sec. The recorded numbers indicate the total time that each mouse was able to maintain its balance.

### Treadmill performance

Treadmill performance was monitored with a Columbus Instruments Exer 3/6 apparatus (Columbus, OH). Groups of 6 mice at a time (3 WT and 3 *Myc*KO) were evaluated according to a published protocol. ^158^ Briefly, mice were allowed to run along a treadmill (elevated 10° from the horizontal) at a gradually increasing pace until reaching exhaustion, which was defined as the time at which they preferred to rest for >5 sec. upon an immobile metal shock plate at the bottom of the treadmill. The total distance run until reaching the point of exhaustion was recorded for each animal.

### Metabolic cage profiling

These were performed essentially as described previously. ^29^ Briefly, control and *Myc*KO mice of the indicated ages were housed individually in metabolic cages (Columbus Instruments) and allowed to acclimate for 24 hr while being provided *ad lib* access to water and a standard mouse chow containing 5% fat (Picolab 5053; LabDiet, St. Louis, MO, USA). VO_2_ and VCO_2_ were recorded every 20 min over the subsequent 48 hr along with food intake and overall activity. At the conclusion of this observation period, mice were starved overnight (12 hr) and then provided with a standard diet for 24 hr followed by a high-fat diet (45%) for an additional 24 hr while again monitoring RERs. Data analyses were performed with a web-based Analysis software package CalR (https://calrapp.org/cite.html).

### Glucose tolerance tests and serum glucose, lactate and ketone measurements

Mice were fasted for 5 hr. at which time whole blood glucose, lactate and ketone levels were obtained using meters and compatible strips according to the directions provided by the suppliers (Glucose AimStrip Plus, Germaine Laboratories, Inc. San Antonio, TX; Lactate Plus Analyzer, Sports Resource Group, Inc., Hawthorne NY; Keto-Mojo Ketone Meter, Keto-Check, Inc. Napa, CA). To perform glucose tolerance tests and to measure insulin levels, the above mice were injected with 2g of dextrose/kg body mass with blood glucose levels being subsequently measured at the indicated times. Serum insulin levels were measured using an Ultra Sensitive Mouse Insulin ELISA Kit according to the directions provided by the supplier (Crystal Chem, Elk Grove Village, IL).

### ImageJ quantification of ORO staining

ORO- and hematoxylin-stained tissue sections were imaged on a Leica DFC7000T microscope with 5x and 40x magnification. Multiple overlapping images of each section were acquired for the full area. The images of each section were joined using the stitching plugin of the open source software FIJI. ^159–161^ After subtracting background from each image, color de-convolution ^162^ was performed in FIJI where the colors were specified in advance from ROIs respectively corresponding to unstained tissue, strongly stained tissue and the slide background. Quantification of Oil-Red-O positive staining was performed as described in ImageJ documentation (https://imagej.nih.gov/ij/docs/examples/stained-sections/index.html). Higher resolution images were acquired at 5x magnification(Figure 2C).

### Nucleic acid isolation

DNAs and RNAs were isolated from mouse tissues using DNeasy and RNeasy kits, respectively according to the directions of the supplier (Qiagen, Inc. Germantown, MD). Exceptions to this were made in the case of adipose tissue and skeletal muscle for which we utilized a RNeasy Lipid Tissue extraction Kit and QIAzol Lysis Reagent (Qiagen, Inc., Germantown, MD), respectively. Total RNAs were reverse transcribed using a SuperScript IV First-Strand Synthesis System according to the directions of the supplier (Thermo Fisher Scientific, Pittsburgh, PA). To determine the degree of Myc transcript reduction in control and *Myc*KO tissues, 2 separate TaqMan-based qRT-PCR assays were performed that compared the exon 2: exon 1 ratio signals in each WT and *Myc*KO tissue (Figure S1D&E).

### Blue native gel electrophoresis (BNGE), *in situ* enzymatic assays for ETC enzymatic function

Non-denaturing gel electrophoresis was performed largely as described previously. ^56, 73^ Briefly, purified mitochondria (approx. 1 mg of total protein), were lysed by the addition of digitonin and then incubated on ice for 20 min. Coomassie blue solution (5% Coomassie blue G250 in 750 mM 6-aminocaproic acid) was added and the suspension was then centrifuged at 14,000× *g* for 20 min at 4°C. The supernatant was diluted in the supplier’s buffer, loaded onto a 3-12% Native PAGE Novex Bis-Tris gel (Life Technologies, Carlsbad, CA) and electrophoresed for 4 hr at 4C at 80 V. Gels were then stained with Bio-Safe Coomassie G250 (Bio-Rad, Hercules, CA) for 30 min and de-stained exhaustively in deionized water. Stained gels were scanned and the imaged using an AlphaEaseFC 2200 scanner and AlphaEaseFC software. Enzymatic assays for mitochondrial complexes and super-complexes were performed as previously described for Complex I (NADH ubiquinone oxidoreductase), Complex III (CIII) (decylubiquinol cytochrome c oxidoreductase), Complex IV (CIV) (cytochrome c oxidase) and Complex V (ATPase). ^73^ Band intensities were measured and quantified using Image J software and normalized with their corresponding bands on the Coomassie stained blue native gel.

### ROS assessment

CM-H_2_DCFDA and MitoSOX™ Red dyes were utilized to measure reactive oxygen species (ROS) levels (Molecular Probes, Eugene, OR, USA) according to the manufacturer’s protocol. This was achieved by exposing monolayer cultures of mouse embryonic fibroblasts (MEFs) maintained at a temperature of 37 ℃. Quantifications were performed on 6 biological replicates comprising 20,000 cells/sample using a BD LSRII flow cytometer (Becton-Dickinson Biosciences, San Jose, CA, USA) and results were analyzed using FlowJo v10 software. This was done as described in Wang et al. ^23^

### β-galactosidase staining

Tissue sections were stained for β-galactosidase using a Senescence Detection Kit (ab65351) according to the directions of the supplier (Abcam, Inc., Waltham, MA).

### SDS-polyacrylamide gel electrophoresis (SDS-PAGE) and immunoblotting

At the time of sacrifice, individual tissues were removed, and immediately placed on ice. They were then divided into small sections, snap-frozen in liquid nitrogen and maintained at −80C for long-term storage. To prepare samples for SDS page, tissue fragments were disrupted in PAGE buffer using a Bullet Blender as previously described. ^56, 163^ Protein concentration was quantified using the Bradford reagent (Bio-Rad, Inc., Hercules, CA). Electrophoresis, semi-dry blotting and protein detection was performed as previously described. ^56^ Antibodies used for the detection of specific proteins were used largely according to the directions of the suppliers and are shown in Table S3.

### Immunohistochemistry and immunohistofluorescence staining

All tissues were fixed in 10% formalin, paraffin embedded and cut into 4 μm thick sections for standard hematoxylin/eosin staining or immunostaining procedures as previously described (Wang 2022). Prior to staining for Myc, heat-induced antigen retrieval was performed using a citrate buffer (pH 6.0) for 30 minutes. Sections were incubated with a rabbit anti-Myc antibody (1:250; N262, SantaCruz) at 4C for 72 hours. A biotinylated secondary antibody was used to amplify the signal using an avidin–biotin substrate (Vector Laboratories, Inc., Newark, CA). Immunohistofluorescence staining for γH2AX was done as described in Wang et al ^23^.

### Transcriptional profiling

RNAs were purified from omental adipose tissue, liver and skeletal muscle as described above followed by DNAase digestion. ^13, 14, 23^ RIN values were determined using an Agilent 2100 Bioanalyzer (Agilent Technologies, Foster City, CA) and only those with values of >8.5 were processed further. Sequencing libraries were generated with a NEBNext Ultra Directional RNA Library Prep kit according to the supplier’s directions (New England Biolabs, Beverly, MA). Sequencing was performed as previously described on a NovaSeq 600 instrument (Illumina, Inc., San Diego, CA) by Novagene, Inc. (Sacramento, CA). ^13, 14, 23^ Original data were deposited in the NCBI Gene Expression database and are available through the Gene Expression Omnibus (GEO) ^156^ under accession number GSE223676.

To identify differentially expressed transcripts, we utilized CLC Genomic Workbench version 21(Qiagen) and mapped raw reads to the GRCm38.p6 mouse reference genome. Functionally related and differentially expressed groups were identified using clusterProfiler (R package version 4.2) ^151, 152^ by first screening the MSigDB data bases (http://www.gseamsigdb.org/gsea/msigdb/ as described previously. ^13, 14^ We also screened the Enrichr collection to identify additional groups of gene sets that were either absent from or underrepresented in MSigDB (http://amp.pharm.mssm.edu/Enrichr). ^148–150^

Representative gene sets along with their normalized enrichment score (NES) and q values were displayed graphically using the Ridgeline plot application from Clusterprofiler (https://rdrr.io/bioc/enrichplot/man/ridgeplot.html).

To identify non-canonically spliced transcripts, we utilized the nf-core/rnaseq-3.4 analysis pipeline with the percentage of non-canonical splices being calculated from multi-qc of STAR section pct_noncanonical_splices = num_noncanonical_splices/total_reads*100. ^153, 154^ Tabula Muris Consortium mosue single cell RNAseq data to evaluate the expression of Myc and Myc targets expression were obtained from https://figshare.com/ndownloader/files/27856758 and analyzed as described. ^104, 164^ Myc transcript levels in tissues obtained from young and old human tissues were downloaded from the GTEx Portal (GTEx Analysis V8 release: RNAseq gene TPMs by tissue) (https://gtexportal.org/home/datasets).165

### Quantification and statistical analysis

Quantification and statistical analysis were performed using R software v4.2.0 ^166^ (R Foundation for Statistical Computing, Vienna, Austria) and GraphPad Prism v9.00 (GraphPad Software Inc., USA). The ComplexHeatmap and ggplot2 packages were utilized for boxplot and heatmap visualizations, while the survminer package was used for survival curve plotting. The number of samples per group (n) for each experiment is indicated either in the figure legend or within the figure itself. A two-tailed, unpaired t-test was employed to assess significant differences between normally distributed populations, while a two-tailed Mann-Whitney exact test was used for non-normally distributed populations. A p-value below 0.05 was considered statistically significant. Significance is denoted as follows: * = p < 0.05, ** = p < 0.01, *** = p < 0.001, **** = p < 0.0001, and “ns” indicates not significant. Detailed statistical analysis information can also be found in each figure legend.

## Supplemental information

**Document S1.** Figures S1–S7, Table S3.

**Video S1.** External appearance WT and *Myc*KO mice, related to Figure 1.

**Table S1.** Efficiency and persistence of *Myc* gene excision and reduction of expression relative to comparably aged WT tissues, related to Result.

**Table S2.** List of the individual gene sets from Fig. 7A pertaining to mRNA splicing in WT and *Myc*KO mice, related to Figure 7A.

**Data S1.** Data underlying the display items in the manuscript, related to Figure 1, Figure 2, Figure 3, Figure 4, Figure 5, Figure 6, Figure 7, and S1–S7

**Data S2.** The original full length western blots, related to Fig. 3I, Fig. 5C, and Fig. S1G.

