## Supplementary material for "Premature Aging and Reduced Cancer Incidence Associated with Near-Complete Body-Wide *Myc* Inactivation": Data S2

20 month GAPDH Myc Fig 3-I

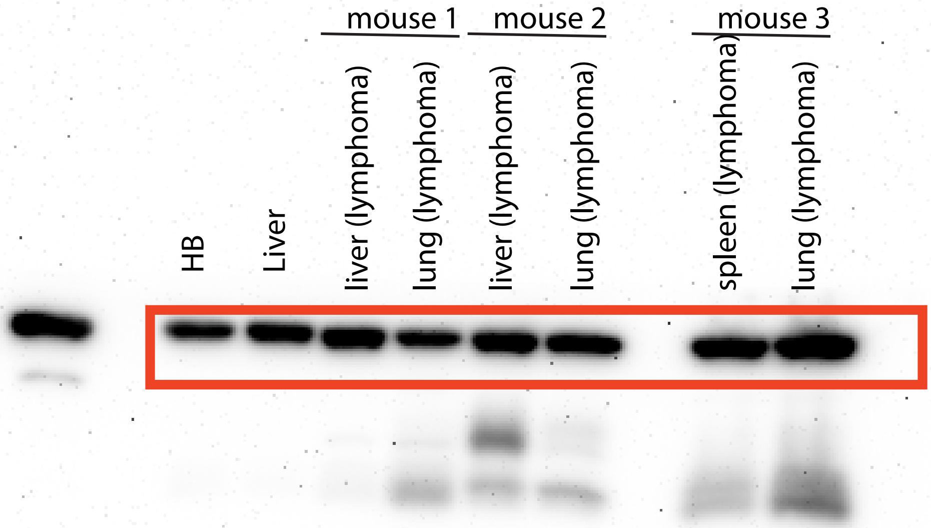

20 month tumor Myc Fig 3-I

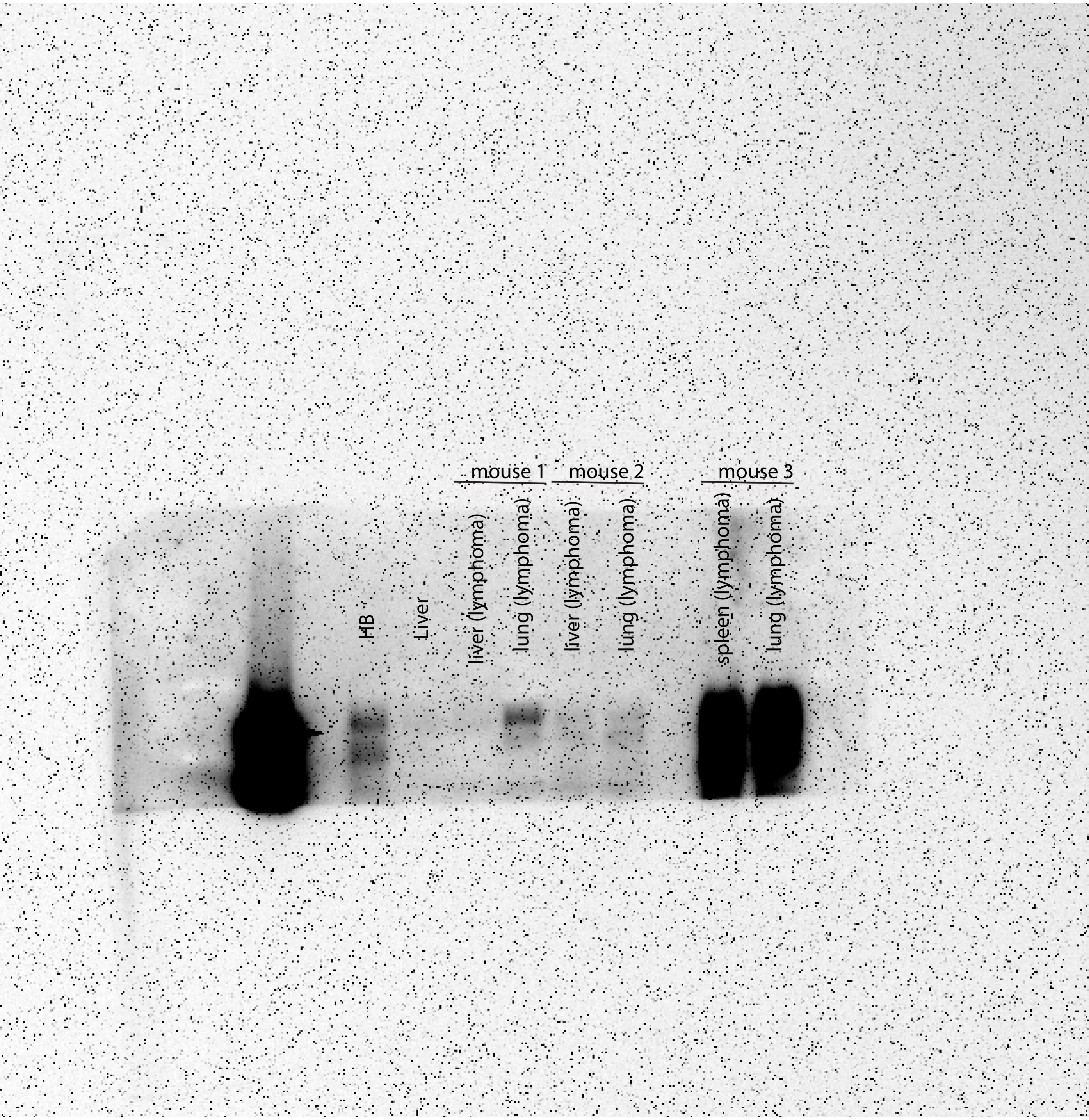

5 month liver PFK-L Fig 5-C

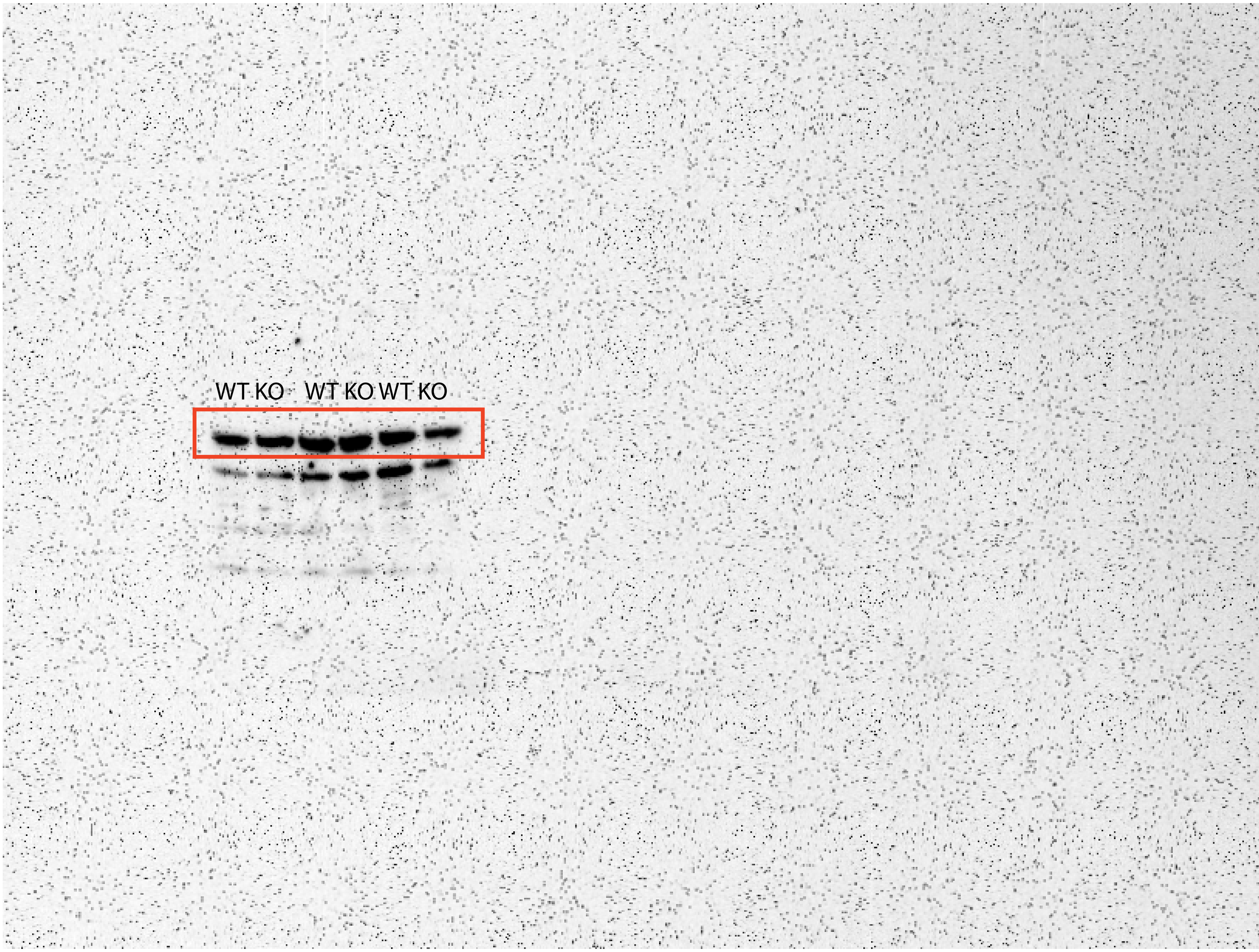

5 month Glut 1 Fig 5-C

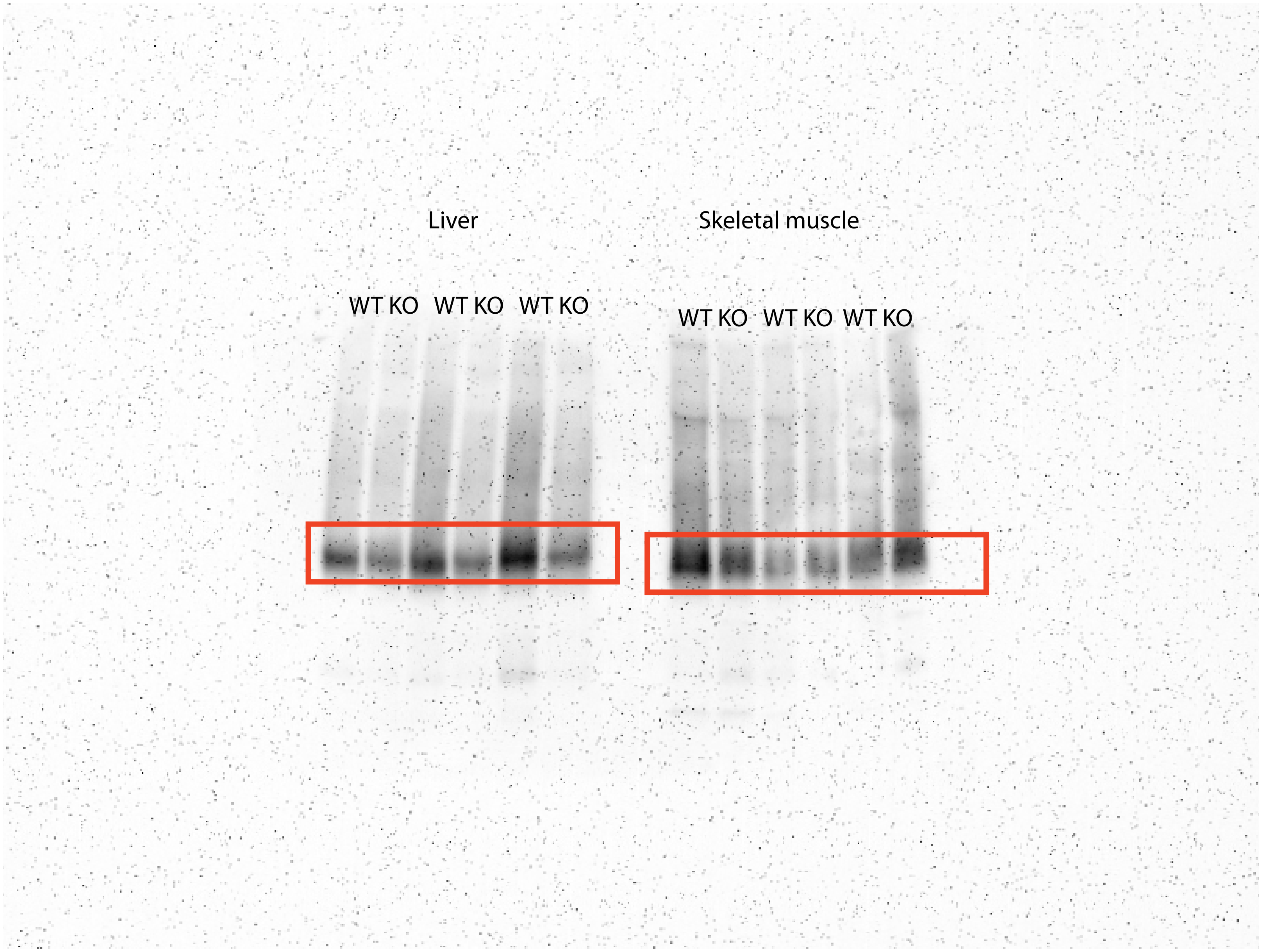

5 months liver GAPDH Fig 5-C

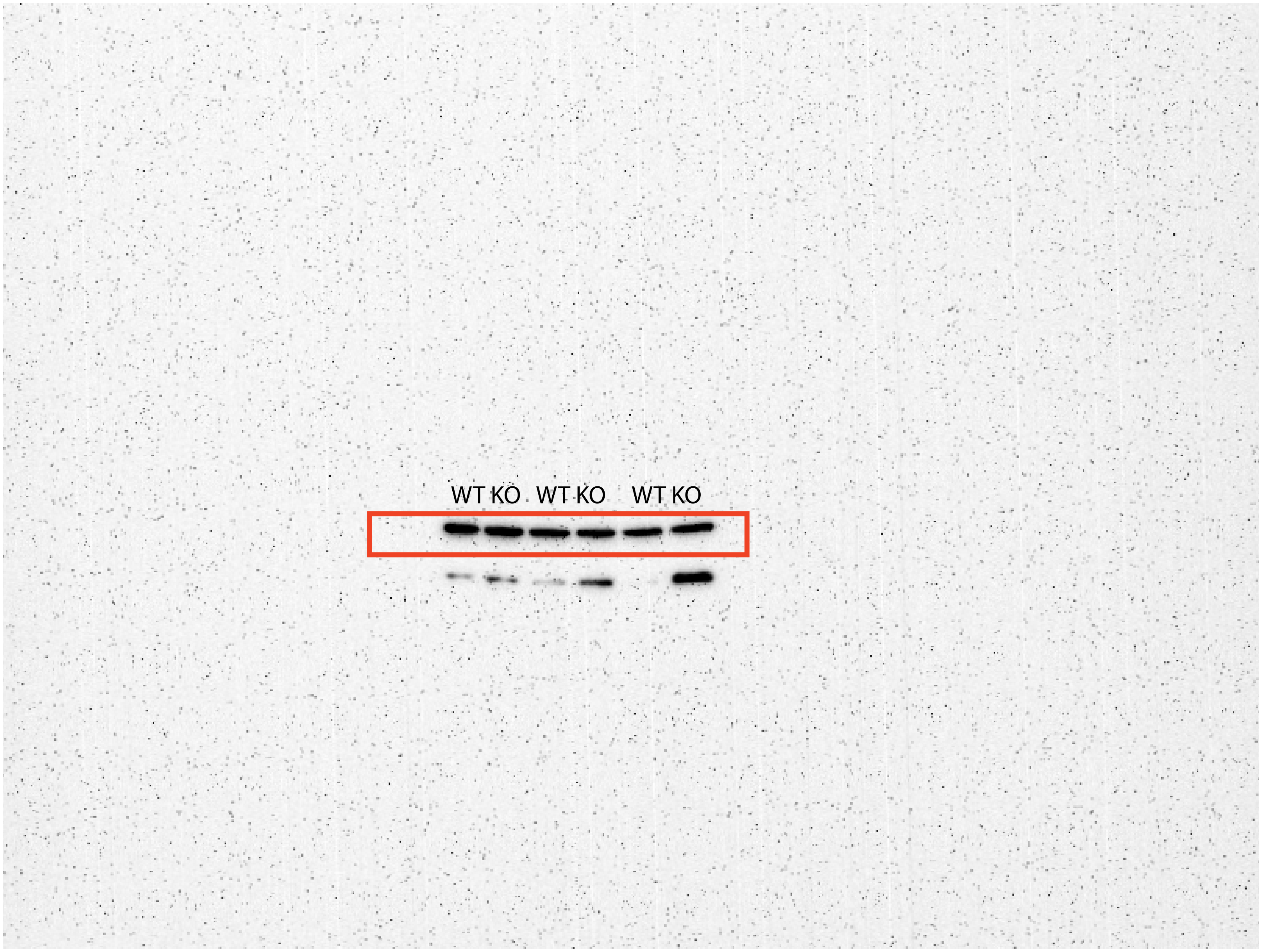

5 months liver Glut 2 Fig 5-C

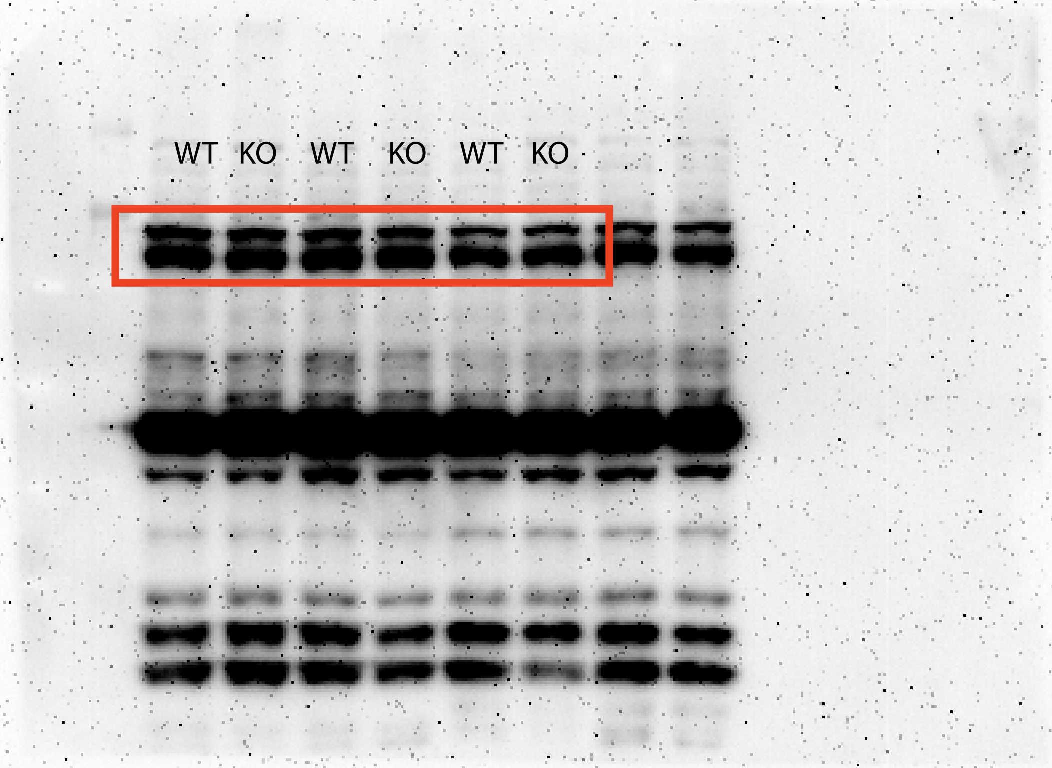

5 months liver glut 4 Fig 5-C

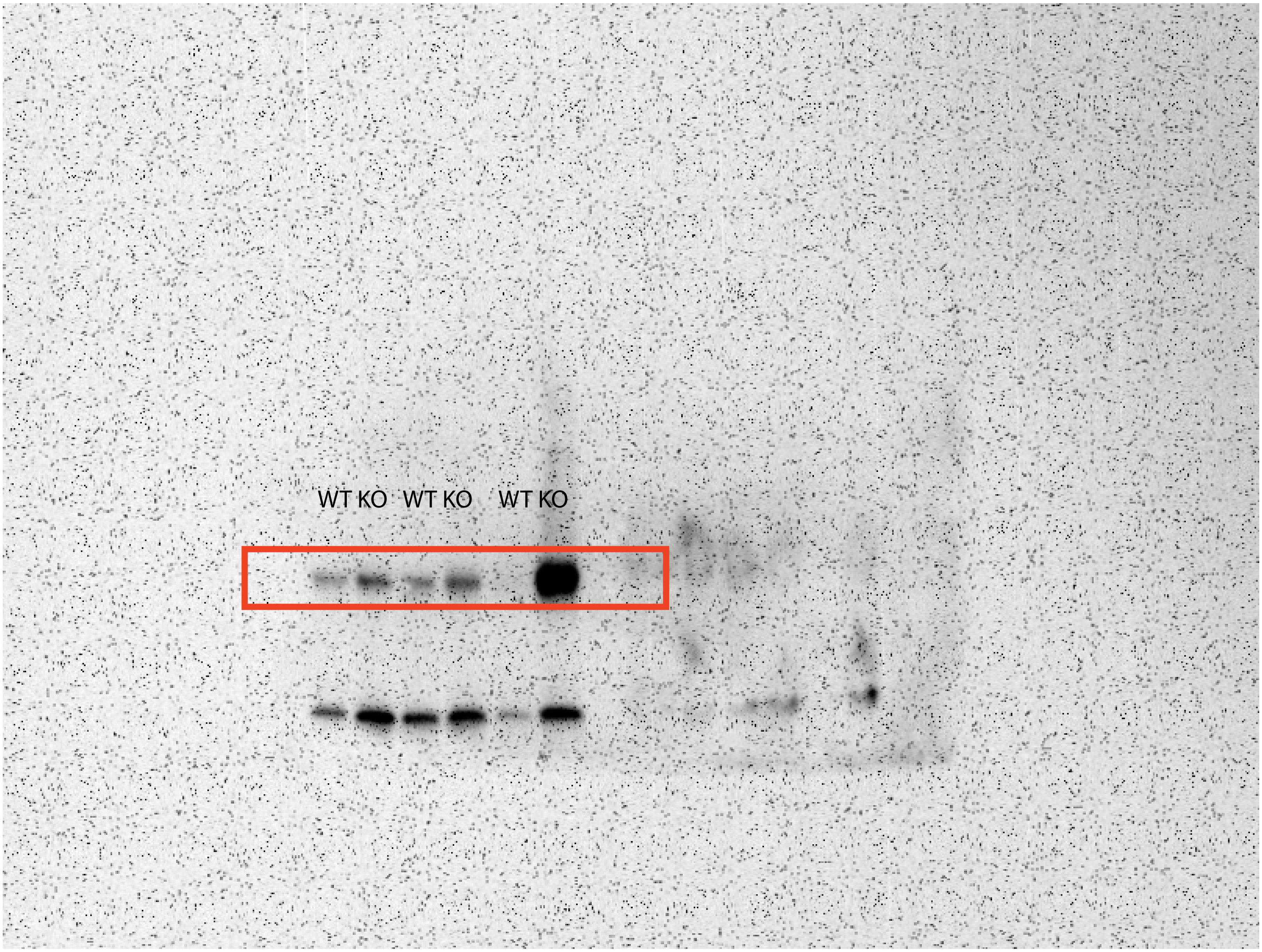

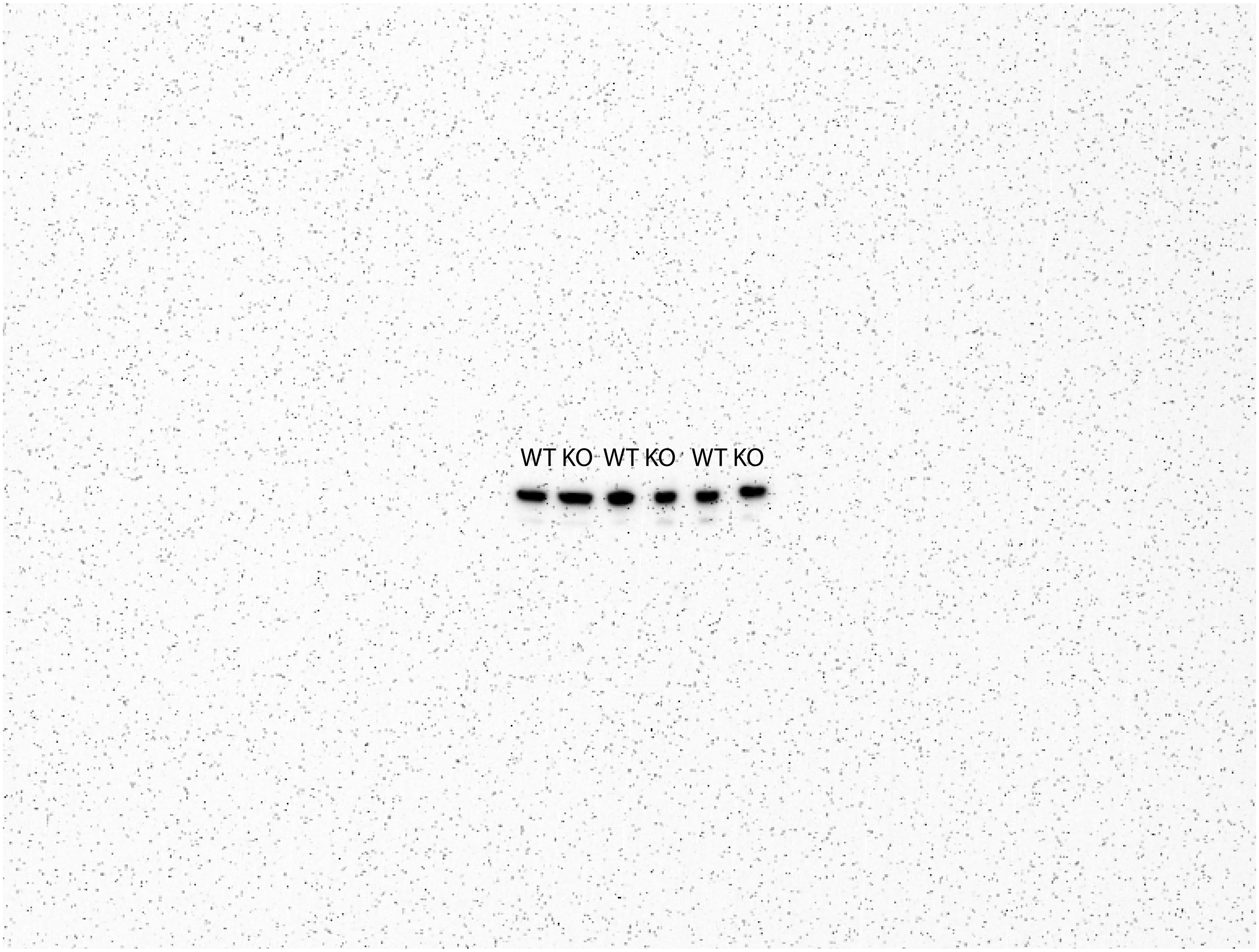
WT KO WT KO WT KO

5 months liver pPDH FIg 5-C

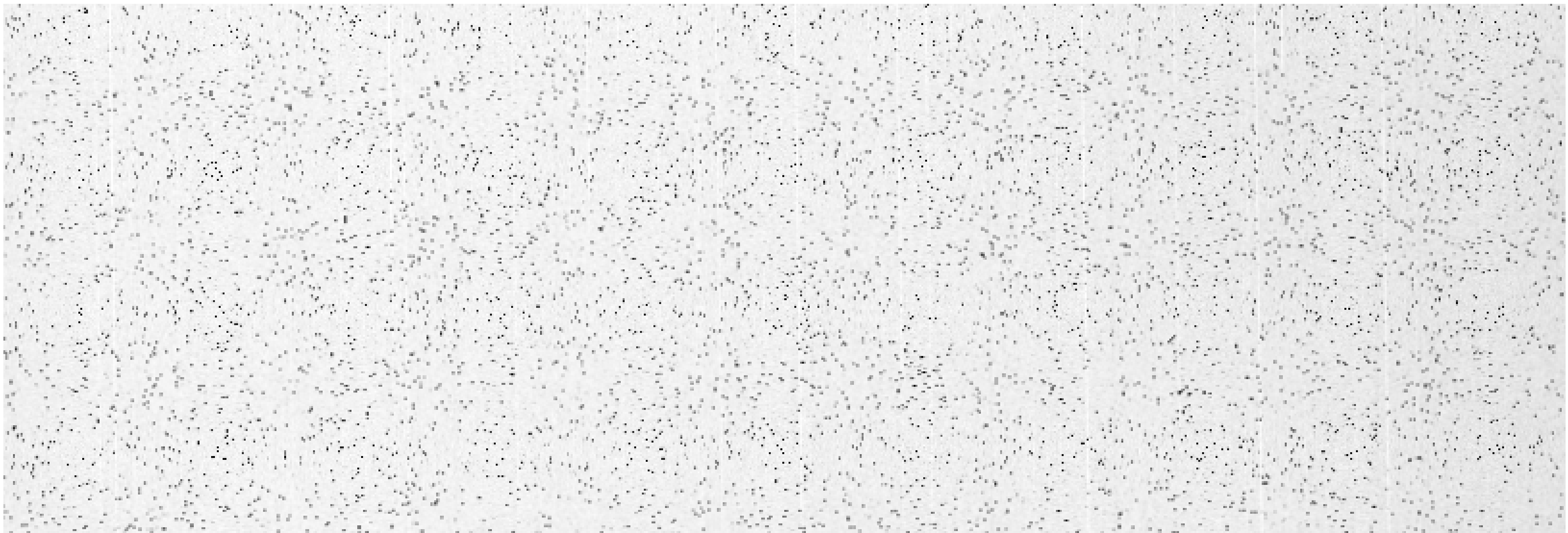

5 months skeletal muscle Glut 4 Fig 5-C

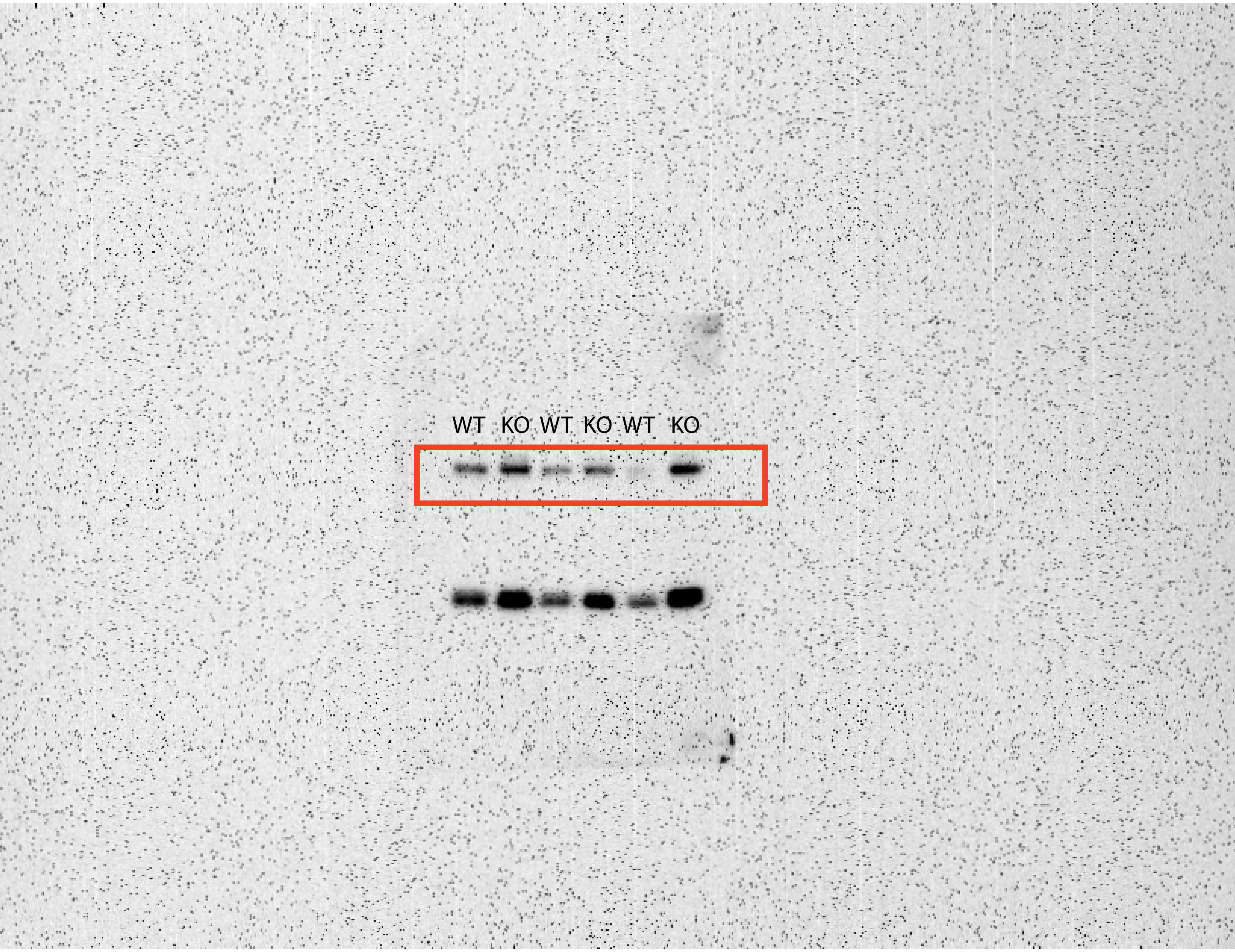

5 months skeletal muscle PDH Fig 5-C

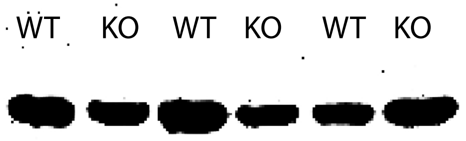

5 months skeletal muscle PFK-M Fig 5-C

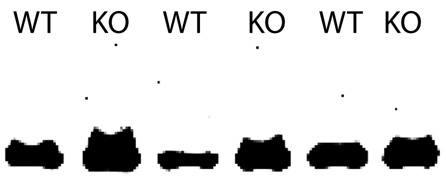

5 months skeletal muscle pPDH Fig 5-C

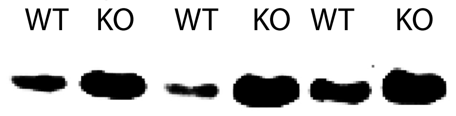

5 month skeletal muscle GAPDH Fig 5-C

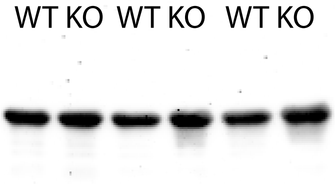

20 months liver GAPDH Fig 5-C

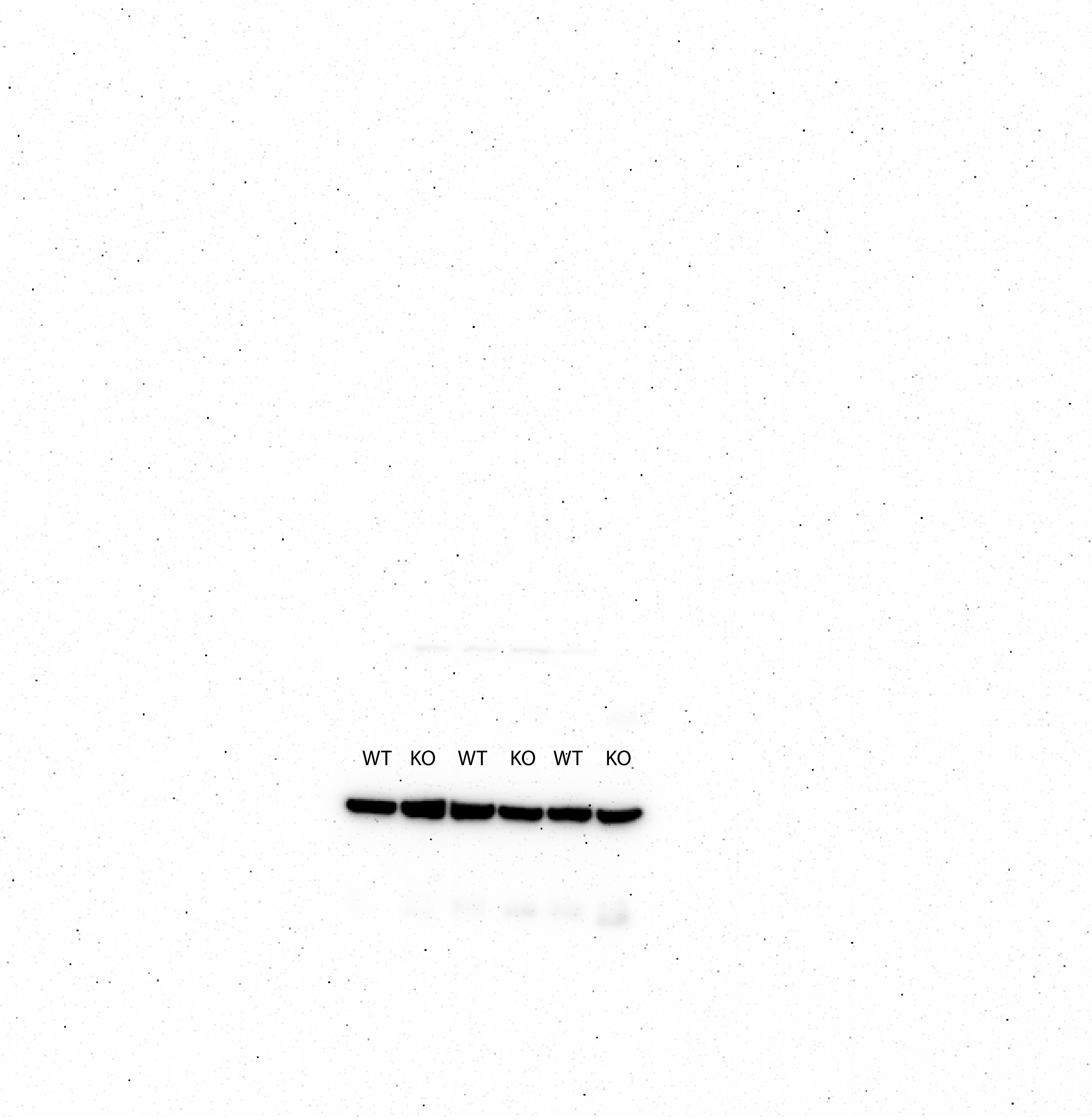

20 months liver Glut 1 Fig 5-C

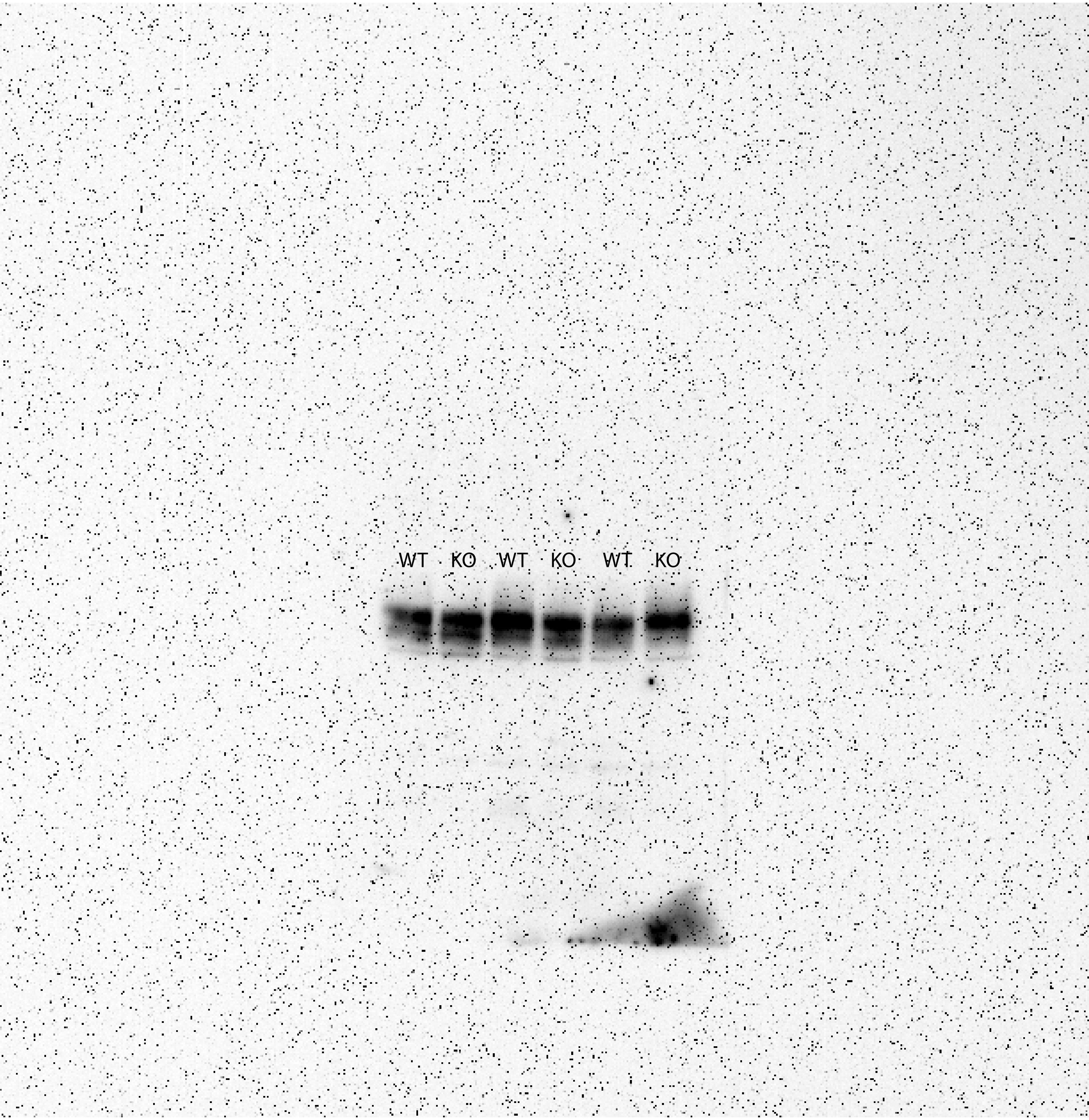

20 months liver Glut 2 Fig 5-C

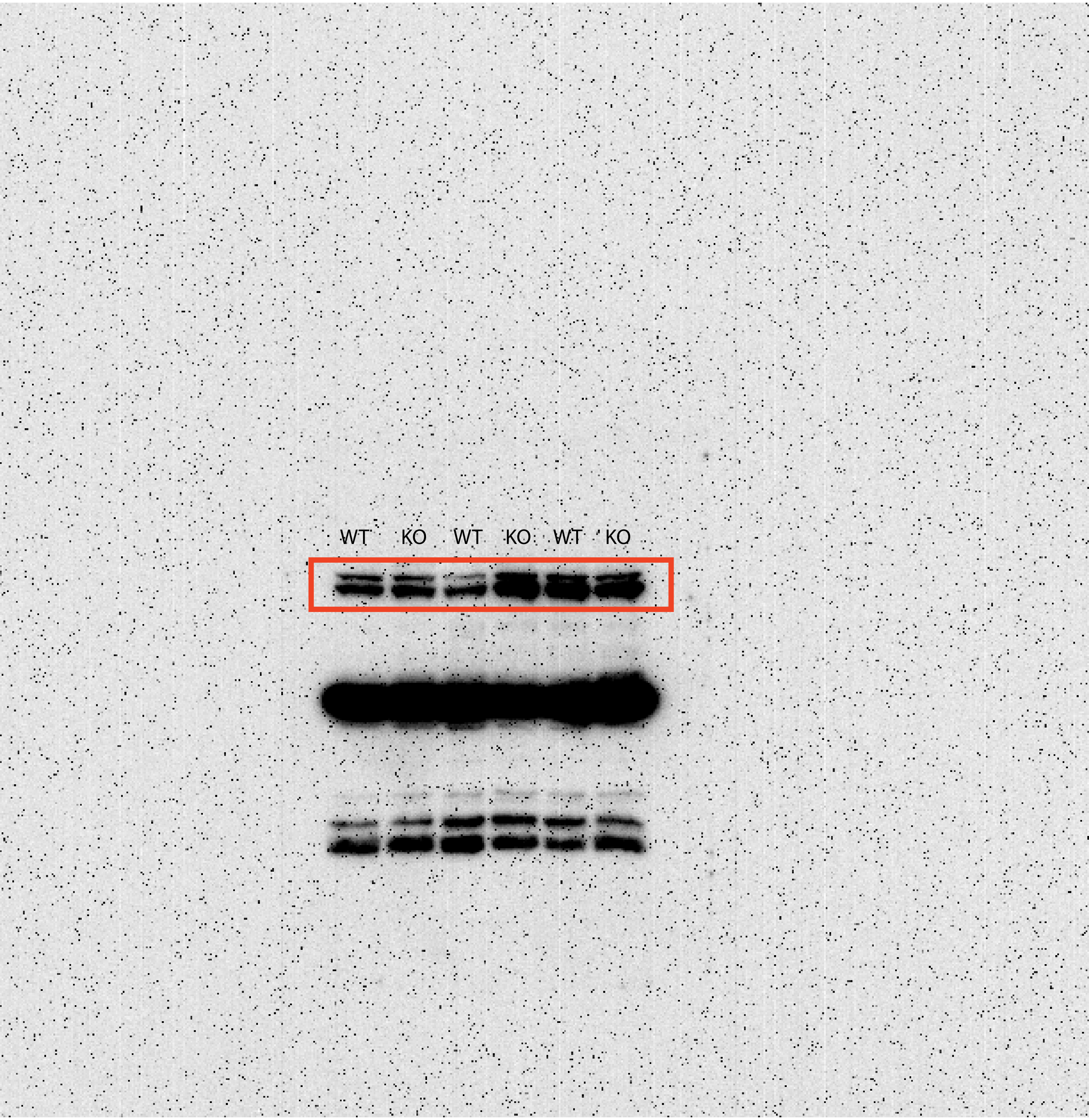

20 months liver Glut 4 Fig 5-C

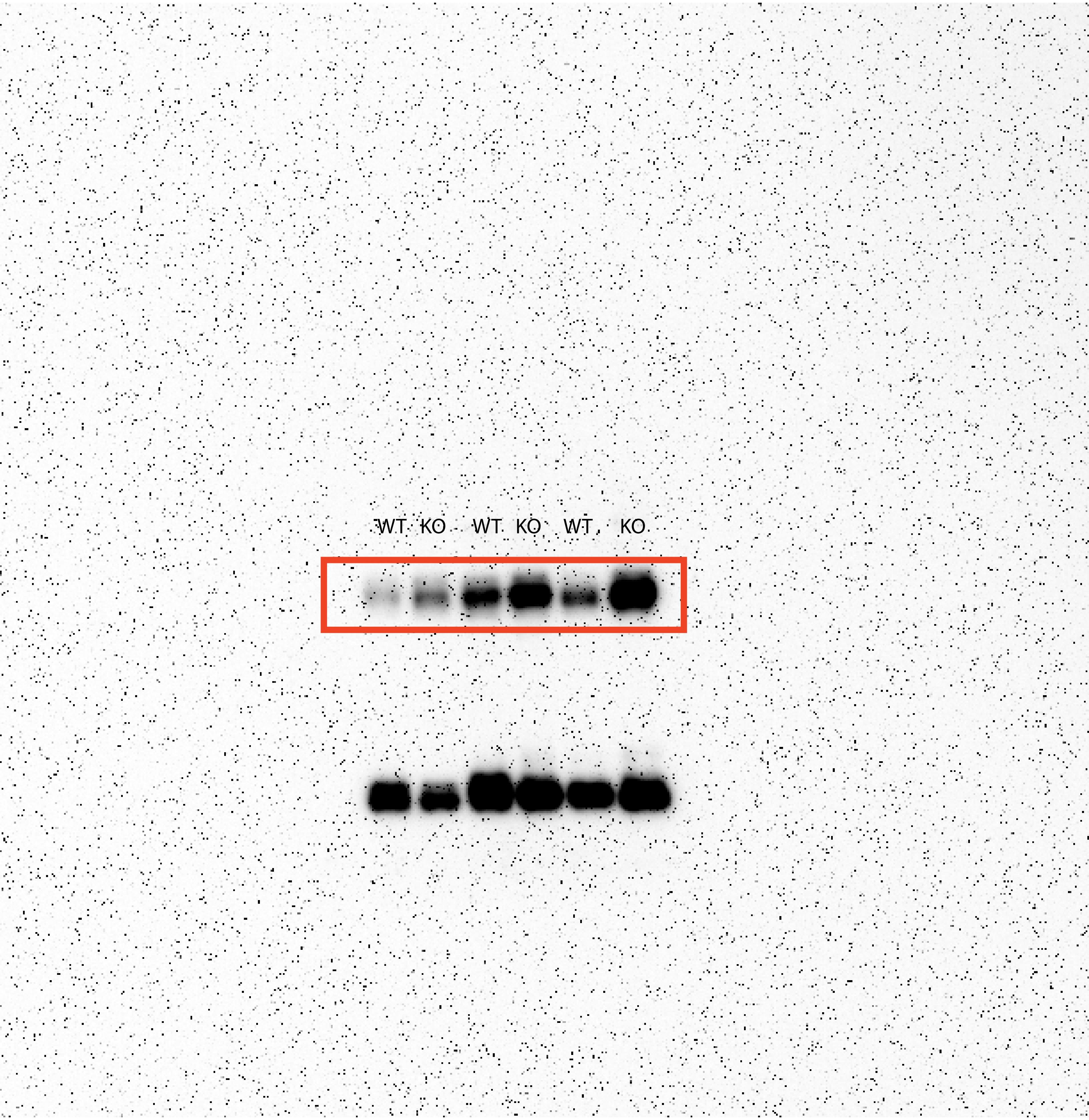

20 months liver PDH Fig 5-C

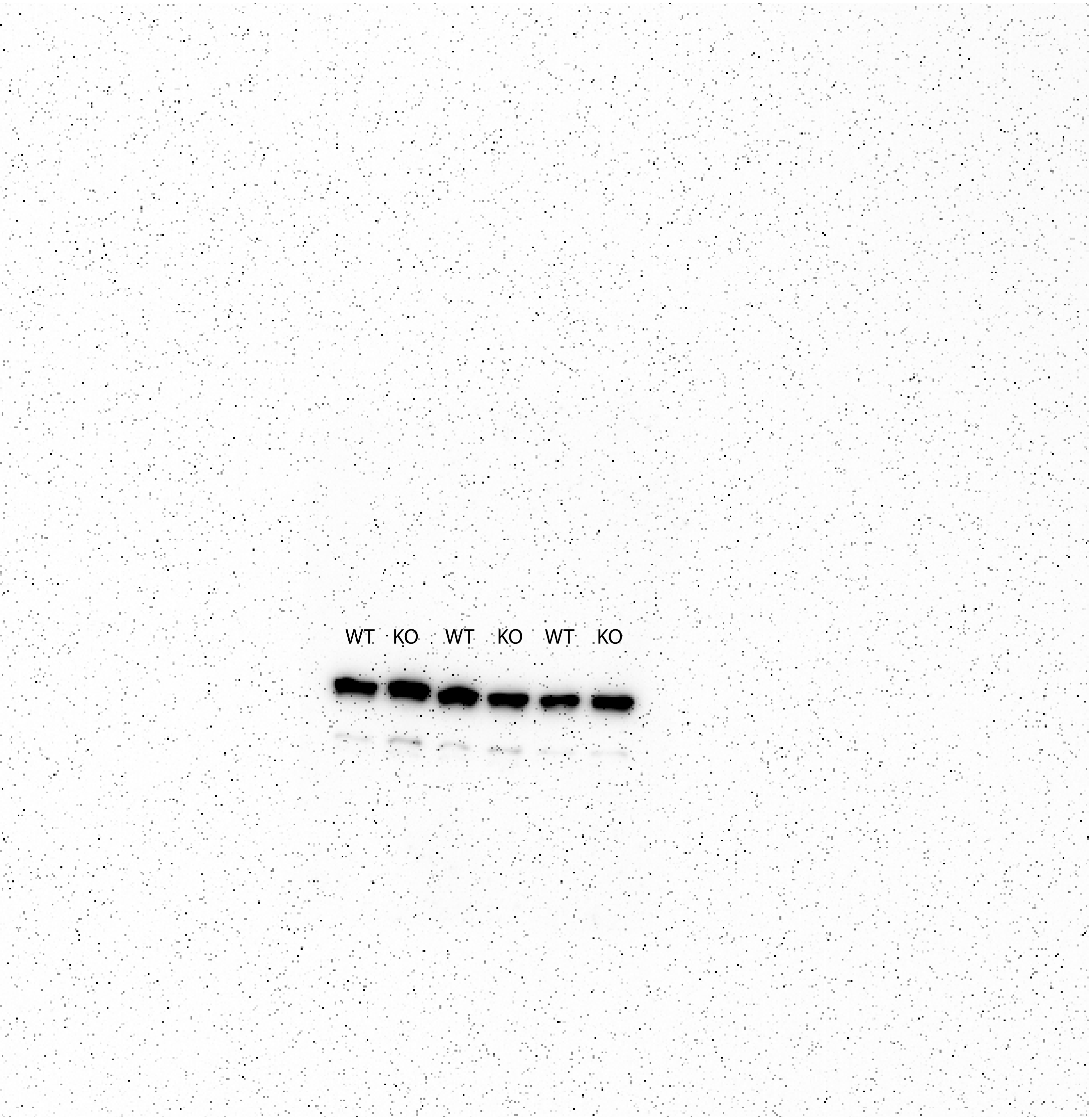

20 months liver PFK-L Fig 5-C

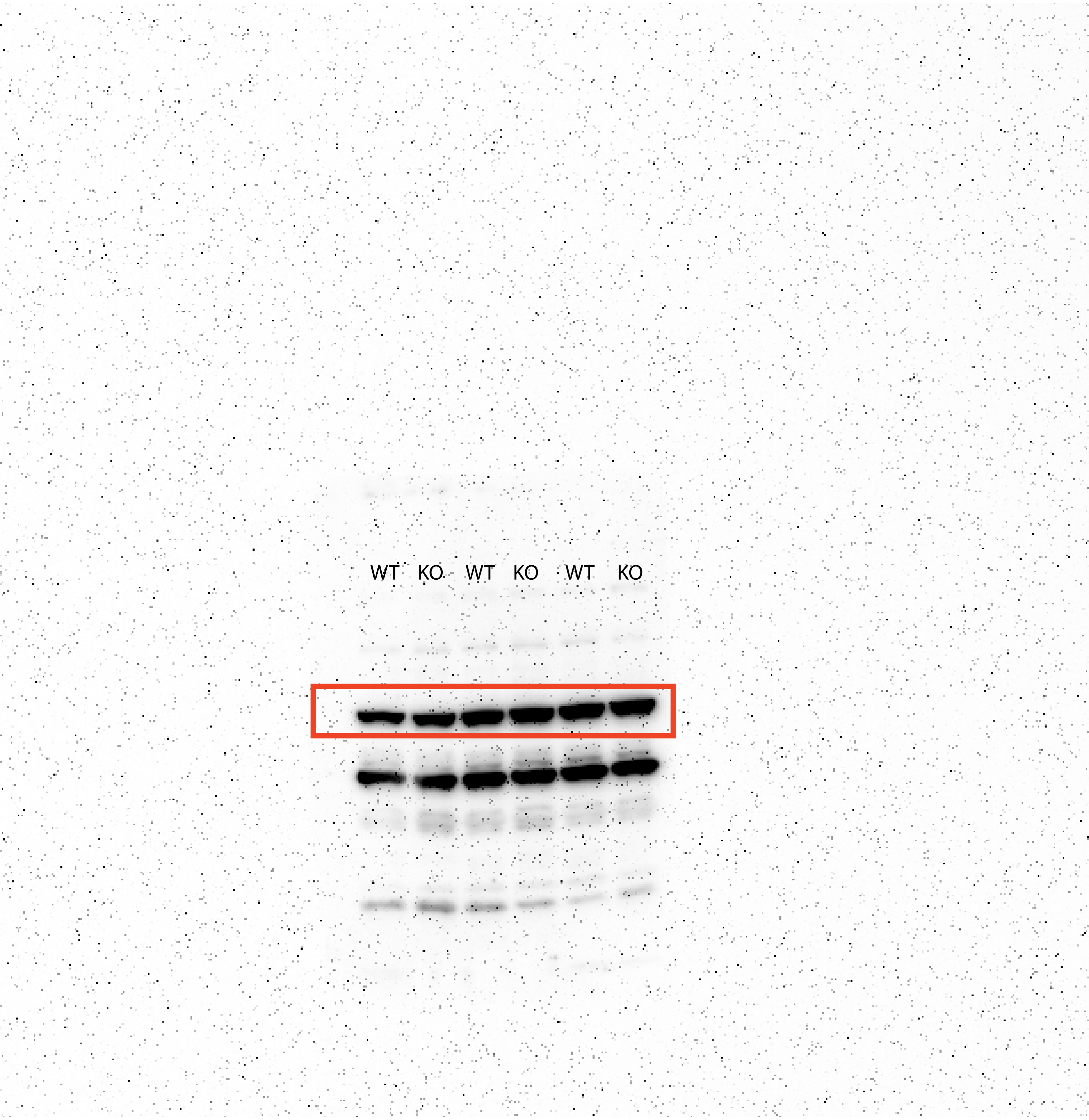

20 months liver pPDH Fig 5-C

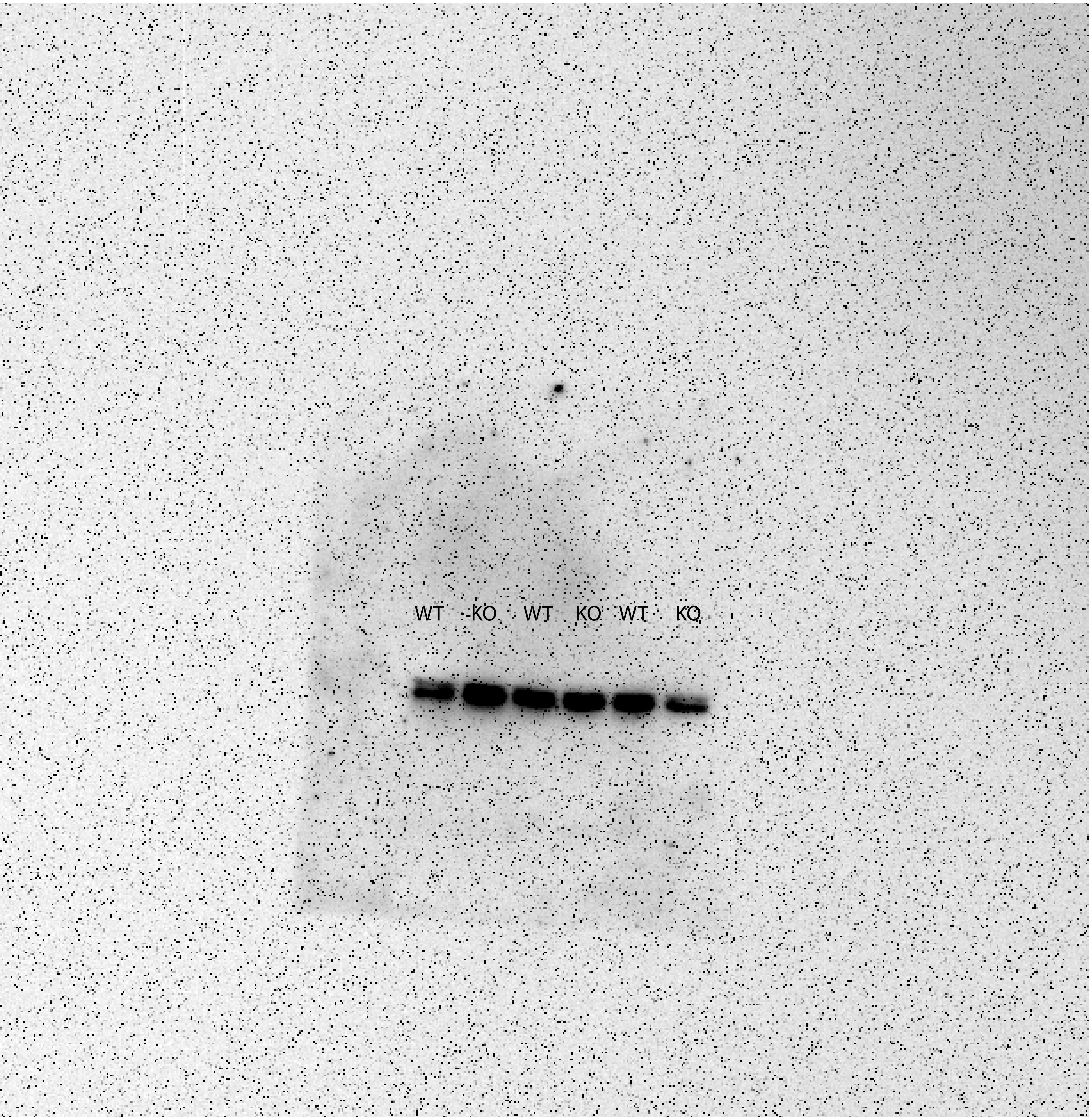

20 months skeletal muscle GAPDH Fig 5-C

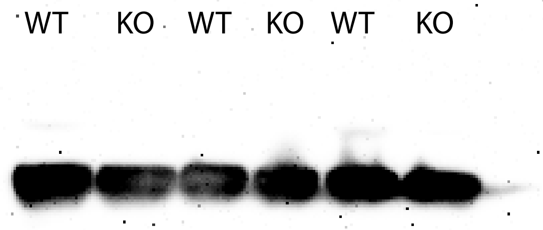

20 months skeletal muscle Glut1 Fig 5-C

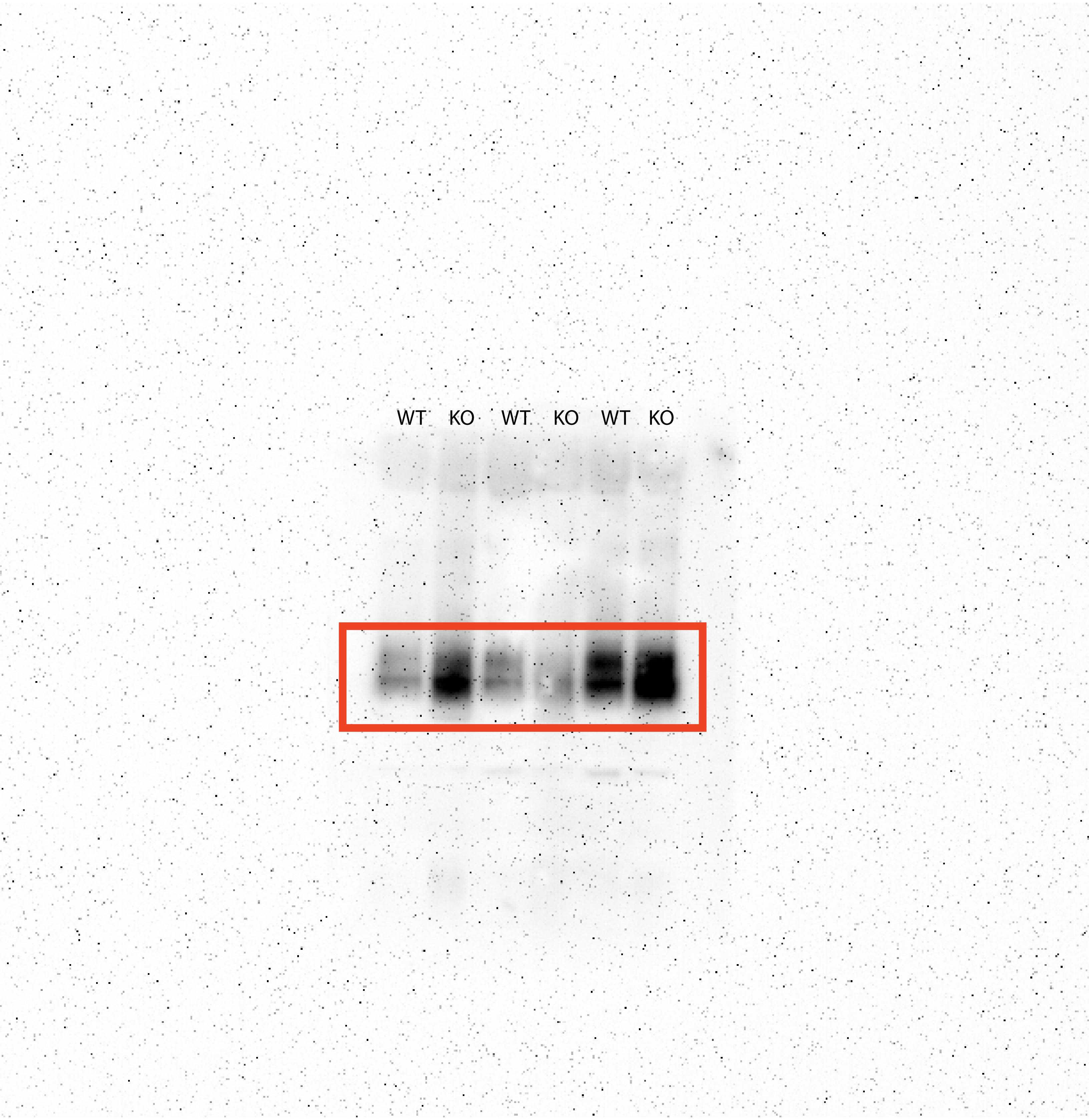

20 months skeletal muscle Glut 4 Fig 5-C

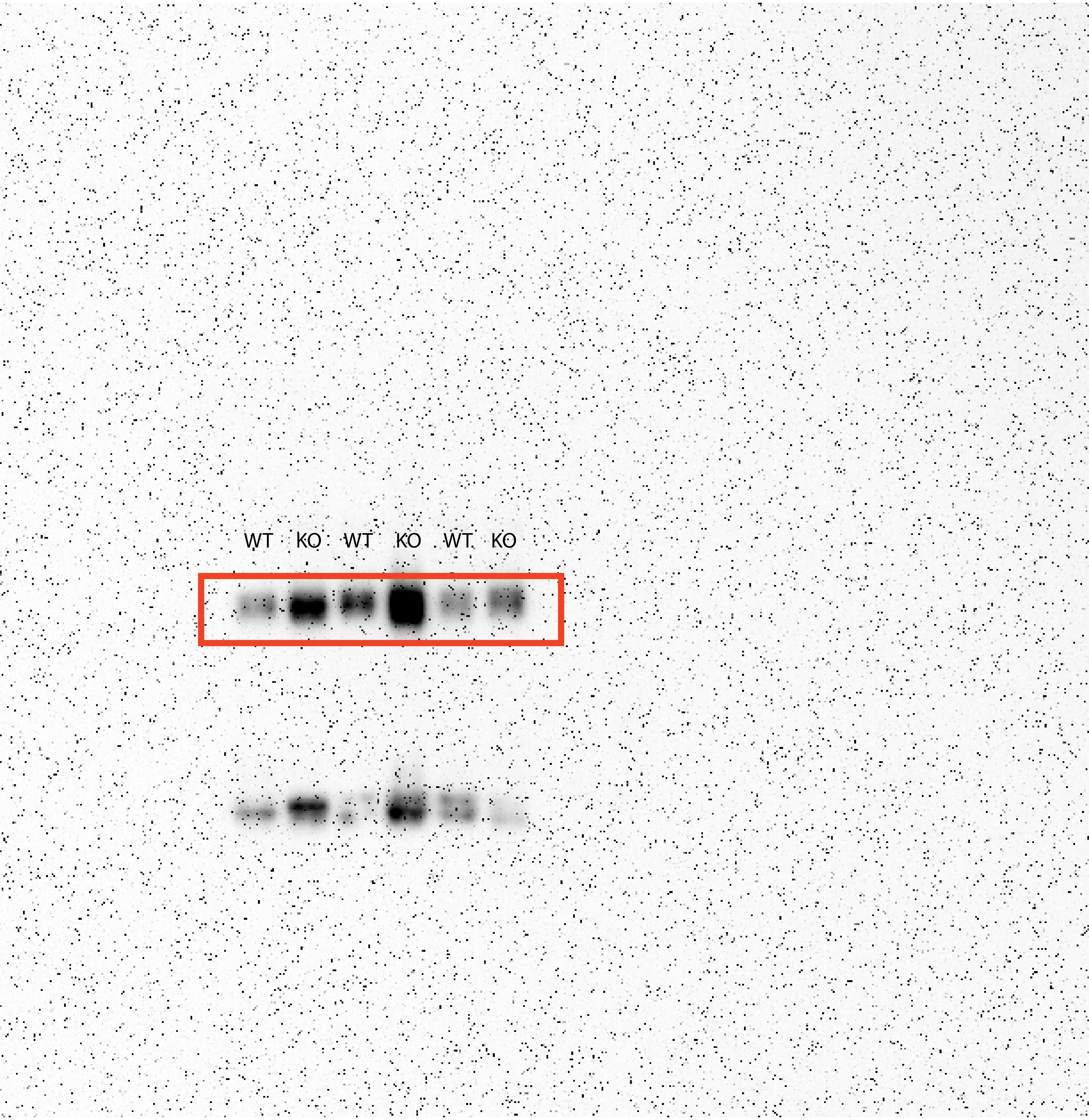

20 months skeletal muscle PDH Fig 5-C

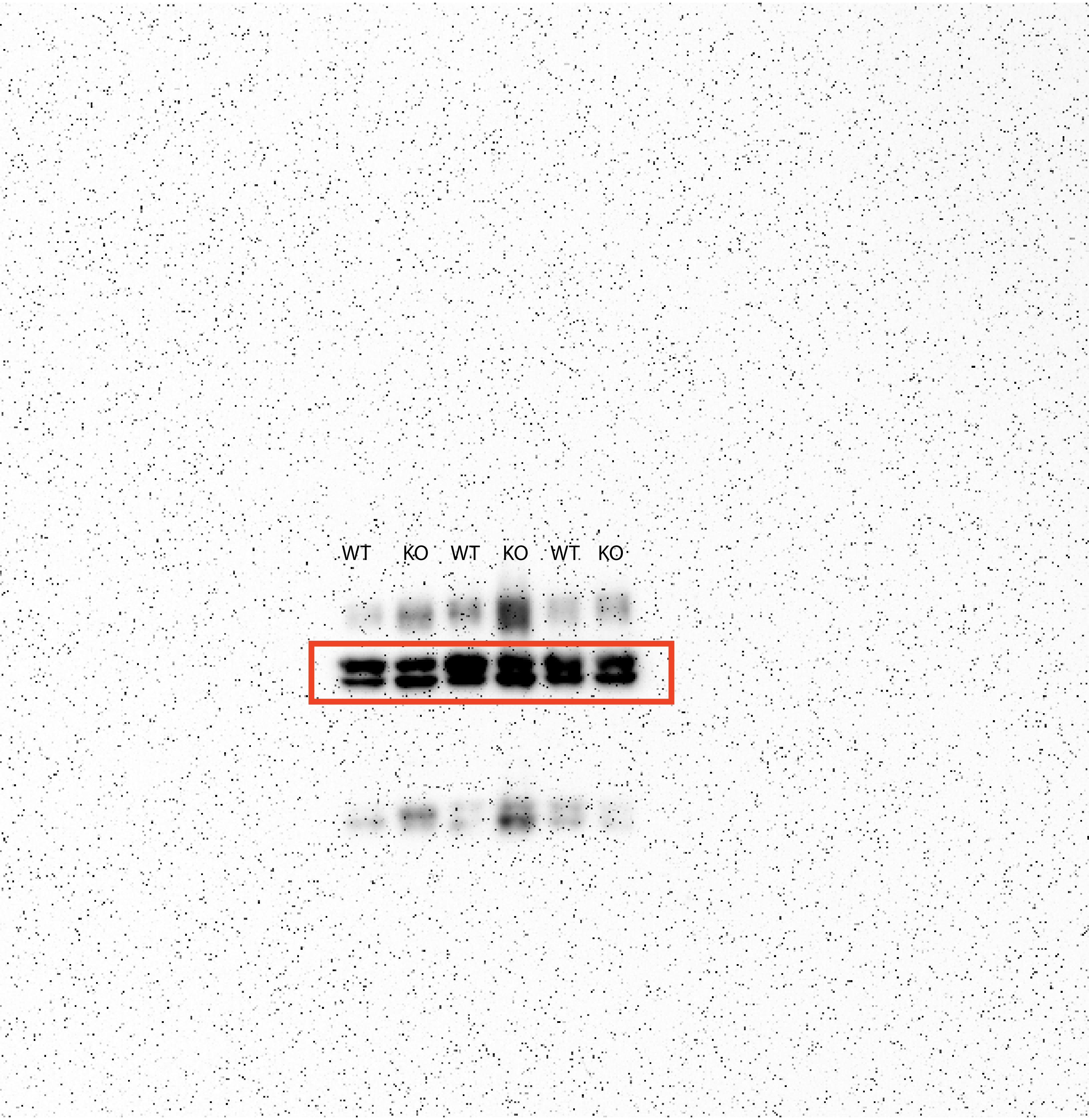

20 months skeletal muscle PFK-M Fig 5-C

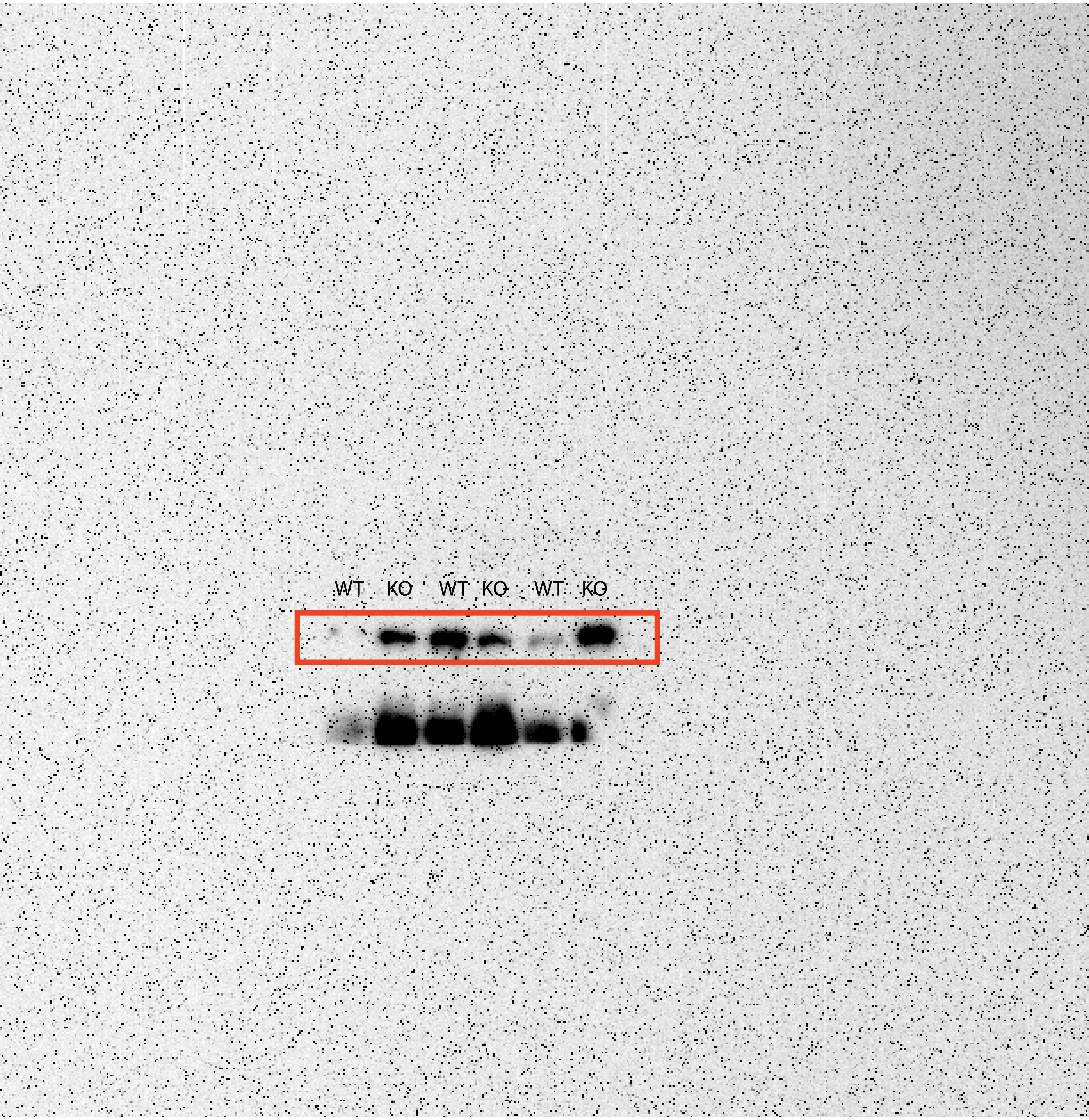

20 months skeletal muscle pPDH Fig 5-C

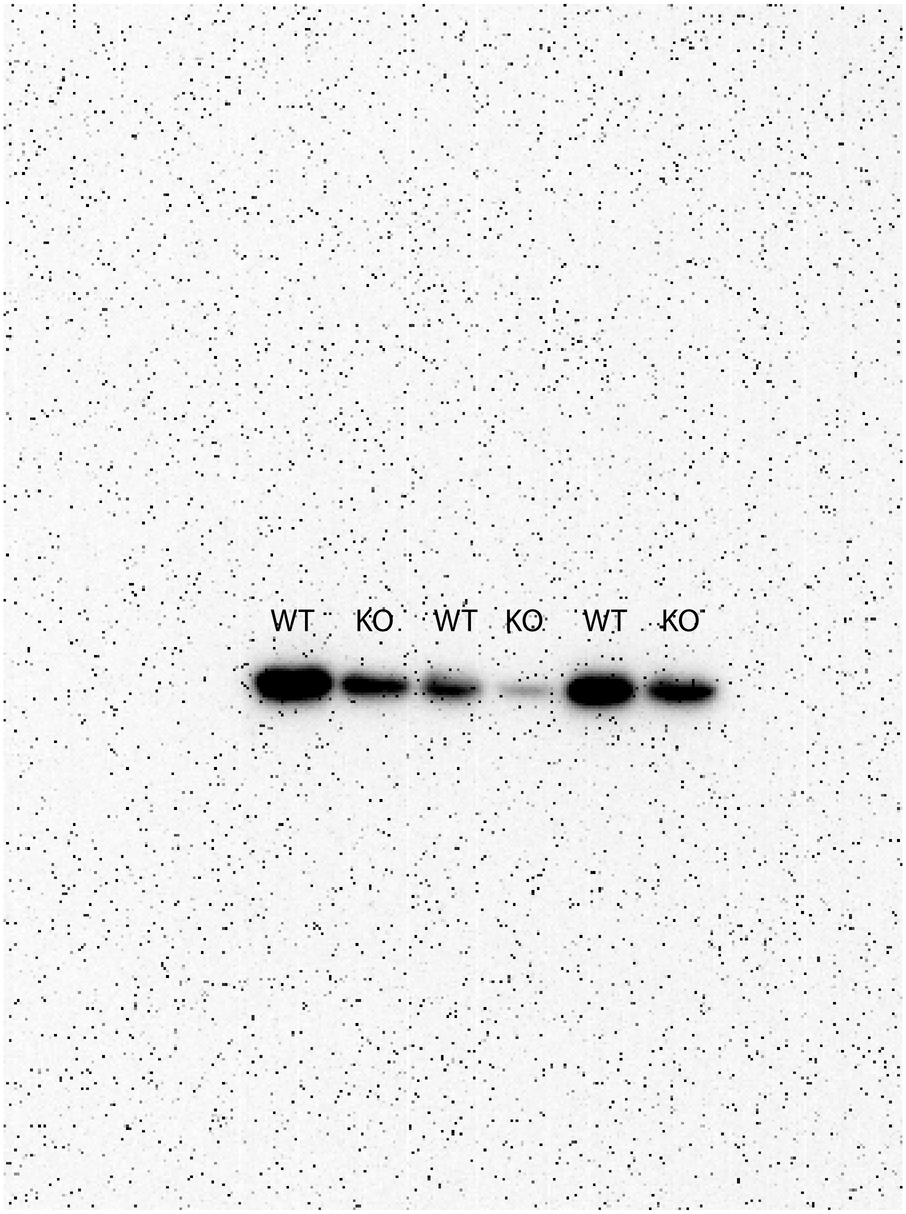

### Figure S1G

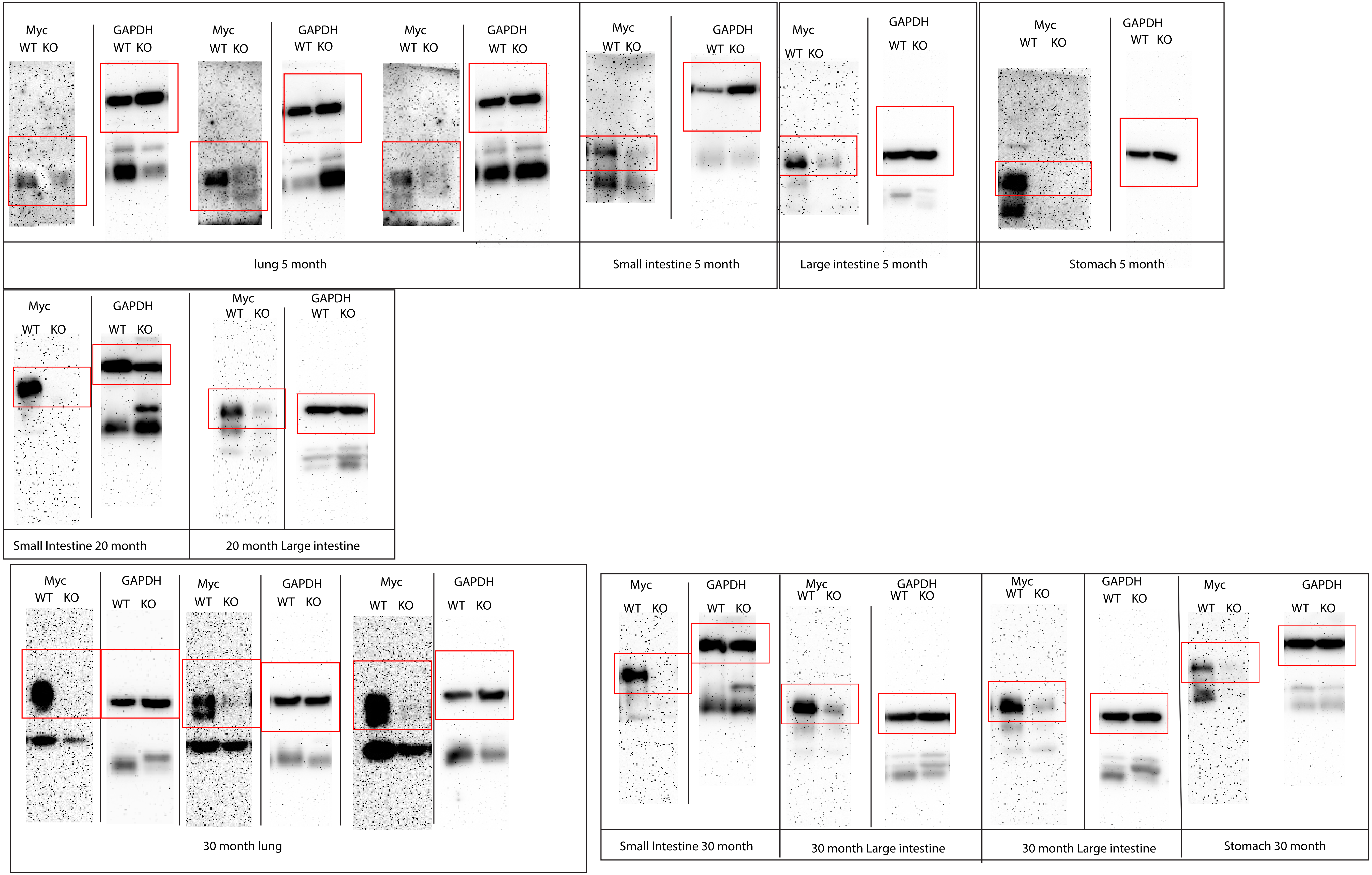

Myc
WT KO
GAPDH
WT KO
Myc
WT KO
GAPDH
WT KO
Myc
WT KO
GAPDH
WT KO
lung 5 month
Myc
WT KO
GAPDH
WT KO
Small intestine 5 month
Myc
WT KO
GAPDH
WT KO
Large intestine 5 month
Myc
WT KO
GAPDH
WT KO
Stomach 5 month
Myc
WT KO
GAPDH
WT KO
Small Intestine 20 month
Myc
WT KO
GAPDH
WT KO
20 month Large intestine
Myc
WT KO
GAPDH
WT KO
Myc
WT KO
GAPDH
WT KO
Myc
WT KO
GAPDH
WT KO
30 month lung
Myc
WT KO
GAPDH
WT KO
Small Intestine 30 month
Myc
WT KO
GAPDH
WT KO
30 month Large intestine
Myc
WT KO
GAPDH
WT KO
30 month Large intestine
Myc
WT KO
GAPDH
WT KO
Stomach 30 month
