## Supplementary material for "Premature Aging and Reduced Cancer Incidence Associated with Near-Complete Body-Wide *Myc* Inactivation": Suppl Figures S1-S7 and Table S3

Fig. S1

A

B

C

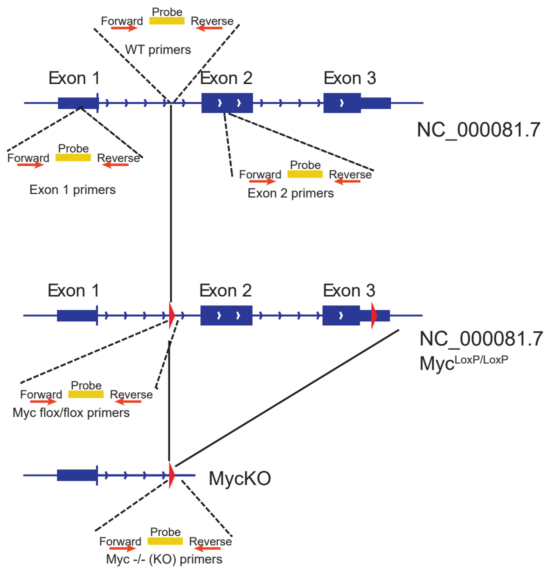


D

| | | Start (bp) | End (bp) | Sequence (5' -3') |
|---|---|---|---|---|
| WT (NC_000081.7) | Forward | 61858492 | 61858513 | GGGAATCCTCACATTCCTACTT |
| | Reverse | 61858643 | 61858626 | GATTCAGCACTGGGTGCA |
| | Probe | 61858549 | 61858572 | /56-FAM/TAGGAAGACTGCGGTGAGTCGTGA/3IABkFQ/ |
| $Myc^{loxP/loxP}$ (NC_000081.7) | Forward | 61858570 | 61858588 | TGATCTGAGCGGTTCCGTA |
| | Reverse | 61858697 | 61858675 | CACCTCCCTTCTACACTCTAAAC |
| | Probe | Loxp | Loxp | /56-FAM/CCTGCACGA/ZEN/TCCGGAACCCTTAAT/3IABkFQ/ |
| Myc -/- (KO) (NC_000081.7) | Forward | 61858570 | 61858588 | TGATCTGAGCGGTTCCGTA |
| | Reverse | 61858697 | 61858675 | TAAAGTCCCAAAGACACTCCAG |
| | Probe | Loxp | Loxp | /56-FAM/CCTGCACGATCCGGAACCCTTAAT/3IABkFQ/ |
| Exon 1 (NC_000081.7) | Forward | 61857404 | 61857425 | GCTGTAGTAATTCCAGCGAGAG |
| | Reverse | 61857507 | 61857488 | ACTCCAGAGCTGCCTTCTTA |
| | Probe | 61857447 | 61857468 | /56-FAM/TTGGAAGAG/ZEN/CCGTGTGTCCAGA/3IABkFQ/ |
| Exon 2 (NC_000081.7) | Forward | 61859421 | 61859442 | CTCCGTACAGCCCTATTTCATC |
| | Reverse | 61859542 | 61859524 | TGGGAAGCAGCTCGAATTT |
| | Probe | 61859464 | 61859480 | /56-FAM/TATCACCAG/ZEN/CAACAGCA |
| Cre (OM228709.1) | Forward | 3366 | 3344 | GCGGTCTGGCAGTAAAAACTATC |
| | Reverse | 3265 | 3287 | GTGAAACAGCATTGCTGTCACTT |
| | Probe | 3323 | 3300 | /56-FAM/AAACATGCT/ZEN/TCATCGTCGGTCCGG/3IABkFQ/ |
| Rosa WT (NC_000072.7) | Forward | 113053072 | 113053056 | CTGGCTTCTGAGGACCG |
| | Reverse | 113052875 | 113052894 | CCGAAAATCTGTGGGAAGTC |
| | Probe | 113052960 | 113052983 | /56-FAM/TTTAAGCCT/ZEN/GCCCAGAAGACTCCC/3IABkFQ/ |

E

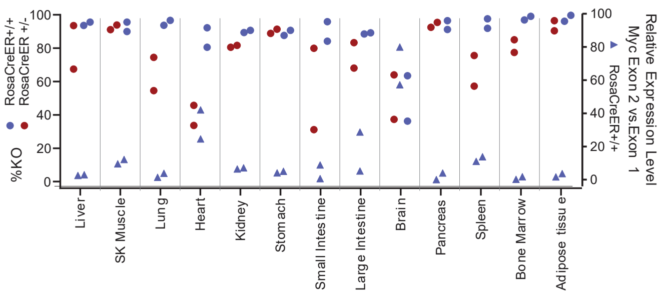


F

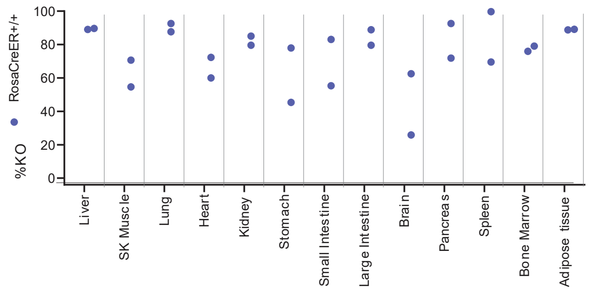


G

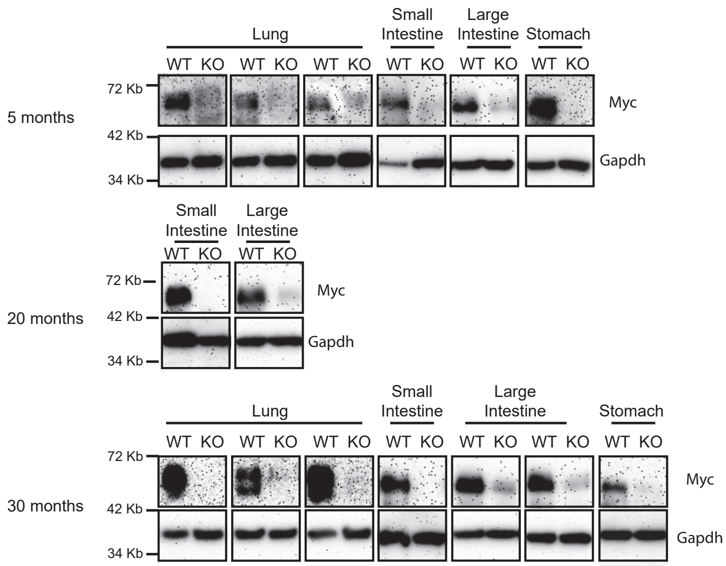


H

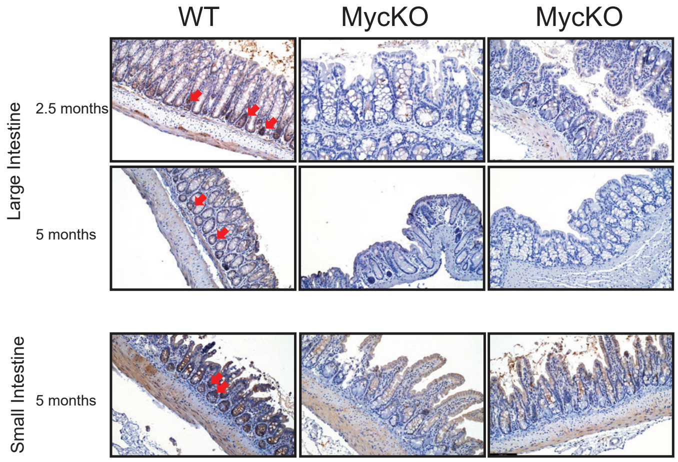

**Figure S1. Strategy for qPCR-based Taq-Man-based assay for identifying *Myc* alleles, related to STAR Methods.**

(A, B). The *Myc* loci of WT and $myc^{loxP/loxP}$ (B6.129S6-$Myc^{tm2Fwa}$/Mmjax) mice. LoxP sites are indicated by red triangles. In A, the red arrows indicate the primer sets used to amplify the WT allele. The reverse primer in intron 1 overlaps the inserted LoxP site in the ($Myc^{loxp/loxP}$) allele shown in B. Amplification with this primer set is thus possible only in WT mice and generates a PCR fragment detected with the unique TaqMan probe. In B, red arrows indicate the primer set flanking the 5' LoxP site that specifically amplifies this region and is detected with TaqMan probe.

(C). The excised and recombined *Myc* locus following CreER activation.

(D). Sequences and genomic locations for primers and probes shown in A-C as well as the qRT-PCR primers and probes used for qRT-PCR. Annotations for primers and probes are based on the numbering of the murine genomic sequence: https://www.ncbi.nlm.nih.gov/gene/17869.

(E). qPCR and qRT-PCR-based quantification of *Myc* locus excision and expression in ~6-7 wk old mice that had been treated for 5 days with tamoxifen at the time of weaning (~4 wks and weights >15 g). 2 mice with 2 copies of CreER and 2 with a single copy of CreER were used to determine how *Myc* locus excision efficiency was influenced by CreER copy number. After confirming the number of CreER alleles using the TaqMan-based approach described above, the WT:KO *Myc* allele ratio was determined from a standard curve of known amounts of each DNA (Wang et al., 2018; Wang et al., 2022b)(Wang et al., 2018; Wang et al., 2022b). qRT PCR reactions for Myc transcripts (https://www.ncbi.nlm.nih.gov/gene/17869 ) were performed on RNAs extracted from tissues of mice with two copies of ROSA-CreER as previously described

[1,2].

(F). Persistence of *Myc* locus deletion. qPCR performed as described in E on the indicted

tissues from 20-22 month-old *Myc*KO mice. See Table S1 for a summary of qPCR and qRT-PCR results performed on various tissues of additional mice.

(G). Immuno-blotting for Myc protein in the indicated tissues from 5-6 months, 20-22 months and 30-33 months old WT and *Myc*KO mice.

(H). Immunohistochemical staining for Myc protein in the small and large intestines of WT and *Myc*KO mice. Note, in both tissues, the absence of detectable Myc in the crypts which is where the highest levels of expression are confined. [3] Scale bar = 100 μm.

Fig. S2

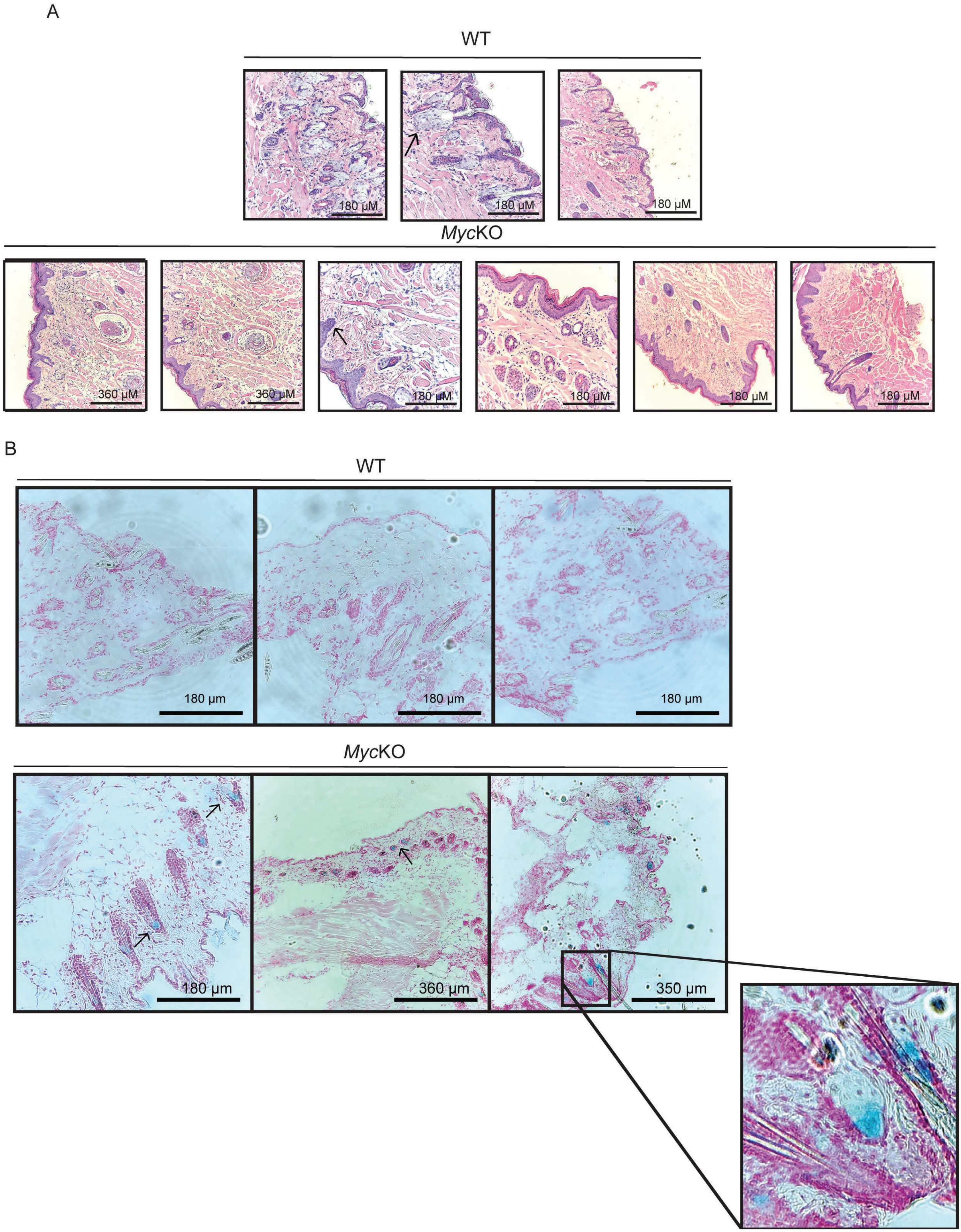
A
WT
180 μM
180 μM
180 μM
MycKO
360 μM
360 μM
180 μM
180 μM
180 μM
180 μM
B
WT
180 μm
180 μm
180 μm
MycKO
180 μm
360 μm
350 μm

**Figure S2. Histopathology of skin from alopecic regions of *Myc*KO mice and premature onset of senescence, related to Figure 1B&C.**

(A). H&E-stained sections of peri-orbital skin from a 5 mo *Myc*KO mouse and normal skin from the corresponding region of a WT control mouse. In WT panels, the arrow shows normal hair follicles and their adjacent sebaceous glands. In *Myc*KO panels, the arrows show thickening and hyperkeratinization of the epidermis, loss of surface invaginations and a generalized paucity of hair follicles and sebaceous glands. (B). SA-β-gal staining of the same areas from the WT and *Myc*KO mice shown in A. Arrows indicate SA-β-gal-positive cells in the latter samples adjacent to rare hair follicles. Most of SA-β-gal-positive cells detected resided within the inner and outer root sheath and base of the follicle (http://eulep.pdn.cam.ac.uk/~skinbase/). A higher power magnification of one of the fields containing SA-β-gal-positive cells is shown to the right.

Fig. S3

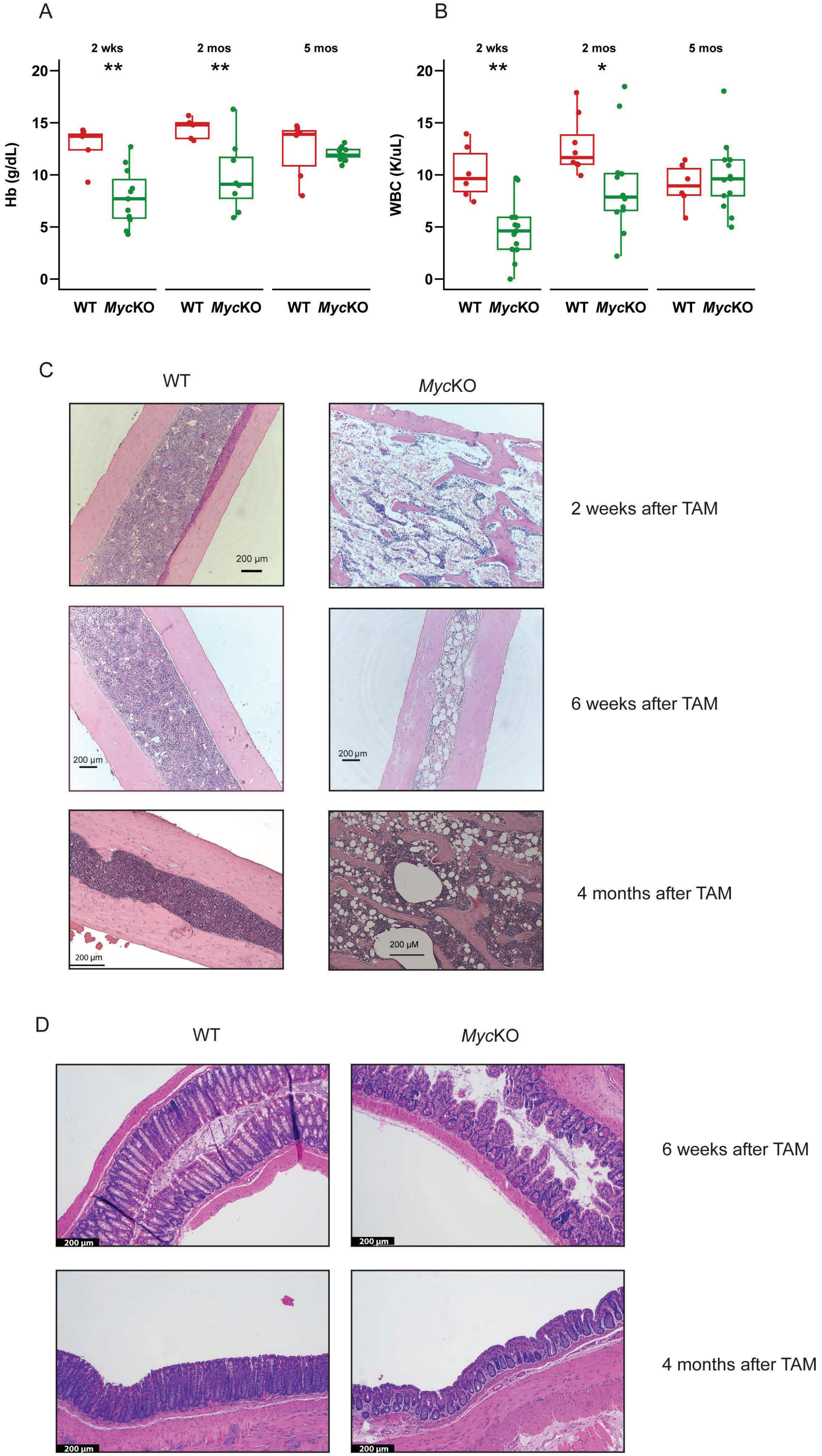
A
2 wks
2 mos
5 mos
**
**
Hb (g/dL)
0
5
10
15
20
WT MycKO
WT MycKO
WT MycKO
B
2 wks
2 mos
5 mos
**
*
WBC (K/uL)
0
5
10
15
20
WT MycKO
WT MycKO
WT MycKO
C
WT
MycKO
200 µm
2 weeks after TAM
200 µm
200 µm
6 weeks after TAM
200 µm
200 µM
4 months after TAM
D
WT
MycKO
200 µm
200 µm
6 weeks after TAM
200 µm
200 µm
4 months after TAM

**Figure S3. *Myc*KO mice develop transient mild-moderate, anemia, leukopenia and bone marrow hypoplasia, related to Result.**

(A). Hemoglobin levels in WT and age-matched *Myc*KO mice at the indicated ages, which are expressed relative to the time of starting 4-hydroxytamoxifen therapy. N = 6-14 at each age, Unpaired t test, ∗=$p < 0.05$, ∗∗=$p < 0.01$.

(B). Peripheral white blood cell counts performed at the times shown in A.

(C). H&E-stained bone marrows of WT and *Myc*KO mice of the indicated ages.

(D). H&E-stained colonic tissues from WT and *Myc*KO mice of the indicated ages.

Scale bar = 200 μm

Fig. S4

A

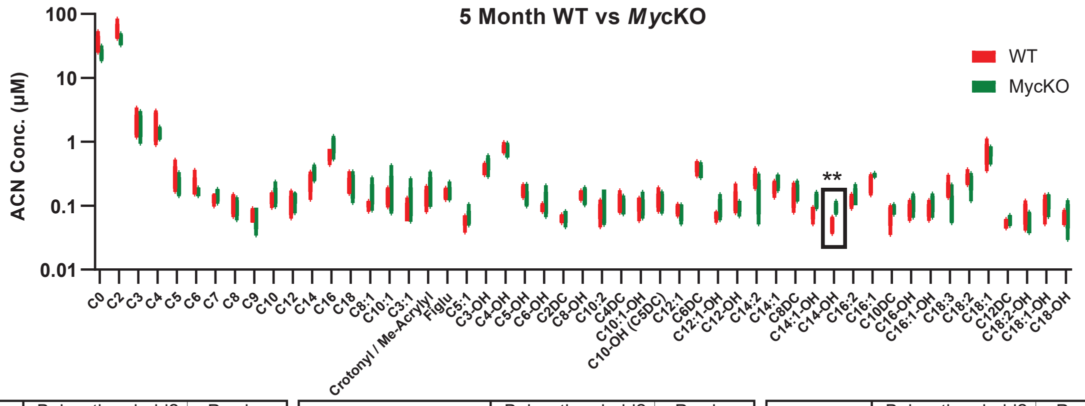


| | Below threshold? | P value |
|---|---|---|
| C0 | No | 0.082805 |
| C2 | No | 0.107371 |
| C3 | No | 0.891553 |
| C4 | No | 0.310665 |
| C5 | No | 0.243173 |
| C6 | No | 0.395319 |
| C7 | No | 0.342432 |
| C8 | No | 0.408316 |
| C9 | No | 0.932381 |
| C10 | No | 0.639306 |
| C12 | No | 0.252238 |
| C14 | No | 0.123141 |
| C16 | No | 0.112089 |
| C18 | No | 0.812097 |
| C8:1 | No | 0.179578 |
| C10:1 | No | 0.757984 |
| C3:1 | No | 0.365815 |

| | Below threshold? | P value |
|---|---|---|
| Crotonyl / Me-Acrylyl | No | 0.341981 |
| Figlu | No | 0.931718 |
| C5:1 | No | 0.726059 |
| C3-OH | No | 0.206181 |
| C4-OH | No | 0.735646 |
| C5-OH | No | 0.158956 |
| C6-OH | No | 0.359565 |
| C2DC | No | 0.437708 |
| C8-OH | No | 0.815773 |
| C10:2 | No | 0.357354 |
| C4DC | No | 0.519504 |
| C10:1-OH | No | 0.495535 |
| C10-OH (C5DC) | No | 0.804210 |
| C12:1 | No | 0.551986 |
| C6DC | No | 0.184811 |
| C12:1-OH | No | 0.258790 |
| C12-OH | No | 0.468176 |

| | Below threshold? | P value |
|---|---|---|
| C14:2 | No | 0.110939 |
| C14:1 | No | 0.174621 |
| C8DC | No | 0.640394 |
| C14:1-OH | No | 0.052795 |
| C14-OH | Yes ** | 0.003299 |
| C16:2 | No | 0.426658 |
| C16:1 | No | 0.066959 |
| C10DC | No | 0.267349 |
| C16-OH | No | 0.179856 |
| C16:1-OH | No | 0.179856 |
| C18:3 | No | 0.093283 |
| C18:2 | No | 0.605520 |
| C18:1 | No | 0.788717 |
| C12DC | No | 0.268356 |
| C18:2-OH | No | 0.243451 |
| C18:1-OH | No | 0.368712 |
| C18-OH | No | 0.548166 |

B

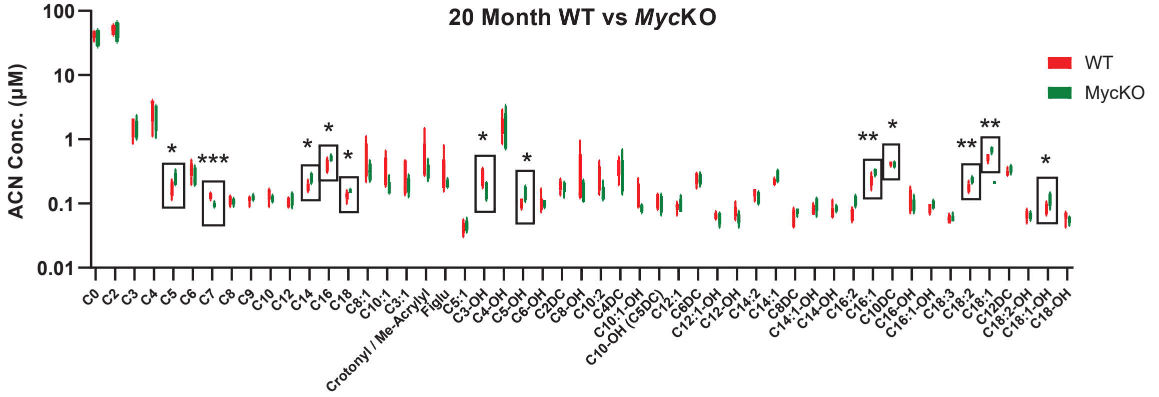


| | Below threshold? | P value |
|---|---|---|
| C0 | No | 0.301834 |
| C2 | No | 0.611193 |
| C3 | No | 0.856345 |
| C4 | No | 0.485919 |
| C5 | Yes * | 0.035634 |
| C6 | No | 0.260523 |
| C7 | Yes *** | 0.000783 |
| C8 | No | 0.497159 |
| C9 | No | 0.496558 |
| C10 | No | 0.222842 |
| C12 | No | 0.359745 |
| C14 | Yes * | 0.013240 |
| C16 | Yes * | 0.044914 |
| C18 | Yes * | 0.037790 |
| C8:1 | No | 0.319791 |
| C10:1 | No | 0.124682 |
| C3:1 | No | 0.442019 |

| | Below threshold? | P value |
|---|---|---|
| Crotonyl / Me-Acrylyl | No | 0.264035 |
| Figlu | No | 0.333312 |
| C5:1 | No | 0.498076 |
| C3-OH | Yes * | 0.016120 |
| C4-OH | No | 0.768323 |
| C5-OH | Yes * | 0.012020 |
| C6-OH | No | 0.996976 |
| C2DC | No | 0.720109 |
| C8-OH | No | 0.294558 |
| C10:2 | No | 0.137534 |
| C4DC | No | 0.413080 |
| C10:1-OH | No | 0.125231 |
| C10-OH (C5DC) | No | 0.590178 |
| C12:1 | No | 0.708297 |
| C6DC | No | 0.770883 |
| C12:1-OH | No | 0.590031 |
| C12-OH | No | 0.130382 |

| | Below threshold? | P value |
|---|---|---|
| C14:2 | No | 0.197887 |
| C14:1 | No | 0.073848 |
| C8DC | No | 0.096298 |
| C14:1-OH | No | 0.177224 |
| C14-OH | No | 0.335738 |
| C16:2 | Yes ** | 0.003900 |
| C16:1 | Yes * | 0.014118 |
| C10DC | No | 0.686387 |
| C16-OH | No | 0.293995 |
| C16:1-OH | No | 0.150042 |
| C18:3 | No | 0.969488 |
| C18:2 | Yes ** | 0.008329 |
| C18:1 | Yes ** | 0.003805 |
| C12DC | No | 0.117119 |
| C18:2-OH | No | 0.595252 |
| C18:1-OH | Yes * | 0.043729 |
| C18-OH | No | 0.225025 |

**Figure S4. Young and old *Myc*KO mice show evidence of Complex I defects and more generalized mitochondrial dysfunction, related to Figure 4F&G**

(A). 5 month-old mice. The results from Figure 4F are again shown along with the actual values obtained for each serum acyl carnitine. Unpaired t test, $**$=$p < 0.01$

(B). 20 month-old mice. The results from Figure 4G are again shown along with the actual values obtained for each serum acyl carnitine. Unpaired t test, $*$=$p < 0.05$, $**$=$p < 0.01$, $***$=$p < 0.001$.

Fig. S5

A

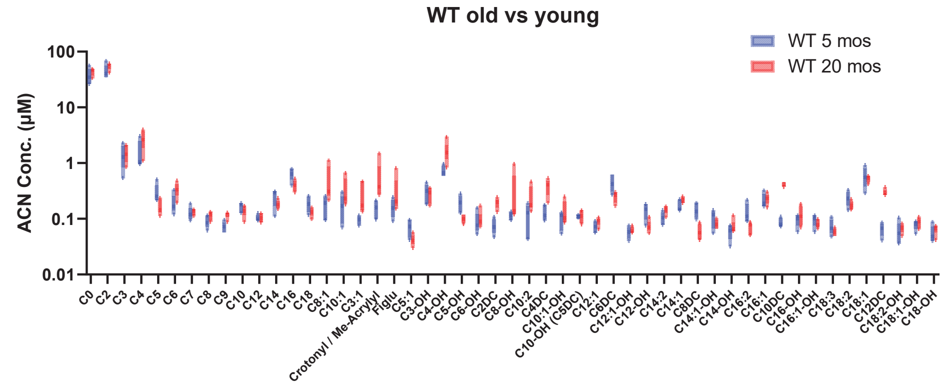


| | Below threshold? | P value |
|---|---|---|
| C0 | No | 0.265224 |
| C2 | No | 0.190087 |
| C3 | No | 0.774480 |
| C4 | No | 0.102879 |
| C5 | Yes * | 0.036385 |
| C6 | Yes * | 0.027669 |
| C7 | No | 0.693571 |
| C8 | No | 0.097887 |
| C9 | Yes *** | 0.000949 |
| C10 | No | 0.214036 |
| C12 | No | 0.969853 |
| C14 | No | 0.099644 |
| C16 | Yes ** | 0.009118 |
| C18 | No | 0.055543 |
| C8:1 | No | 0.083082 |
| C10:1 | No | 0.124479 |
| C3:1 | No | 0.050142 |

| | Below threshold? | P value |
|---|---|---|
| Crotonyl / Me-Acrylyl | No | 0.080009 |
| Figlu | No | 0.202828 |
| C5:1 | Yes * | 0.023385 |
| C3-OH | No | 0.708989 |
| C4-OH | Yes * | 0.017946 |
| C5-OH | Yes ** | 0.006523 |
| C6-OH | No | 0.612026 |
| C2DC | Yes *** | 0.000196 |
| C8-OH | No | 0.202814 |
| C10:2 | No | 0.067778 |
| C4DC | Yes ** | 0.001957 |
| C10:1-OH | No | 0.274583 |
| C10-OH (C5DC) | No | 0.879301 |
| C12:1 | No | 0.084011 |
| C6DC | Yes * | 0.014555 |
| C12:1-OH | No | 0.372662 |
| C12-OH | Yes ** | 0.007948 |

| | Below threshold? | P value |
|---|---|---|
| C14:2 | Yes * | 0.024092 |
| C14:1 | Yes * | 0.035823 |
| C8DC | Yes ** | 0.001306 |
| C14:1-OH | No | 0.173232 |
| C14-OH | No | 0.233375 |
| C16:2 | Yes * | 0.024105 |
| C16:1 | No | 0.718401 |
| C10DC | Yes **** | <0.000001 |
| C16-OH | No | 0.328667 |
| C16:1-OH | No | 0.242023 |
| C18:3 | No | 0.137024 |
| C18:2 | No | 0.099872 |
| C18:1 | No | 0.583925 |
| C12DC | Yes **** | <0.000001 |
| C18:2-OH | No | 0.493457 |
| C18:1-OH | No | 0.419368 |
| C18-OH | No | 0.325839 |

B

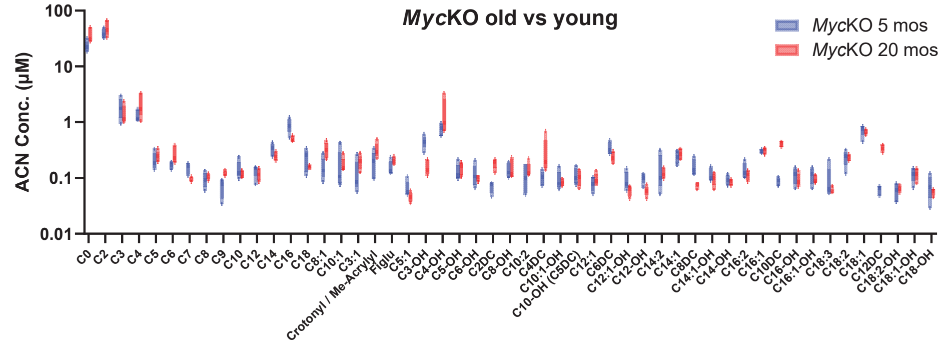


| | Below threshold? | P value |
|---|---|---|
| C0 | Yes * | 0.030896 |
| C2 | No | 0.184339 |
| C3 | No | 0.361412 |
| C4 | No | 0.107424 |
| C5 | No | 0.425804 |
| C6 | Yes * | 0.041003 |
| C7 | Yes ** | 0.005872 |
| C8 | No | 0.415367 |
| C9 | Yes ** | 0.003738 |
| C10 | No | 0.355998 |
| C12 | No | 0.370061 |
| C14 | No | 0.083011 |
| C16 | Yes * | 0.011845 |
| C18 | No | 0.150392 |
| C8:1 | Yes * | 0.018524 |
| C10:1 | No | 0.958586 |
| C3:1 | No | 0.135059 |

| | Below threshold? | P value |
|---|---|---|
| Crotonyl / Me-Acrylyl | No | 0.084518 |
| Figlu | No | 0.092821 |
| C5:1 | No | 0.129585 |
| C3-OH | Yes *** | 0.000969 |
| C4-OH | No | 0.177247 |
| C5-OH | No | 0.446142 |
| C6-OH | No | 0.554716 |
| C2DC | Yes *** | 0.000204 |
| C8-OH | No | 0.742848 |
| C10:2 | No | 0.419328 |
| C4DC | No | 0.111577 |
| C10:1-OH | No | 0.622889 |
| C10-OH (C5DC) | No | 0.594450 |
| C12:1 | No | 0.389429 |
| C6DC | Yes * | 0.048989 |
| C12:1-OH | No | 0.089490 |
| C12-OH | Yes ** | 0.005306 |

| | Below threshold? | P value |
|---|---|---|
| C14:2 | No | 0.590118 |
| C14:1 | No | 0.502287 |
| C8DC | Yes ** | 0.009042 |
| C14:1-OH | No | 0.281853 |
| C14-OH | No | 0.377949 |
| C16:2 | No | 0.263590 |
| C16:1 | No | 0.453775 |
| C10DC | Yes **** | <0.000001 |
| C16-OH | No | 0.345020 |
| C16:1-OH | No | 0.325627 |
| C18:3 | No | 0.302453 |
| C18:2 | No | 0.995663 |
| C18:1 | No | 0.610594 |
| C12DC | Yes **** | <0.000001 |
| C18:2-OH | No | 0.521089 |
| C18:1-OH | No | 0.967776 |
| C18-OH | No | 0.311576 |

**Figure S5. Both young and old WT and *Myc*KO mice show evidence of age-related differences in serum acyl carnitine levels, related to Figure 4F&G**

(A). 5 month old and 20 month-old WT mice. Unpaired t test, ∗=$p < 0.05$, ∗∗=$p < 0.01$, ∗∗∗=$p < 0.001$, ∗∗∗∗=$p < 0.0001$.

(B). 5 month old and 20 month-old *Myc*KO mice. Unpaired t test, ∗=$p < 0.05$, ∗∗=$p < 0.01$, ∗∗∗=$p < 0.001$, ∗∗∗∗=$p < 0.0001$.

Fig. S6

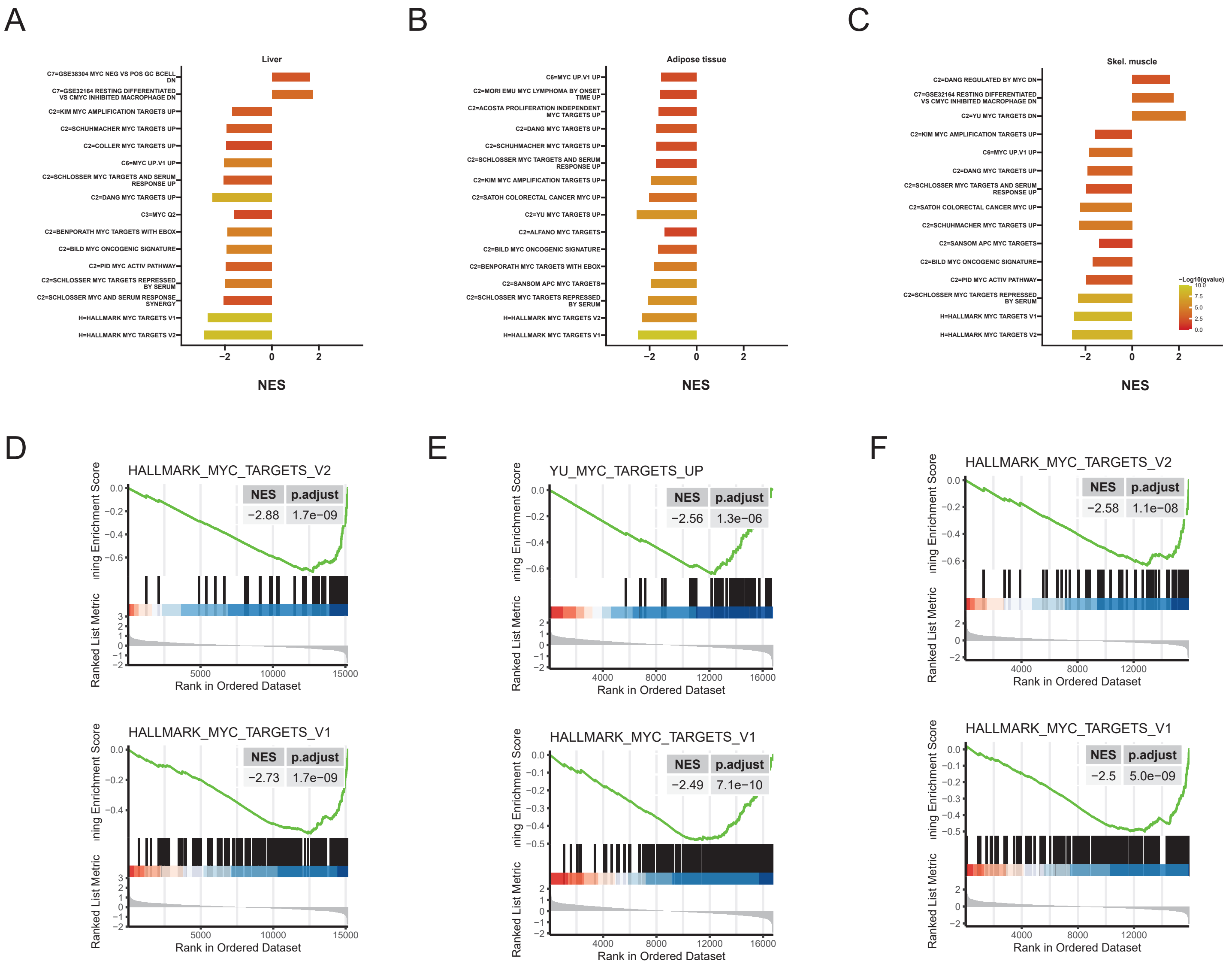
A
Liver
C7=GSE38304 MYC NEG VS POS GC BCELL DN
C7=GSE32164 RESTING DIFFERENTIATED VS CMYC INHIBITED MACROPHAGE DN
C2=KIM MYC AMPLIFICATION TARGETS UP
C2=SCHUHMACHER MYC TARGETS UP
C2=COLLER MYC TARGETS UP
C6=MYC UP.V1 UP
C2=SCHLOSSER MYC TARGETS AND SERUM RESPONSE UP
C2=DANG MYC TARGETS UP
C3=MYC Q2
C2=BENPORATH MYC TARGETS WITH EBOX
C2=BILD MYC ONCOGENIC SIGNATURE
C2=PID MYC ACTIV PATHWAY
C2=SCHLOSSER MYC TARGETS REPRESSED BY SERUM
C2=SCHLOSSER MYC AND SERUM RESPONSE SYNERGY
H=HALLMARK MYC TARGETS V1
H=HALLMARK MYC TARGETS V2
NES
B
Adipose tissue
C6=MYC UP.V1 UP
C2=MORI EMU MYC LYMPHOMA BY ONSET TIME UP
C2=ACOSTA PROLIFERATION INDEPENDENT MYC TARGETS UP
C2=DANG MYC TARGETS UP
C2=SCHUHMACHER MYC TARGETS UP
C2=SCHLOSSER MYC TARGETS AND SERUM RESPONSE UP
C2=KIM MYC AMPLIFICATION TARGETS UP
C2=SATOH COLORECTAL CANCER MYC UP
C2=YU MYC TARGETS UP
C2=ALFANO MYC TARGETS
C2=BILD MYC ONCOGENIC SIGNATURE
C2=BENPORATH MYC TARGETS WITH EBOX
C2=SANSOM APC MYC TARGETS
C2=SCHLOSSER MYC TARGETS REPRESSED BY SERUM
H=HALLMARK MYC TARGETS V2
H=HALLMARK MYC TARGETS V1
NES
C
Skel. muscle
C2=DANG REGULATED BY MYC DN
C7=GSE32164 RESTING DIFFERENTIATED VS CMYC INHIBITED MACROPHAGE DN
C2=YU MYC TARGETS DN
C2=KIM MYC AMPLIFICATION TARGETS UP
C6=MYC UP.V1 UP
C2=DANG MYC TARGETS UP
C2=SCHLOSSER MYC TARGETS AND SERUM RESPONSE UP
C2=SATOH COLORECTAL CANCER MYC UP
C2=SCHUHMACHER MYC TARGETS UP
C2=SANSOM APC MYC TARGETS
C2=BILD MYC ONCOGENIC SIGNATURE
C2=PID MYC ACTIV PATHWAY
C2=SCHLOSSER MYC TARGETS REPRESSED BY SERUM
H=HALLMARK MYC TARGETS V1
H=HALLMARK MYC TARGETS V2
−Log10(qvalue)
NES
D
HALLMARK_MYC_TARGETS_V2
NES p.adjust
−2.88 1.7e−09
HALLMARK_MYC_TARGETS_V1
NES p.adjust
−2.73 1.7e−09
E
YU_MYC_TARGETS_UP
NES p.adjust
−2.56 1.3e−06
HALLMARK_MYC_TARGETS_V1
NES p.adjust
−2.49 7.1e−10
F
HALLMARK_MYC_TARGETS_V2
NES p.adjust
−2.58 1.1e−08
HALLMARK_MYC_TARGETS_V1
NES p.adjust
−2.5 5.0e−09
Ranked List Metric
Rank in Ordered Dataset

**Figure S6. Dysregulation of Myc target gene sets in 5 mo MycKO tissues, related to Results.**

(A-C). GSEA summaries of positively-regulated direct Myc targets in liver, adipose tissue, and skeletal muscle, respectively from 5 month-old MycKO mice. All Myc targets from the MSigDB Collection (http://www.gsea-msigdb.org/gsea/msigdb/collections.jsp) were analyzed by GSEA to determine the overall directionality of expression of each set's component transcripts.

(D-F). Select examples of individual GSEA profiles from A-C.

Fig. S7

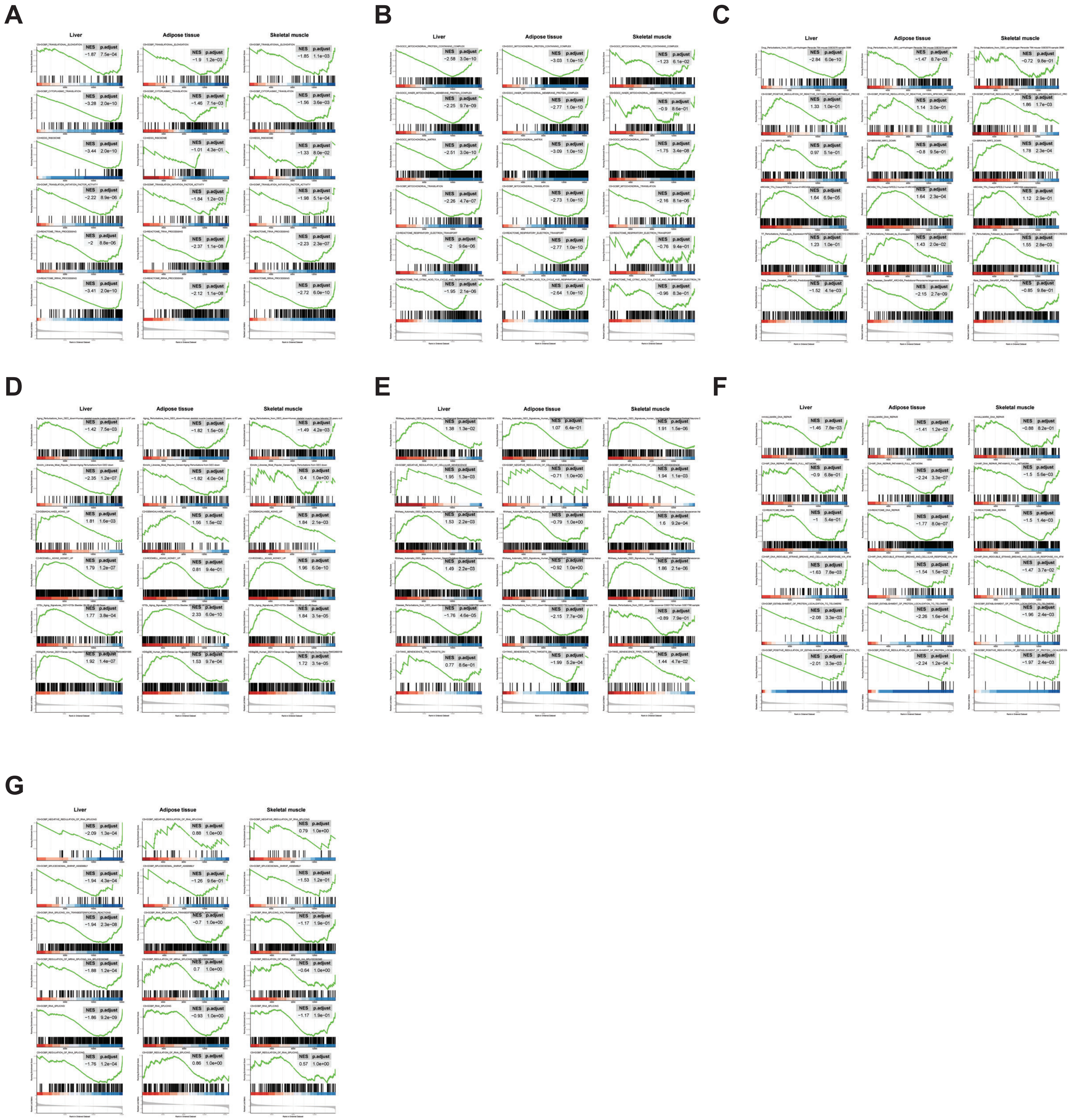
A
Liver
Adipose tissue
Skeletal muscle
B
Liver
Adipose tissue
Skeletal muscle
C
Liver
Adipose tissue
Skeletal muscle
D
Liver
Adipose tissue
Skeletal muscle
E
Liver
Adipose tissue
Skeletal muscle
F
Liver
Adipose tissue
Skeletal muscle
G
Liver
Adipose tissue
Skeletal muscle

**Figure S7. GSEA profiles in liver, adipose tissue and skeletal muscle from 5 month-old WT and MycKO mice, related to Figure 6A.** Normalized enrichment scores and q values are indicated in the upper right corner of each profile. Data used to generate the ridge plots shown in Figure 6A are re-graphed and included here**.**

(A) GSEA profiles for “Translation/ribosomal structure and function”.
(B) GSEA profiles for “Mitochondrial structure and function”.
(C) GSEA profiles for “Oxidative stress response”.
(D) GSEA profiles for “Aging”.
(E) GSEA profiles for “Senescence”.
(F) GSEA profiles for “DNA damage response and repair”.
(G) GSEA profiles for “mRNA splicing”.

**Table S3.** Antibodies used for the current studies, related to STAR Methods.

| Name of antigen target | Type of antibody | Vendor | Catalog number | dilution | Purpose |
|---|---|---|---|---|---|
| **Glut 1** | Rabbit mAb | Abcam | Ab115730 | 1:2,000 | WB |
| **Glut 2** | Rabbit pAb* | Proteintech | 20436-1-AP | 1:1,000 | WB |
| **Glut 4** | Mouse mAb | Cell signaling | 2213 | 1:1,000 | WB |
| **GAPDH** | Mouse mAb | Sigma | G8795 | 1:10,000 | WB |
| **γH2A.X** | Rabbit mAb | Abcam | ab81299 | 1:200 | IHF |
| **PDH** | Rabbit mAb | CST | 3205 | 1:1,000 | WB |
| **c-Myc** | Rabbit mAb | Cell signaling | 13987 | 1:1,000<br>1:400 | WB |
| **c-Myc(N-262)** | Rabbit pAb | Santa Cruze | sc-764 | 1:250 | IHC |
| **p-PDH** | Rabbit pAb | Cal biochem | AP1062 | 1:300 | WB |
| **PFK-L** | Rabbit pAb | Avivva sys bio | 855570 | 1:400 | WB |
| **PFK-M** | Mouse mAb | R+D systems | MAB7687 | 1:3,000 | WB |
| **IgG** | HRP-Goat-anti-rabbit | Cell signaling | 7074 | 1:5,000<br>1:2,000 | Secondary for WB |
| **IgG** | HRP-Goat-anti-mouse | Cell signaling | 7076 | 1:10,000 | Secondary for WB |

*pAb=Polyclonal antibody
